# PhiPipe: a multi-modal MRI data processing pipeline with test-retest reliability and predicative validity assessments

**DOI:** 10.1101/2022.06.22.497141

**Authors:** Yang Hu, Qingfeng Li, Kaini Qiao, Xiaochen Zhang, Bing Chen, Zhi Yang

**Affiliations:** Laboratory of Psychological Health and Imaging, Shanghai Mental Health Center, Shanghai Jiao Tong University School of Medicine, Shanghai, China; Shanghai Key Laboratory of Psychotic Disorders, Shanghai Mental Health Center, Shanghai Jiao Tong University School of Medicine, Shanghai, China; Institute of Psychological and Behavioral Sciences, Shanghai Jiao Tong University, Shanghai, China; Brain Science and Technology Research Center, Shanghai Jiao Tong University, Shanghai, China; Beijing University of Posts and Telecommunications, Beijing, China; Jing Hengyi School of Education, Hangzhou Normal University, Zhejiang, China

## Abstract

Magnetic Resonance Imaging (MRI) has been one of the primary instruments to measure the properties of the human brain non-invasively in vivo. MRI data generally needs to go through a series of processing steps (i.e., a pipeline) before statistical analysis. Currently, the processing pipelines for multi-modal MRI data are still rare, in contrast to single-modal pipelines. Furthermore, the reliability and validity of the output of the pipelines are critical for the MRI studies. However, the reliability and validity measures are not available or adequate for almost all pipelines.

Here, we present PhiPipe, a multi-modal MRI processing pipeline. PhiPipe could process T1-weighted, resting-state BOLD, and diffusion-weighted MRI data and generate commonly used brain features in neuroimaging. We evaluated the test-retest reliability of PhiPipe’s brain features by computing intra-class correlations (ICC) in four public datasets with repeated scans. We further evaluated the predictive validity by computing the correlation of brain features with chronological age in three public adult lifespan datasets. The multi-variate reliability and predictive validity of the PhiPipe results were also evaluated. The results of PhiPipe were consistent with previous studies, showing comparable or better reliability and validity when compared with two popular single-modality pipelines, namely DPARSF and PANDA.

The publicly available PhiPipe provides a simple-to-use solution to multi-modal MRI data processing. The accompanied reliability and validity assessments could help researchers make informed choices in experimental design and statistical analysis. Furthermore, this study provides a framework for evaluating the reliability and validity of image processing pipelines.

## 1. Introduction

Magnetic Resonance Imaging (MRI) has been the standard measurement tool in the current field of human brain research. The popularity of MRI comes from its non-invasiveness, flexible image contrasts and whole brain coverage, compared with other brain imaging tools. The MRI has been expected to play an important role in elucidating the brain-behavior relationship and aiding diagnosis and treatment of mental disorders in the years to come.

The MRI data normally requires a series of processing steps before statistical modelling and hypothesis testing, compared with other types of data (such as neuropsychological tests). This is because MRI data is an indirect measure of brain properties and algorithms must be applied to extract features of interest from MRI images. In addition, the MRI data is corrupted by noises and artefacts from various sources such as head motion and field inhomogeneity, which should be mitigated during the data processing. The processing steps taken from the raw MRI data to the final MRI-based brain features before statistical analysis are usually called a processing pipeline. From a conceptual level, we define the software (i.e., algorithm implementation) for individual processing step as the atomic software. Thus, image processing pipelines are made of a combination of atomic softwares. FreeSurfer (Dale et al., 1999; Fischl et al., 1999), AFNI (Cox, 1996; Taylor and Saad, 2013), FSL (Smith et al., 2004) and SPM (Ashburner, 2012) contain a lot of atomic softwares for MRI data processing, on which most existing pipelines would depend. These core softwares also have developed their own pipelines to facilitate batch processing, for instance, FreeSurfer’s recon-all, AFNI’s afni_proc.py, FSL’s Feat and SPM’s batch system. The publicly available pipelines, such as DPARSF (Yan and Zang, 2010), CONN (Whitfield-Gabrieli and Nieto-Castanon, 2012), HCP-Pipelines (Glasser et al., 2013), PANDA (Cui et al., 2013), C-PAC (Craddock et al., 2013), GRETNA (Wang et al., 2015), CCS (Xu et al., 2015), fMRIPrep (Esteban et al., 2019) and CAT12 (Gaser et al., 2022), have remarkably simplified the data processing procedures and contributed to the standardization of image processing steps, making the results from multiple studies more comparable and making multi-center studies possible (See Supplementary Table S1 for a longer list of pipelines we investigated).

However, while collecting multi-modal imaging data in one scan session is a common practice, the processing pipelines for multi-modal MRI data are still rare, in comparison with single-modal pipelines. In other words, multi-modal data are collected but processed separately. The downside of using multiple single-modal pipelines is to make multi-modal data fusion difficult. This is because single-modal pipelines usually use different templates and atlases due to their developers’ preferences. Moreover, using multiple single-modal pipelines usually result in redundant calculations.

The major challenge in designing a (multi-modal) pipeline is how to combine different atomic softwares to achieve optimal performance. However, currently there is no consensus on the criterion of optimum. A variety of metrics were used in previous pipeline validations, to just name a few, spatial smoothness (Esteban et al., 2019), consistency within and between datasets (Cruces et al., 2022), between-group difference detection (Cui et al., 2013; Xu et al., 2018), discriminability (Lawrence et al., 2021), inter-pipeline agreement (Li et al., 2021), and age predication/correlation (Tustison et al., 2014; Yan et al., 2016; Alfaro-Almagro et al., 2018). However, these previous validations were usually not systematic by considering only one brain feature in one dataset or the metrics used were not informative for end-users in experiment design and statistical analysis.

Here we propose that, for the brain features generated by the pipeline, the test-retest reliability measured by intra-class correlation (ICC) and predicative validity measured by correlation with chronological age should be taken as the two of the standard metrics in pipeline validation. The brain features instead of the intermediate results are of most interest and, therefore, their reliability and validity are most informative and directly related to the experimental design and statistical analysis. The test-retest reliability of various brain features has been examined extensively (Chen et al., 2015; Buimer et al., 2020). However, the reliability analysis is dependent on the specific pipeline used. Previous studies of test-retest reliability in almost all cases considered one pipeline only, the results of which could not be generalized to all pipelines. In addition, there were other types of reliability measures besides ICC, for instance, Pearson correlation coefficient (Iscan et al., 2015), percent difference (Iscan et al., 2015), coefficient of variation (Owen et al., 2013), Kendall coefficient of concordance (Patriat et al., 2013) and discriminability (Bridgeford et al., 2021) used in previous studies and we choose to use ICC by taking three aspects into consideration: (1) the ICC approaches reliability from the perspective of inter-subject variability, which is often of great interest in this field (Chen et al., 2018); (2) the ICC has a direct relationship with the predictive validity to be examined (Elliott et al., 2020); (3) the ICC is the most widely used measure in previous studies, which could be used to validate the results in the current study.

Predictive validity is much more difficult to test, as there is no ground truth. We suggest that the relationship between age and brain features during the adult lifespan could represent the predictive validity of the brain features. The reasons are three folds: (1) it is obvious that our brain changes with age from the early to the late adulthood and it is safe to assume that there is some relationship between any brain feature and age during the adult lifespan. (2) The age effect is large, wide-spread and reproducible, as demonstrated by previous studies based on more than thousands of subjects (Masouleh et al., 2019; Frangou et al., 2022). In other words, there is a proxy of ground truth. In contrast, the associations between brain features and cognitive functions or personalities are much smaller and hardly replicated (Masouleh et al., 2019; Marek et al., 2022). (3) The age information is most easily accessible in any public datasets and thus the cost of validation and replication is relatively small.

In this study, we first present a multi-modal pipeline, PhiPipe. PhiPipe can process T1-weighted, resting-state BOLD, and diffusion-weighted (DWI) MRI images and generate thirteen commonly-used brain features characterizing the structural and functional properties of brain. We then evaluated the test-retest reliability in four public datasets and the predicative validity in three public datasets for the brain features. We finally compared the reliability and validity results of PhiPipe with those of two other single-modal pipelines, namely, DPARSF (Yan and Zang, 2010) and PANDA (Cui et al., 2013), using the same datasets.

## 2. Methods

### 2.1 The design and implementation of PhiPipe

The PhiPipe (v1.2.0) mainly relies on FreeSurfer (v6.0.0) for T1 processing, AFNI (v20.0.19) for resting-state BOLD fMRI processing, and FSL (v6.0.2) for DWI processing. Different from some pipelines, the PhiPipe is intentionally designed to use one of the three softwares for a single modality to reduce the risks of incompatibility and make the maintenance easier when the dependent softwares are updated. The PhiPipe mainly consists of a set of Bash scripts, which is the same way we usually interact with FreeSurfer, AFNI and FSL. Therefore, anyone who has some experience with FreeSurfer, AFNI or FSL could use PhiPipe easily. Furthermore, the PhiPipe is designed to generate atlas-based (instead of voxel-based or vertex-based) brain features so that downstream statistical analysis could be easier. The PhiPipe is publicly available at https://github.com/phi-group/PhiPipe-release.

Figure 1 shows the flowchart of PhiPipe. The processing steps of T1-weighted images include:

1. Use FreeSurfer’s *recon-all* to perform skull stripping, tissue segmentation, surface reconstruction, anatomical parcellation, etc. Optionally, the skull stripping could be done using CAT12 (Gaser et al., 2022) instead of FreeSurfer. The skull-stripping step to some degree determines the reconstruction quality of pial surface (the surface between gray matter and external CSF), which further has an impact on the estimated brain features like cortical thickness. The FreeSurfer’s skull stripping sometimes could not achieve satisfactory results, so we include the CAT12’s skull stripping as an alternative method. There are two more reasons to introduce CAT12 into the T1 processing of PhiPipe. CAT12 provides an image quality rating score (http://www.neuro.uni-jena.de/cat12-html/cat_methods_QA.html), which could be used to check the raw data quality quantitatively. The rating score has been shown to be highly predicative of manual inspection results (Gilmore et al., 2021). In addition, CAT12 provides an estimate of total intracranial volume (TIV) based on tissue segmentation. TIV is an important confounding factor in structural analysis (Malone et al., 2015). The TIV estimation in FreeSurfer is based on affine registration into a template and biased (Klasson et al., 2018), the accuracy of which may be further degraded when defacing was applied as indicated in FreeSurfer’s release notes (https://surfer.nmr.mgh.harvard.edu/fswiki/ReleaseNotes). Normally, public datasets are defaced to protect participants’ privacy. Therefore, we would like to test whether the TIV estimated by CAT12 could be better than the one by FreeSurfer in real datasets.
2. Parcellate the cortical surface using Schaefer Atlas (Schaefer et al., 2018). The Schaefer Atlas is a cortical parcellation atlas based on resting-state functional connectivity. It divides the cerebral cortex into 100-1000 parcels and in PhiPipe we choose the 100-parcel version to increase the signal-to-noise ratio of each parcel. One advantage of the Schaefer Atlas is that the 100 parcels were assigned into the widely-used Yeo’s seven networks (Yeo et al., 2011), which could be convenient for network analysis. Besides the Schaefer Atlas, the FreeSurfer’s built-in Desikan-Killiany (DK) Atlas (Desikan et al., 2006) and Aseg Atlas (Fischl et al., 2002) are used in PhiPipe.
3. Create masks for whole brain, gray matter, white matter, ventricle, and cortical/subcortical parcellations based on the results above. These masks are used for BOLD and DWI data processing.
4. Extract cortical thickness (CT), cortical area (CA), cortical volume (CV) measures (Fischl and Dale, 2000; Winkler et al., 2018) for the DK Atlas and Schaefer Atlas, and subcortical volume (SV) measures for the Aseg Atlas. Only the seven commonly used bilateral subcortical structures (i.e., thalamus, caudate, putamen, pallidum, hippocampus, amygdala and accumbens) of the Aseg Atlas are included.
5. Nonlinear registration between T1 brain image and MNI152 T1 brain template (also known as the 6^th^ generation non-linear asymmetric version) is conducted using ANTs (v2.2.0, Tustison et al., 2014). The MNI152 T1 brain template is supplied by FSL. The registration results are used in the BOLD processing.
6. Quality control pictures for brain extraction, tissue segmentation, parcellation and registration are created to visually check the processing results. An example of these quality control pictures could be found in Supplementary Figure S1.

**Figure 1.**
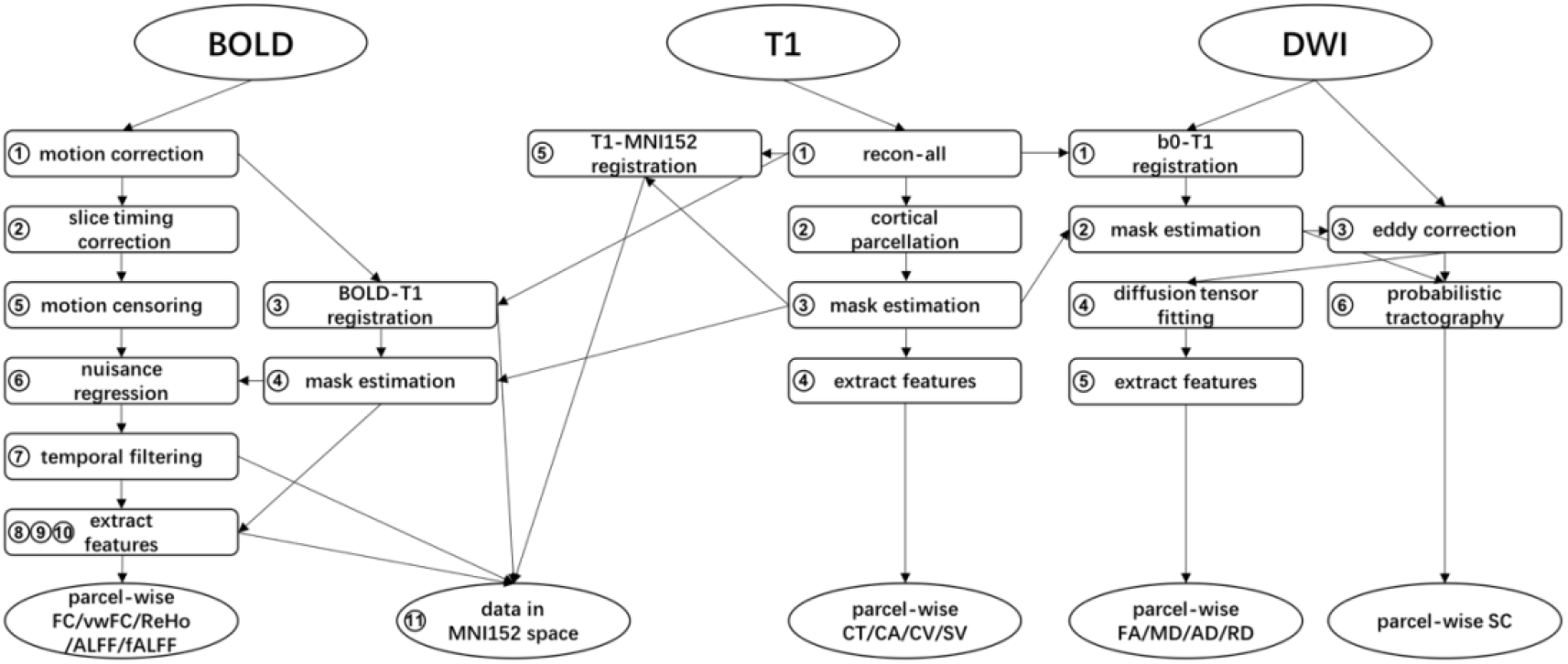
Flow chart of processing steps in PhiPipe. The ellipses represent raw MRI data and the processed brain features. The rectangles represent individual processing steps. The circled numbers correspond to the order of processing steps for each modality.

The processing steps of resting-state BOLD images include:

1. Correct head motion using AFNI’s *3dvolreg* and create measures to quantify head motion. Power’s framewise displacement (FD) is calculated to reflect volume-wise head movement (Power et al., 2012). Motion outliers (volumes with large motion) are detected at a threshold of FD=0.5mm. Mean FD and outlier ratio (the ratio between motion outliers and total volumes) are used as quantitative measures of overall head motion. By default, the first 5 volumes are removed before motion correction to allow for magnetization equilibrium.
2. Correct slice acquisition timing using AFNI’s *3dTshift*. Alternatively, this step could be omitted as previous studies have shown that the impact of slice timing correction is very small in resting-state fMRI (Wu et al., 2011; Shirer et al., 2015).
3. Boundary-based registration between median volume and T1 image is conducted using FreeSurfer’s *bbregister* (Greve and Fischl, 2009).
4. Create masks for whole brain, gray matter, white matter, ventricle, and parcellations based on T1 processing and BOLD-T1 registration results.
5. Interpolate the motion outliers by neighboring volumes (i.e., motion censoring).
6. Regress out nuisance signals, in which mean white matter and ventricle signals, and Friston’s 24-parameter head motion model are included. The linear and quadratic trends are also removed. Alternatively, the global signal (mean signal of the whole brain) could be added in the nuisance regression, as global signal regression is a controversial processing step (Murphy and Fox, 2017).
7. Bandpass filtering at 0.01-0.1 Hz. Alternatively, we could perform high-pass filtering at 0.1 Hz, because previous studies have shown that high-pass filtering instead of bandpass filtering lead to higher reliability and better group discriminability (Shirer et al., 2015). Furthermore, several studies have demonstrated that neural signals existed at higher frequencies than 0.1 Hz (Chen and Glover, 2015; Gohel and Biswal, 2015). Motion censoring, nuisance regression and temporal filtering are performed using AFNI’s *3dTproject* in one step. In addition, the grand mean is scaled to 10000.
8. Calculate functional connectivity (FC) matrix for DK+Aseg/Schaefer+Aseg Atlases. In PhiPipe, there are two versions of FC matrix. The first one is by calculating the Pearson correlation among the mean time series of the brain parcels. This type of FC matrix is the most commonly used one. The second one is by calculating the Pearson correlation among the voxel-wise time series and averaging the Fisher’s Z-transformed voxel-wise correlation based on the brain parcels. To distinguish the two versions, we call the latter one voxel-wise FC matrix (vwFC). The vwFC matrix computation is more time-consuming but conceptually more reasonable to represent the mean functional connectivity strength between any two brain parcels if the functional homogeneity of the brain parcels is uncertain. The FC/vwFC calculation are performed in R (v3.6.1, R Core Team, 2019) and *oro.nifti* package (v0.10.3, Whitcher et al., 2011) was used to read NIFTI file.
9. Calculate regional homogeneity (ReHo, Zang et al., 2004) at each voxel using AFNI’s *3dReHo*. The whole-brain ReHo map is Z-scored and the mean ReHo is extracted for DK+Aseg/Schaefer+Aseg Atlases.
10. Calculate (fractional) amplitude of low frequency fluctuations (ALFF/fALFF, Zang et al., 2007) at each voxel using AFNI’s *3dRSFC*. The whole-brain ALFF/fALFF maps are Z-scored and the mean ALFF/fALFF are extracted for DK+Aseg/Schaefer+Aseg Atlases. For ALFF/fALFF computation, the temporal filtering step aforementioned is omitted. In the computation of parcel-wise brain features, the spatial coverage of each brain parcel (ratio between the voxels with non-zero time series and the total voxels) is also calculated. As there is normally signal loss in orbitofrontal or medial temporal regions for BOLD images due to susceptibility effects, the spatial coverage measure could be used to exclude some brain regions from further analysis.
11. The BOLD images, voxel-wise ReHo and ALFF/fALFF maps are transformed into MNI152 standard space by combining T1-MNI152 and BOLD-T1 registration results to facilitate voxel-wise statistical analysis. The BOLD-T1 registration file is converted into ANTs-compatible format using Convert3D’s *c3d_affine_tool* (v1.0.0, Yushkevich et al., 2006).
12. Quality control pictures of brain extraction, tissue segmentation, parcellation and registration are created for visual check (see Supplementary Figure S2).

The processing steps of DWI images include:

1. Boundary-based registration between b0 image (the first volume without diffusion weighting) and T1 image is performed using FreeSurfer’s *bbregister*.
2. Create masks for whole brain and parcellations based on T1 processing and b0-T1 registration results.
3. Eddy correction, motion correction and outlier replacement are performed using FSL’s *eddy_openmp* (Andersson and Sotiropoulos, 2016). The mean contrast-to-noise ratio after correction (Bastiani et al., 2019) is calculated for quality check.
4. Diffusion tensor model is fitted at each voxel using FSL’s *dtifit*. The fractional anisotropy (FA), mean diffusivity (MD), axial diffusivity (AD) and radial diffusivity (RD) measures are calculated based on the eigenvalues of diffusion tensor. The FA map is registered into the FA58_FMRIB template supplied by FSL using *flirt* (Jenkinson et al., 2002) and *fnirt* (Andersson et al., 2010) to transform the FA/MD/AD/RD maps into the MNI152 space. For multi-shell DWI data, the volumes with lowest non-zero b-value are selected for diffusion tensor calculation.
5. Extract mean FA/MD/AD/RD for JHU Label/Tract Atlases. The JHU Label Atlas consists of 48 white matter regions (Mori et al., 2008), while the JHU Tract Atlas consists of 20 white matter structures (Hua et al., 2008). The JHU Label/Tract Atlases are supplied by FSL.
6. Fiber orientation distribution is estimated at each voxel using FSL’s *bedpostx* (Behrens et al., 2003; Behrens et al., 2007; Jbabdi et al., 2012) and probabilistic tractography is performed for DK+Aseg/Schaefer+Aseg Atlases using FSL’s *probtrackx2*. Structural connectivity (SC) probability between two brain parcels is calculated as the ratio of the number of streamlines reaching the target parcel and the total number of streamlines seeding from the source parcel. The structural connectivity probability matrix is averaged with its transpose to make the matrix symmetric. The element of the matrix represents the likelihood that two brain parcels are connected anatomically.
7. Quality control pictures of brain extraction, parcellation and registration are created for visually check (see Supplementary Figure S3).

### 2.2 Datasets

The demographic characteristics and key acquisition parameters of public datasets used in this study (after quality control) are shown in Table 1. Four datasets with repeated scans, namely, BNU1 (Lin et al., 2015), IPCAS1 (Zhao et al., 2013), IPCAS2, and HNU1 (Chen et al., 2015) for test-retest reliability analysis were chosen from the Consortium for Reliability and Reproducibility (Zuo et al., 2014). The inclusion criteria were: (1) the dataset must have T1, resting-state BOLD and DWI data; (2) the average interval between the first scan and the last scan of each subject should be no longer than two months. And we assumed that there would be no systematic changes taken during this time interval. For BNU1, IPCAS1 and IPCAS2 datasets, each subject had two scans, while for HNU1 dataset, each subject had ten scans. (3) The MRI data should have a whole-brain coverage. Partial coverage of cerebellum was allowed for BOLD and DWI data. The HNU1 dataset had no usable DWI data due to partial brain coverage. However, we still included the HNU1 dataset as it had a different experimental design and we would like to see if the experimental design has a significant impact on the reliability. Three datasets, namely, pNKI (pilot NKI), eNKI (enhanced NKI, Nooner et al., 2012), and SALD (Wei et al., 2018), covering the adult lifespan were selected from the International Neuroimaging Data-Sharing Initiative (Mennes et al., 2013). The inclusion criteria were: (1) the subjects in the dataset should range from early to late adulthood. (2) the dataset should have at least two modalities of T1, resting-state BOLD and DWI data; (3) The MRI data should have a whole-brain coverage. Partial coverage of cerebellum was allowed for BOLD and DWI data. All MRI data were acquired in 3T MRI scanners. More detailed descriptions about these datasets and the IDs of all involved subjects could be found in the Supplementary Section 1. A seemingly trivial yet important consideration in data selection was that all these involved datasets were easily accessible (without complicated application procedures) so that future studies could also use these same datasets for replication and validation.

**Table 1.** The demographic characteristics and key acquisition parameters of MRI data used in the reliability and predicative validity analyses.

| Dataset | N<sup>a</sup> | Age<sup>b</sup> | Sex<sup>c</sup> | T1 | BOLD |  |  |  | DWI |  |  |
| --- | --- | --- | --- | --- | --- | --- | --- | --- | --- | --- | --- |
|  |  |  |  | Res<sup>d</sup> | Res | TR<sup>e</sup> | Vol<sup>f</sup> | Res | Vol | b-value | Dir<sup>g</sup> |
| Reliability |  |  |  |  |  |  |  |  |  |  |  |
| BNU1 | 45 | 19-27 | 24/21 | 1.3*1.0*1.0 | 3.1*3.1*4.2 | 2.0 | 200 | 2.2*2.2*2.2 | 31 | 1000<sup>h</sup> | 30 |
| IPCAS1 | 18 | 18-24 | 4/14 | 1.3*1.0*1.0 | 4.0*4.0*4.8 | 2.0 | 205 | 1.8*1.8*2.5 | 62 | 1000 | 60 |
| IPCAS2 | 27 | 12-15 | 10/17 | 1.2*0.9*0.9 | 3.7*3.7*4.0 | 2.5 | 212 | 1.9*1.9*3.0 | 39 | 1000 | 36 |
| HNU1 | 22 | 21-30 | 11/11 | 1.0*1.0*1.0 | 3.4*3.4*3.4 | 2.0 | 300 | \ | \ | \ | \ |
| Validity |  |  |  |  |  |  |  |  |  |  |  |
| pNKI | 113 | 18-85 | 49/64 | 1.0*1.0*1.0 | 3.0*3.0*3.3 | 2.5 | 260 | 2.0*2.0*2.0 | 76 | 1000 | 64 |
| eNKI | 502 | 18-83 | 251/251 | 1.0*1.0*1.0 | 3.0*3.0*3.3 | 2.5 | 120 | 2.0*2.0*2.0 | 137 | 1500 | 128 |
| SALD | 414 | 19-80 | 154/260 | 1.0*1.0*1.0 | 3.4*3.4*4.0 | 2.0 | 242 | \ | \ | \ | \ |

<sup>a</sup> sample size
<sup>b</sup> age range (in years)
<sup>c</sup> sample sizes for male/female
<sup>d</sup> spatial resolution (i.e., voxel size, in millimeter)
<sup>e</sup> repetition time (in seconds)
<sup>f</sup> the number of volumes
<sup>g</sup> the number of diffusion gradient directions
<sup>h</sup> the unit is s/mm<sup>2</sup>

### 2.3 Brain features, Quality control and Processing Variants

The seven datasets were first processed using PhiPipe with the default steps and parameters. The brain features included CT, CA, CV and SV for T1 data and DK+Aseg/Schaefer+Aseg Atlases; ALFF, fALFF, ReHo, FC, and vwFC for resting-state BOLD data and DK+Aseg/Schaefer+Aseg Atlases; FA, MD, AD, RD for JHU Label/Tract Atlases and DWI data, and SC for DK+Aseg/Schaefer+Aseg Atlases and DWI data. The FC matrix was Fisher-Z transformed before reliability and validity analysis. Different from FC matrix, SC matrix is inherently sparse (with many zero connections), so only the non-zero edges (connectivity measure between two parcels) across subjects in a dataset for SC measures were used for downstream analysis. In the Results section, we only presented the results for Schaefer+Aseg Atlas and JHU Tract Atlas and the results for DK+Aseg Atlas and JHU Label Atlas were reported in the Supplementary Materials. The conclusions drawn in this study were not dependent on the specific atlas used. The codes to invoke PhiPipe could be found in Supplementary Section 2. The runtime and peak memory usage for processing each modality of a typical subject could be found in the Supplementary Table S2.

The raw data, brain extraction, segmentation, parcellation and registration results of all data modalities were visually checked. Of note, the defacing operation for T1 data would sometimes remove gray matter unintentionally. For resting-state BOLD or DWI data, the number of volumes were sometimes smaller than the expected one for unknown causes. Quantitively, for the T1 data, data were excluded if the image quality rating score outputted by CAT12 was lower than 0.8 (B level); For the resting-state BOLD data, data were excluded if the mean FD > 0.5mm, or the motion outlier ratio > 0.2, or the mean FD was larger than Q3 (the third quantile) + 1.5*IQR (inter-quantile range) in the dataset (see Supplementary Table S3 for a summary of head motion metrics in all datasets). For the DWI data, data were excluded if the mean contrast-to-noise ratio after eddy/motion correction were lower than Q1 (the first quantile) - 1.5*IQR in the dataset. In order to keep the same samples in each modality, if the data of one modality was excluded after quality control, the data of the other modalities were also excluded.

Then the seven datasets were processed using the alternative steps and parameters (see Table 2). The processing variants included: (1) In the T1 processing, the skull stripping was performed using CAT12 instead of FreeSurfer’s *recon-all*. As the BOLD/DWI processing was based on T1 processing, we also examined whether the CAT12’s skull stripping would have an effect on the results of BOLD and DWI processing. (2) In the T1 processing, the TIV was estimated using CAT12 instead of FreeSurfer’s *recon-all*. (3) In the BOLD processing, the steps of slice timing correction, temporal filtering, nuisance regression were manipulated separately to see their independent influence on the reliability and validity. In slice timing correction, we omitted this step (NOSTC); (4) In temporal filtering, we didn’t use low-pass filtering at 0.1 Hz (NOLP); (5) In the nuisance regression, we included the global signal regression (GSR); (6) In addition, we examined the effect of calculation in MNI152 space on the reliability and validity of brain features. By default, all brain features were calculated in subject’s native space in PhiPipe. However, a common practice is transforming subject’s data into standard space and calculating brain features in the standard space. Therefore, we also calculated the same brain features using the same atlases in the MNI152 space. For ALFF/fALFF/ReHo, the voxel-wise measures calculated in the native space were transformed into MNI152 space. The mean values were extracted for each brain feature based on the atlases in MNI152 space. For FC/vwFC, the time series was transformed into MNI152 space for calculation.

**Table 2.**
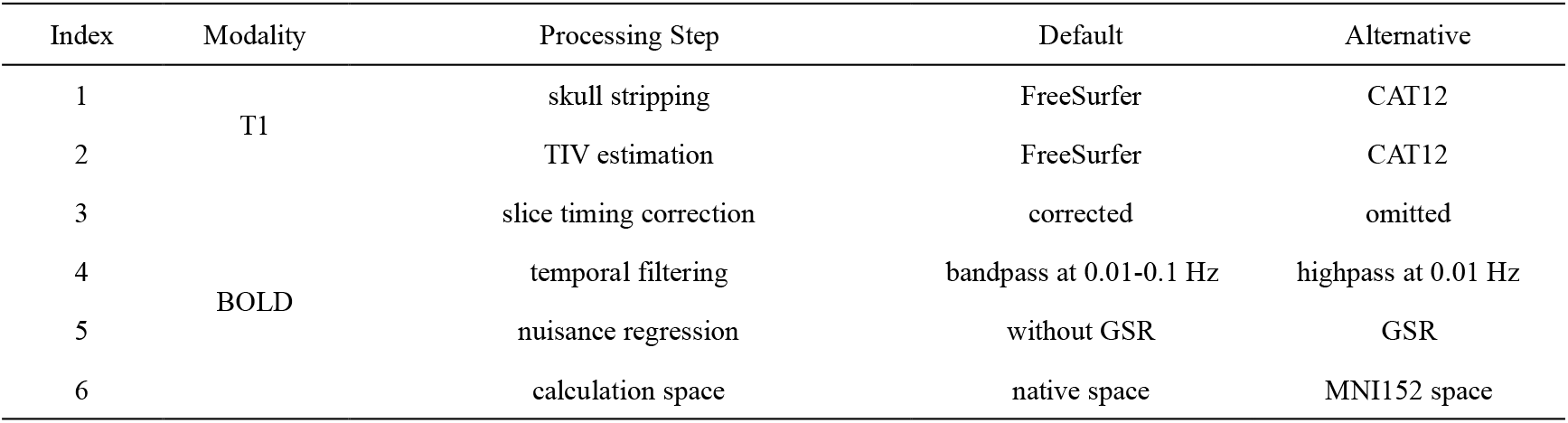
Alternative processing options compared with default settings in reliability and validity assessment.

Finally, in order to compare the results of PhiPipe with other commonly used pipelines, we processed the same datasets using DPARSF (v5.2) and PANDA (v1.3.1). DPARSF was used for BOLD processing, and PANDA was used for DWI processing. In both DPARSF and PANDA, the T1 data was also used for registration. We used the default setting of all parameters for DPARSF and PANDA by assuming that the default parameters were the recommended parameters by the developers and also the most commonly used parameters by the end users. A detailed list of these parameters and more detailed descriptions about the DPARSF/PANDA processing and quality check could be found in Supplementary Section 3-4. For DPARSF and PANDA, only the Schaefer+Aseg Atlas and JHU Tract Atlas were used.

### 2.4 Reliability Analysis

Intra-class correlation (ICC) was the most commonly used measure in assessing the test-retest reliability in this field. Conceptually, the ICC could be formulated as the ratio between the variance of subjects and the observed total variance (Liljequist et al., 2019). The observed total variance consists of the variance due to subjects, the variance due to systematic bias between scans, and the variance due to unaccounted random errors.

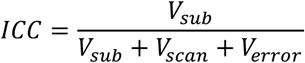

The ICC was estimated by a two-way random-effect ANOVA model using *irr* package (v0.84.1) of R. Theoretically, the ICC ranges from 0 to 1. However, the ICC estimates could be negative. As a common practice, these negative ICC estimations were set to 0 (i.e., totally unreliable). The ICC used in this study was known as ICC(2,1) or ICC(A,1), as there were other forms of ICC (Shrout and Fleiss, 1979; McGraw and Wong, 1996). In order to summarize results, we classified ICC into three levels: Poor (ICC < 0.5), Moderate (0.5 <= ICC <= 0.75), and Good (ICC > 0.75). An ICC of 0.5 means that the true variance of subjects is as large as other sources of variance and an ICC of 0.75 means that the true variance of subjects is three times as large as other sources of variance. Of note, this classification is for simplifying results and whether the ICC is good enough depends on the specific application.

The ICC was calculated for each parcel or edge for all brain features. The mean ICC of all parcels or edges for a brain feature was used to reflect the overall reliability. A bootstrap method was used to compare the overall reliability between default setting and processing variants using PhiPipe, or between PhiPipe and other pipelines (i.e., DPARSF and PANDA). For instance, if we want to assess the influence of CAT12’s skull stripping on the mean ICC of cortical thickness, the subjects were resampled with replacement for 2000 times. In each resample, the mean ICC were calculated for the results with or without CAT12’s skull stripping separately. The difference between the mean ICC with and without CAT12 was further calculated in this resample. The 95% Bias-corrected and accelerated (Bca) confidence interval (CI) was calculated based on the 2000 resamples to infer the statistical significance of ICC difference (Diciccio and Efron, 1996). If the CI does not include the zero, the ICC difference is significant. The codes for ICC, bootstrap and CI calculation were included in the Supplementary Section 5. In the current study, all the p values or confidence intervals were not corrected for multiple comparisons, as it was difficult to define the sets of comparisons should be corrected. Thus, the uncorrected results could be easier interpreted.

To quantify the reproducibility of the ICC results across different datasets, we used two reproducibility measures: (1) the Spearman (rank) correlation of parcel-wise or edge-wise ICC between two datasets for a brain feature was calculated and this procedure was repeated for all dataset pairs. The mean correlation of all dataset pairs was used to represent the relative consistency of ICC across different datasets. A high correlation means that the parcel or edge had higher ICC in one dataset would also have higher ICC in another dataset. (2) the mean absolute difference (MAD) of parcel-wise or edge-wise ICC between two datasets for a brain feature was calculated and this procedure was repeated for all dataset pairs. The mean MAD of all dataset pairs was used to represent the absolute agreement of ICC across different datasets. A low MAD means that the parcel or edge had an ICC of, for instance, 0.1 in one dataset would also have an ICC of about 0.1 in another dataset. To make the results comparable across MRI modalities, only the datasets with three modalities were included.

### 2.5 Predicative Validity Analysis

Pearson correlation coefficient between each brain feature and age was calculated for each parcel or edge to assess the predicative validity. Previous studies have shown that there were a linear or monotonous relationship between age and brain features in general during the adult lifespan, especially after the mid-life (for instance, Lebel et al., 2012; Onoda et al., 2012; Betzel et al., 2014; Beck et al., 2021; Dima et al., 2022; Frangou et al., 2022). To distinguish the correlation between brain feature and age with other correlation analyses in the current study, Age-R was used to denote specifically the correlation between a brain feature and age. We should note that there is a direct relationship between observed Age-R and ICC (Elliott et al., 2020), as shown in the formula below. As we could assume the ICC of age is one (perfect reliable), the square root of ICC for a brain feature sets the upper bound of the observed Age-R.

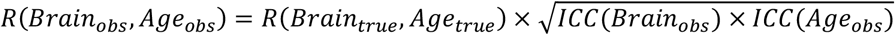

In order to summarize the results, we classified the Age-R into three levels: Low (|Age-R| < 0.1), Moderate (0.1 <= |Age-R| <= 0.3), High (|Age-R| > 0.3). The classification criterion was based on the normal range of correlation observed in previous studies (Elliott et al., 2020; Marek et al., 2022). As in the case of ICC, whether the Age-R is high enough depends on the specific application. It should also be noted that a brain feature with high Age-R just means that this feature has high validity in predicting age but does not mean this feature is better than the features with low Age-R. The comparison of brain features is valid only when they measure the same brain properties.

The Age-R was calculated for each parcel or edge for all brain features. As the Age-R could be positive or negative, in order to summarize results, we squared the Age-R (Age-R-Squared) to reflect the amount of variance explained by the brain features. The mean Age-R-Squared of all parcels or edges for a brain feature was used to reflect the overall validity. To compare the mean Age-R-Squared between default setting and processing variants using PhiPipe, or between PhiPipe and other pipelines, the same bootstrap method was used as in the case of ICC.

To exclude the confounding effects of sex and TIV, we repeated the analysis using partial correlation with sex and TIV as covariates. To exclude the influence of potential outliers and non-Gaussian distribution, we repeated the Age-R analysis using Spearman correlation. To examine whether there is a non-linear relationship between age and brain features, we repeated the analysis using a multiple regression model with both the linear and quadratic terms of age. If the coefficient of quadratic term was significant, it means that the brain-age relationship is significantly non-linear. Furthermore, we assessed whether the quadratic model is better than the linear regression model based on the Akaike’s Information Criterion (AIC). More complex non-linear brain-age relationships were not examined as the current sample size was still limited.

To quantify the reproducibility of the Age-R results across different datasets, we used two reproducibility measures: (1) the Spearman correlation of Age-R between two datasets for a brain feature was calculated and this procedure was repeated for all dataset pairs. The mean correlation of all dataset pairs was used to represent the relative consistency of Age-R across different datasets. A high correlation means that the parcel or edge had higher Age-R in one dataset would also have higher Age-R in another dataset. (2) the mean absolute difference (MAD) of Age-R between two datasets for a brain feature was calculated and this procedure was repeated for all dataset pairs. The mean MAD of all dataset pairs was used to represent the absolute agreement of Age-R across different datasets. A low MAD means that the parcel or edge had an Age-R of −0.1 in one dataset would also have an Age-R of about −0.1 in another dataset. To make the results comparable across MRI modalities, only the datasets with three modalities were included.

In addition, since group difference detection was widely used in previous pipeline validations, and group comparisons are common in study designs, we further examined the validity of the brain features in terms of effect size of age group differences. Specifically, we split the subjects into younger and older groups at a cutoff age of 40 years and compared the means between the two groups using Welch’s two sample T-test for all brain features, atlases and datasets. The mean of absolute Cohen’s d of all parcels or edges for a brain feature was used as a measure of overall group difference. To summarize results, the Cohen’s d was classified into three levels: Small (|d| < 0.2), Moderate (0.2 <= |d| < 0.6), and Large (|d| >= 0.6). Cohen’s d was calculated using the *effsize* (v0.8.1, Torchiano, 2020) package of R. More details about the demographics of the younger and older groups were presented in Supplementary Table S4.

### 2.6 Multivariate Reliability and Predicative Validity Analysis

As multivariate models are becoming more and more popular in this field, we also assessed multivariate reliability and predicative validity of the brain features. For multivariate reliability, we used a recently proposed distance-based ICC (dbICC) statistic (Xu et al., 2021). Bootstrap with 2000 resamplings was used to estimate the 95% confidence interval. For multivariate validity, a linear-kernel support vector regression (SVR) model with nested 10-fold cross-validation (CV) was used for age predication. Specifically, the data was randomly divided into 10 folds, of which 9 folds were used for model training and the remaining fold was used for model testing. The training/testing processes were repeated 10 times so that each fold would be used for model testing once. Within the model training process, a 10-fold CV was used again to find the optimal hyperparameters via grid search. The correlation between the real and predicated age (denoted as CV-R) was used as the measure of multivariate predicative validity. Besides SVR, Connectome-based Predicative Modeling (CPM, Shen et al., 2017) was used for age predication. The same nested 10-fold cross validation was applied as in SVR. The multivariate analyses were only conducted on the brain features generated by the default settings of PhiPipe. The *dbicc* (v0.13) package of R was used for dbICC calculation. The *e1071* (v1.7-8, Meyer et al., 2021) package of R was used for SVR modeling. The codes to perform dbICC calculation and SVR-/CPM-based age predication were presented in Supplementary Section 6.

The visualization of results in the current study was performed using *ggplot2* (v3.3.2, Wickham, 2016), *corrplot* (v0.92, Wei and Simko, 2021) and *ggseg* (v1.6.4, Mowinckel and Vidal-Piñeiro, 2020) packages of R.

## 3. Results

### 3.1 The reliability and validity of brain features using the default settings of PhiPipe

Figure 2A presented the mean and standard deviation of ICC results for all brain features and datasets. In addition, Table 3 showed the percentage of significant ICCs and reliability levels for all brain features in the BNU1 dataset. In Figure 3, the parcel-wise or edge-wise ICCs were shown for all brain features in the BNU1 dataset. In consistent with previous studies (for instance, Zuo and Xing, 2014; Chen et al., 2015; Noble et al., 2019; Buimer et al., 2020; Drobinin et al., 2020), both T1 and DWI brain features showed moderate or good overall reliability, while the resting-state BOLD brain features showed poor or moderate reliability. For T1 data, the CA showed the highest and almost perfect reliability, while the CT showed the lowest reliability. For resting-state BOLD data, the ALFF showed the highest reliability while the FC/vwFC showed the lowest reliability. For DWI data, the FA showed the highest reliability while the SC showed the lowest reliability. The results for other atlases and datasets could be found in the Supplementary Table S5-S8 and Figure S4-S6.

**Figure 2.**
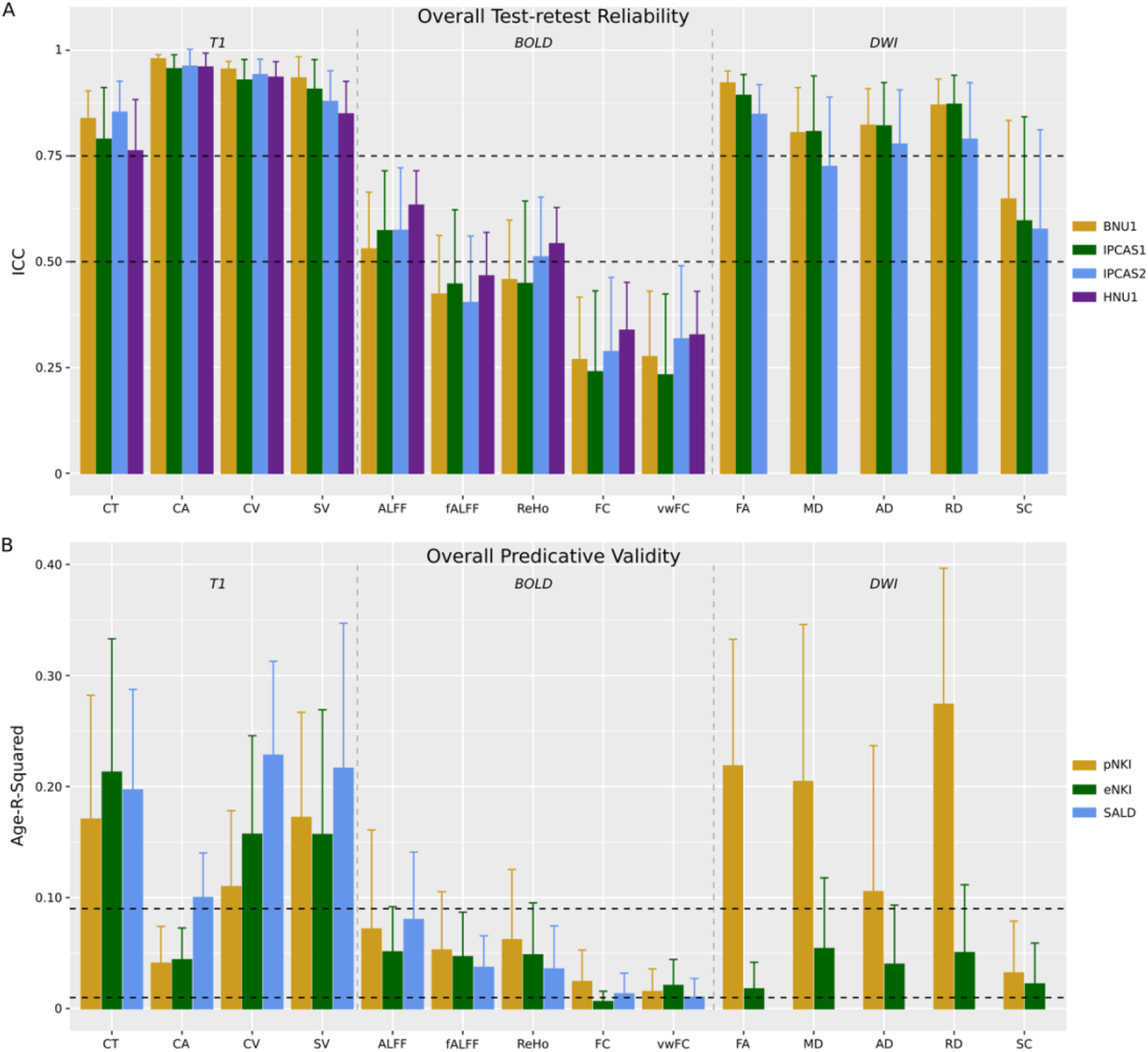
The mean and standard deviation of ICC and Age-R-Squared (the squared correlation between age and a brain feature) for all brain features across all datasets. The dashed horizontal lines indicated different levels of reliability and validity. For instance, ICCs above the line of ICC=0.75 means good reliability based on the criteria in the current study.

**Table 3.** The summary of reliability for all brain features in the BNU1 dataset.

| Modality | Feature | Atlas | N<sup>a</sup> | ICC<sup>b</sup> | Sig<sup>c</sup> | Poor<sup>d</sup> | Moderate<sup>d</sup> | Good<sup>d</sup> |
| --- | --- | --- | --- | --- | --- | --- | --- | --- |
| T1 | CT | Schaefer | 100 | 0.84 (0.065) | 100 | 0 | 12 | 88 |
|  | CA | Schaefer | 100 | 0.98 (0.010) | 100 | 0 | 0 | 100 |
|  | CV | Schaefer | 100 | 0.96 (0.019) | 100 | 0 | 0 | 100 |
|  | SV | Aseg | 14 | 0.93 (0.050) | 100 | 0 | 0 | 100 |
| BOLD | ALFF | Schaefer+Aseg | 114 | 0.53 (0.13) | 99 | 38 | 59 | 3 |
|  | fALFF | Schaefer+Aseg | 114 | 0.42 (0.14) | 88 | 68 | 32 | 0 |
|  | ReHo | Schaefer+Aseg | 114 | 0.46 (0.14) | 90 | 52 | 48 | 0 |
|  | FC | Schaefer+Aseg | 6441 | 0.27 (0.15) | 56 | 94 | 6 | 0 |
|  | vwFC | Schaefer+Aseg | 6441 | 0.28 (0.16) | 58 | 92 | 8 | 0 |
| DWI | FA | JHU-Tract | 20 | 0.92 (0.028) | 100 | 0 | 0 | 100 |
|  | MD | JHU-Tract | 20 | 0.81 (0.11) | 100 | 0 | 35 | 65 |
|  | AD | JHU-Tract | 20 | 0.82 (0.087) | 100 | 0 | 20 | 80 |
|  | RD | JHU-Tract | 20 | 0.87 (0.061) | 100 | 0 | 5 | 95 |
|  | SC | Schaefer+Aseg | 854 | 0.65 (0.19) | 97 | 21 | 46 | 33 |

<sup>a</sup> the number of parcels or edges for a brain feature
<sup>b</sup> the mean (standard deviation) of ICC
<sup>c</sup> the percentage of significant ICCs. The significance level was $p < 0.05$ (uncorrected for multiple comparisons).
<sup>d</sup> the percentage of ICCs belonging to different levels of reliability (Poor/Moderate/Good)

**Figure 3.**
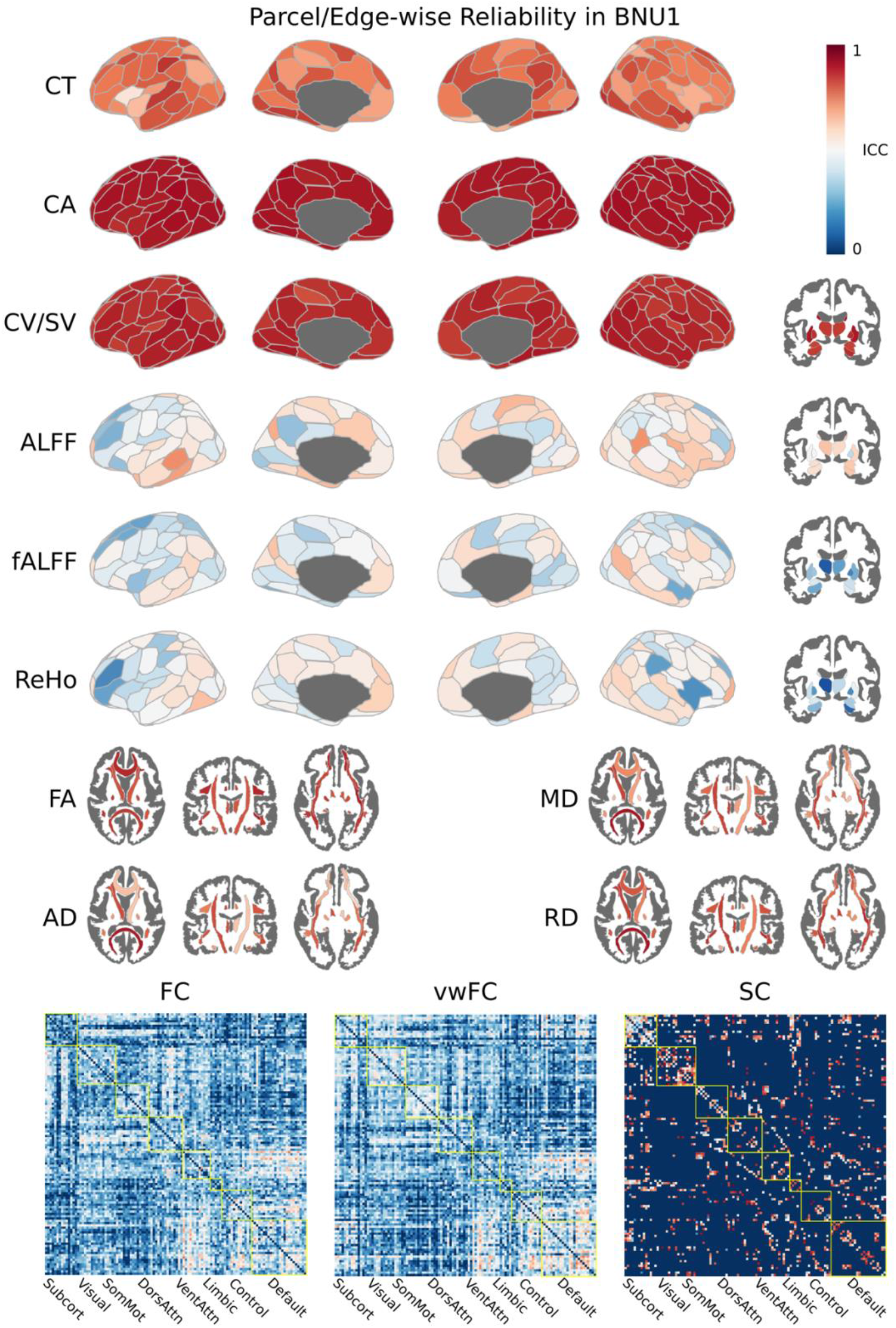
The parcel-wise and edge-wise ICC for all features in the BNU1 dataset. Blue means ICC < 0.5, and red means ICC > 0.5. For the connectivity matrices, the elements are ordered based on Yeo’s 7-network parcellation and subcortical regions (indicated by yellow lines).

Figure 2B presented the mean and standard deviation of Age-R-Squared results for all brain features and datasets. In addition, Table 4 showed the percentage of significant, positive and negative Age-Rs and validity levels for all brain features in the pNKI dataset. In Figure 4, the parcel-wise or edge-wise Age-Rs were shown for all brain features in the pNKI dataset. Both T1 and DWI brain features showed moderate or high overall validity, while the resting-state BOLD brain features showed low or moderate validity. The lower validity of BOLD brain features was probably due to the lower reliability. For T1 data, the CT/CV/SV showed much higher validity than the CA. A possible reason is that the cortical area calculated in FreeSurfer by default is the area of white surface (i.e., the surface between gray matter and white matter) and this brain feature may be less sensitive to age. Almost all T1 brain features showed negative Age-R, in consistent with previous findings (Dima et al., 2022; Frangou et al., 2022). For resting-state BOLD data, the ALFF/fALFF/ReHo showed similar validity while the FC/vwFC showed the lowest validity. In most cases, the ALFF/fALFF/ReHo showed negative Age-R in cortical regions and positive Age-R in subcortical regions, which was consistent with a study using brain features characterizing local FC (Wen et al., 2020). For FC/vwFC, negative Age-Rs were more likely to be observed in within-network edges while positive Age-Rs were more likely to be observed in between-network edegs, in agreement with previous findings (Betzel et al., 2014; Damoiseaux, 2017). There were also significant difference between the FC and vwFC features, which may indicate that the low Age-R for a specific edge was unstable. For DWI data, we found that the validity had a big difference in two datasets, which probably reflects the influence of different scanning parameters. For instance, the b-value was 1000 s/mm^2^ for pNKI dataset while 1500 s/mm^2^ for eNKI dataset. In most cases, FA/SC showed negative Age-R while MD/RD showed positive Age-R. The Age-R pattern of AD is more variable. These results conformed to previous findings well (Gong et al., 2009; Westlye et al., 2010; Lebel et al., 2012; Beck et al., 2021). The results for other atlases and datasets could be found in the Supplementary Table S9-S11 and Figure S7-S8. As shown in Supplementary Table S12-S17 and Figure S9, using sex and TIV covariates or Spearman correlation did not change the general pattern of results. In most cases, the quadratic term of age was not significant for T1 and BOLD brain features. The quadratic trends were significant for most FA/MD/AD/RD measures and the inclusion of quadratic term could explain about 5% more variances in the current datasets (for more details, see Supplementary Table S18-S20). The quadratic trends of diffusion tensor brain features observed in the current study were also reported in previous findings (Westlye et al., 2010; Lebel et al., 2012; Beck et al., 2021).

**Table 4.**
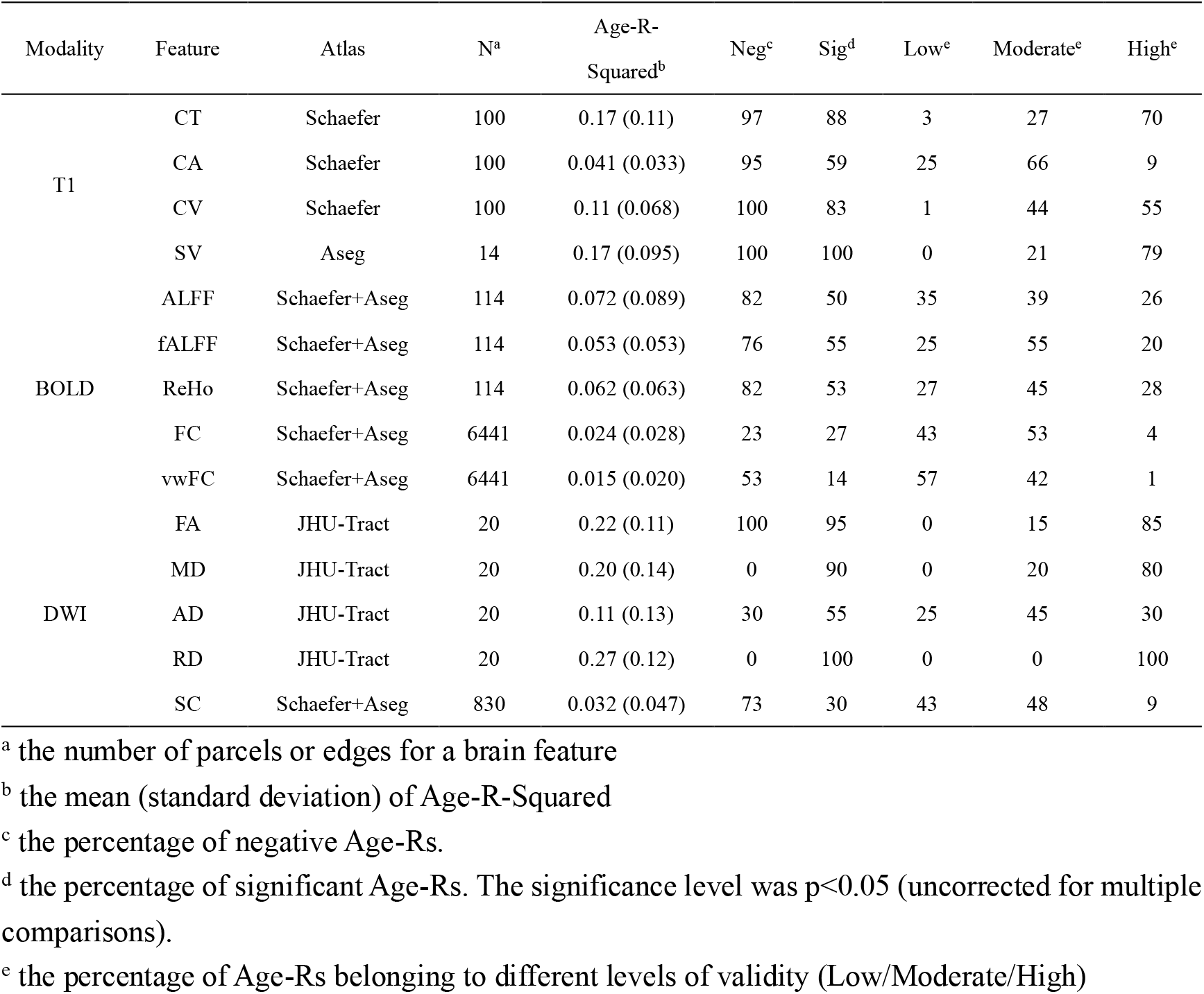
The summary of predicative validity for all brain features in the pNKI dataset.

**Figure 4.**
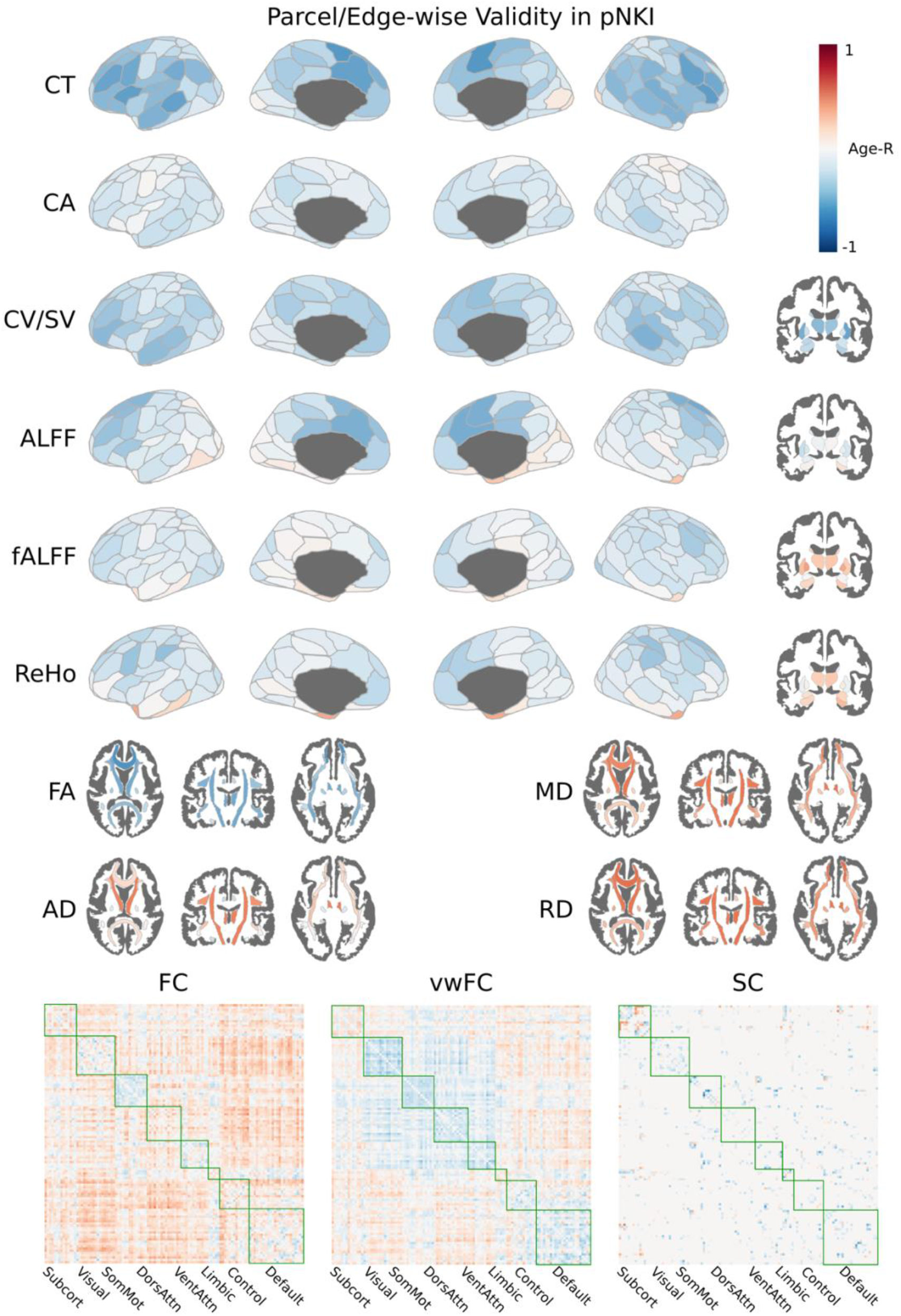
The parcel-wise and edge-wise Age-R (the correlation between age and a brain feature) for all brain features in the pNKI dataset. Blue indicates negative Age-R, and red indicates positive Age-R. For the connectivity matrices, the elements are ordered based on Yeo’s 7-network parcellation and subcortical regions (indicated by green lines).

The group difference results showed that T1 and DWI features had moderate or large overall group difference, while the resting-state BOLD features had small or moderate group difference. The younger group had higher CT/CA/CV/SV/FA, and lower MD/RD than the older group in almost all brain regions. More details about the group difference results were presented in Supplementary Table S22-S24 and Figure S10.

As shown in Figure 5A, the between-dataset correlation of ICCs in most cases were lower than 0.5 for brain features of T1 and DWI data and lower than 0.25 for brain features of resting-state BOLD data, which means the rank of parcel-wise or edge-wise ICC was not preserved across datasets. As shown in Figure 5B, the MAD of ICC results in most cases were lower than 0.1 for brain features of T1 and DWI data and higher than 0.15 for brain features of resting-state BOLD data.

**Figure 5.**
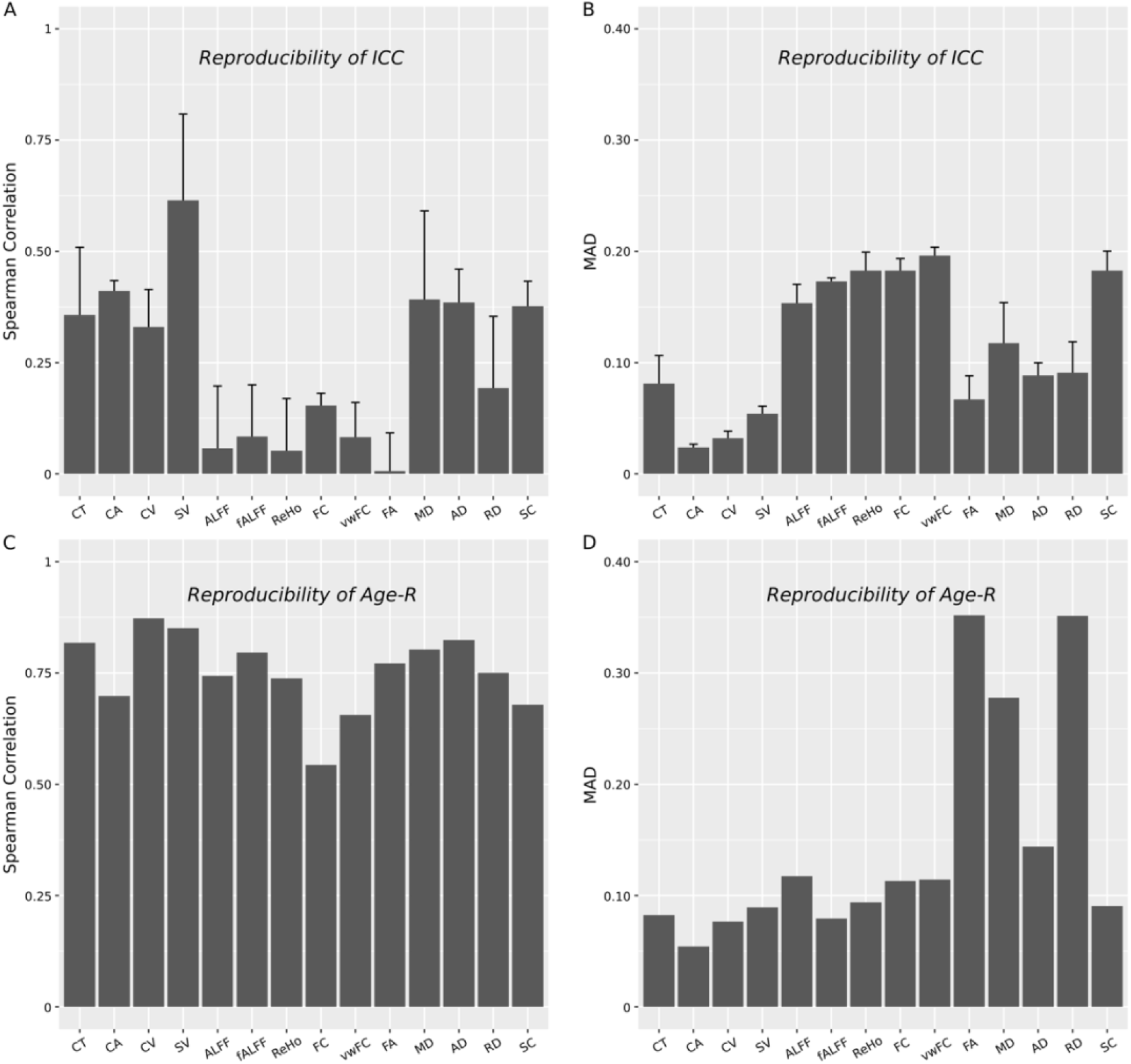
The reproducibility of ICC and Age-R for all brain features across all datasets. The data was presented as mean and standard deviation of reproducibility among dataset pairs. For Age-R analysis, only two datasets (pNKI and eNKI) were involved so that there was no standard deviation.

The between-dataset correlation of Age-R results in most cases were about 0.75 for all brain features (Figure 5C), which means the rank of parcel-wise or edge-wise Age-R was preserved across datasets. The MAD of Age-R results were about 0.1 for brain features of T1 and resting-state BOLD fMRI data and higher than 0.25 for diffusion tensor features of DWI data (Figure 5D). These results provided a reference about the reliability and validity changes we could expect from using a new dataset. More details about the reproducibility of ICC and Age-R results could be found in Supplementary Table S21.

### 3.2 The influence of processing variants on reliability and validity

As shown in Figure 6A, in most cases, using CAT12 instead of FreeSurfer for skull stripping had little impact on the ICC results of all brain features. As shown in Figure 6B, using CAT12’s skull stripping in some cases led to decreased Age-R-Squared for CT and CV but had little impact on other brain features. As shown in Table 5, using CAT12 instead of FreeSurfer for TIV estimation in general led to higher reliability, although the improvement was not statistically significant. As the ICC of TIV was almost perfect (>0.98), we used the coefficient of variation to assess the reliability of TIV estimation. Using CAT12’s TIV estimation in general led to significantly lower Age-R. As in the ideal case, the correlation between TIV and age should be zero, the lower Age-R using CAT12’s TIV estimation means higher validity. We should also note that there were significant Age-R in the SALD dataset, which may indicate that there was a generation bias in this sample. The statistical inference of Age-R difference between CAT12 and FreeSurfer for TIV estimation was performed using *cocor* package (v1.1-3, Diedenhofen and Musch, 2015) of R. More numerical details could be found in Supplementary Table S25-S26.

**Figure 6.**
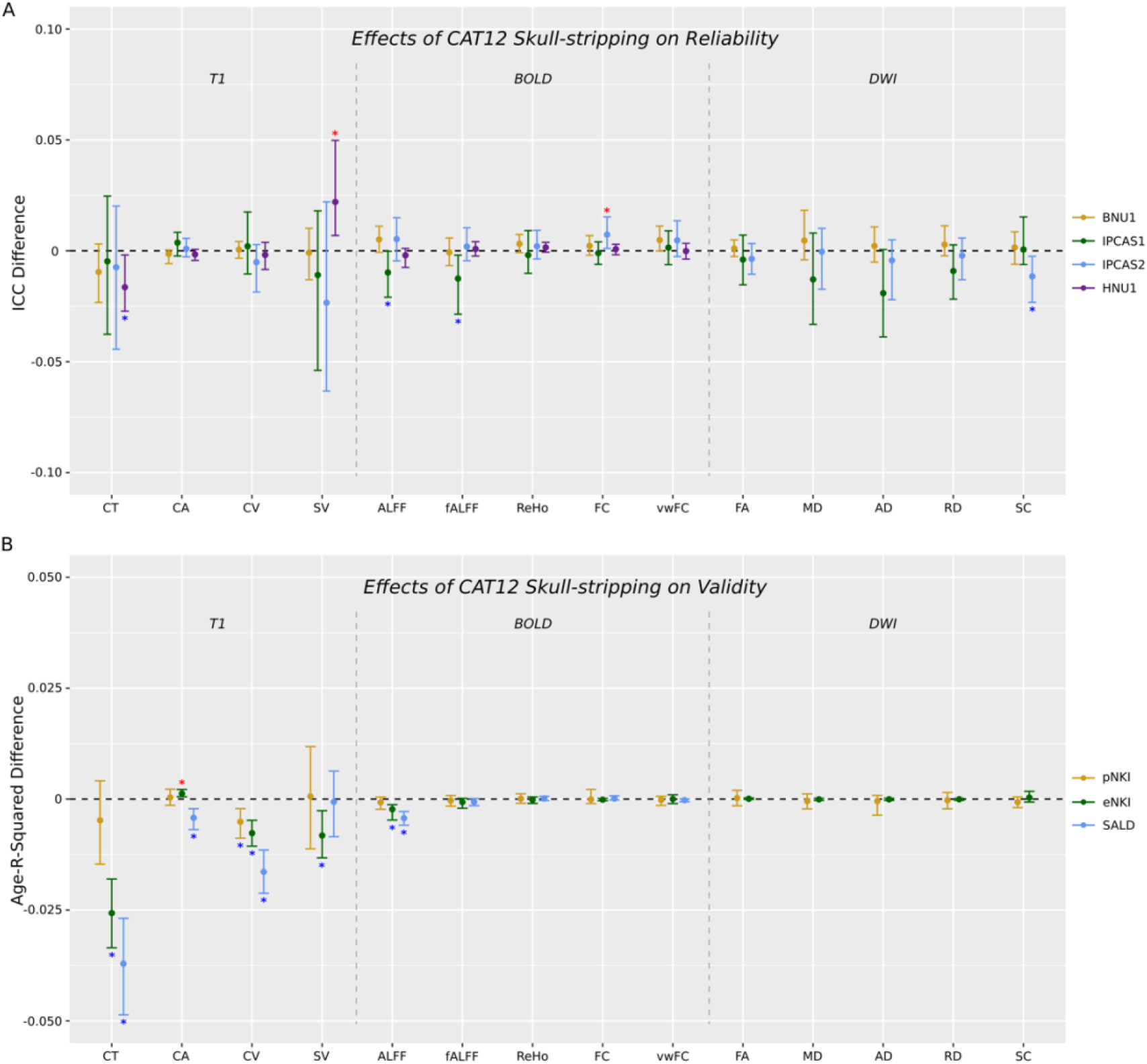
The effect of CAT12’s skull-stripping on the ICC and Age-R-squared for all brain features across all datasets. The Y axis means the difference of mean ICC or Age-R-Squared between the results using default and alternative parameters and settings. The data are presented as (alternative - default), so a positive value means increased reliability and validity using alternative options compared with the default settings. The error bars represent 95% CIs. If the CI does not contain zero, the difference is statistically significant. The red/blue asterisks indicate significant increase or decrease.

**Table 5.** The reliability and validity of TIV estimated by FreeSurfer and CAT12

| Dataset | Method | TIV<sup>a</sup> | CoV/Age-R<sup>b</sup> | T/Z<sup>c</sup> | Correlation<sup>d</sup> |
| --- | --- | --- | --- | --- | --- |
| Reliability |  |  |  |  |  |
| BNU1 | FreeSurfer | 1546.64 (174.80) | 1.03 (2.38) | -2.01 (0.05) | 0.87 |
|  | CAT12 | 1468.89 (140.97) | 0.29 (0.28) |  |  |
| IPCAS1 | FreeSurfer | 1471.03 (117.77) | 0.56 (0.62) | -1.62 (0.12) | 0.93 |
|  | CAT12 | 1405.88 (106.98) | 0.32 (0.16) |  |  |
| IPCAS2 | FreeSurfer | 1490.33 (114.06) | 0.58 (0.52) | -1.98 (0.059) | 0.84 |
|  | CAT12 | 1423.79 (101.13) | 0.34 (0.33) |  |  |
| HNU1 | FreeSurfer | 1501.57 (141.96) | 0.81 (0.71) | 0.34 (0.74) | 0.89 |
|  | CAT12 | 1521.77 (146.75) | 0.88 (0.63) |  |  |
| Validity |  |  |  |  |  |
| pNKI | FreeSurfer | 1517.47 (208.50) | 0.11 | 1.37 (0.17) | 0.82 |
|  | CAT12 | 1450.09 (149.36) | 0.034 |  |  |
| eNKI | FreeSurfer | 1499.62 (158.02) | -0.099 | -3.20 (0.0014) | 0.94 |
|  | CAT12 | 1438.33 (145.05) | -0.049 |  |  |
| SALD | FreeSurfer | 1474.70 (139.20) | -0.36 | -5.28 (<0.001) | 0.94 |
|  | CAT12 | 1408.03 (129.32) | -0.28 |  |  |

<sup>a</sup> the mean (standard deviation) of TIV in mm<sup>3</sup>
<sup>b</sup> the mean (standard deviation) of Coefficient of Variation (CoV) for TIV, and the correlation between TIV and age (Age-R)
<sup>c</sup> the T/Z value (p value) in comparing the CoV and Age-R between two methods. For CoV, the paired t-test was used. For Age-R, the Pearson-Filon's z test was used.
<sup>d</sup> the correlation of estimated TIVs between the two methods. For each subject in the test-retest datasets, the estimated TIVs of repeated scans were averaged.

As shown in Figure 7A, NOSTC had little and inconsistent impact on the reliability of BOLD brain features; NOLP consistently improved the reliability of ReHo, FC and vwFC; GSR had inconsistent effects on reliability across brain features and datasets; calculation in MNI space increased the reliability of ALFF and FC. As shown in Figure 7B, NOSTC improved the validity of ALFF with a very small effect size; NOLP consistently increased the validity of ReHo; GSR consistently increased the validity of fALFF; calculation in MNI space led to decreased validity of fALFF and ReHo. In summary, the effects of processing variants in general were dependent on the brain features and datasets, and NOLP showed the most consistent and largest impact on reliability and validity. These results were consistent with previous findings (Wu et al., 2011; Shirer et al., 2015; Murphy and Fox, 2017). More numerical details could be found in Supplementary Table S27-S28.

**Figure 7.**
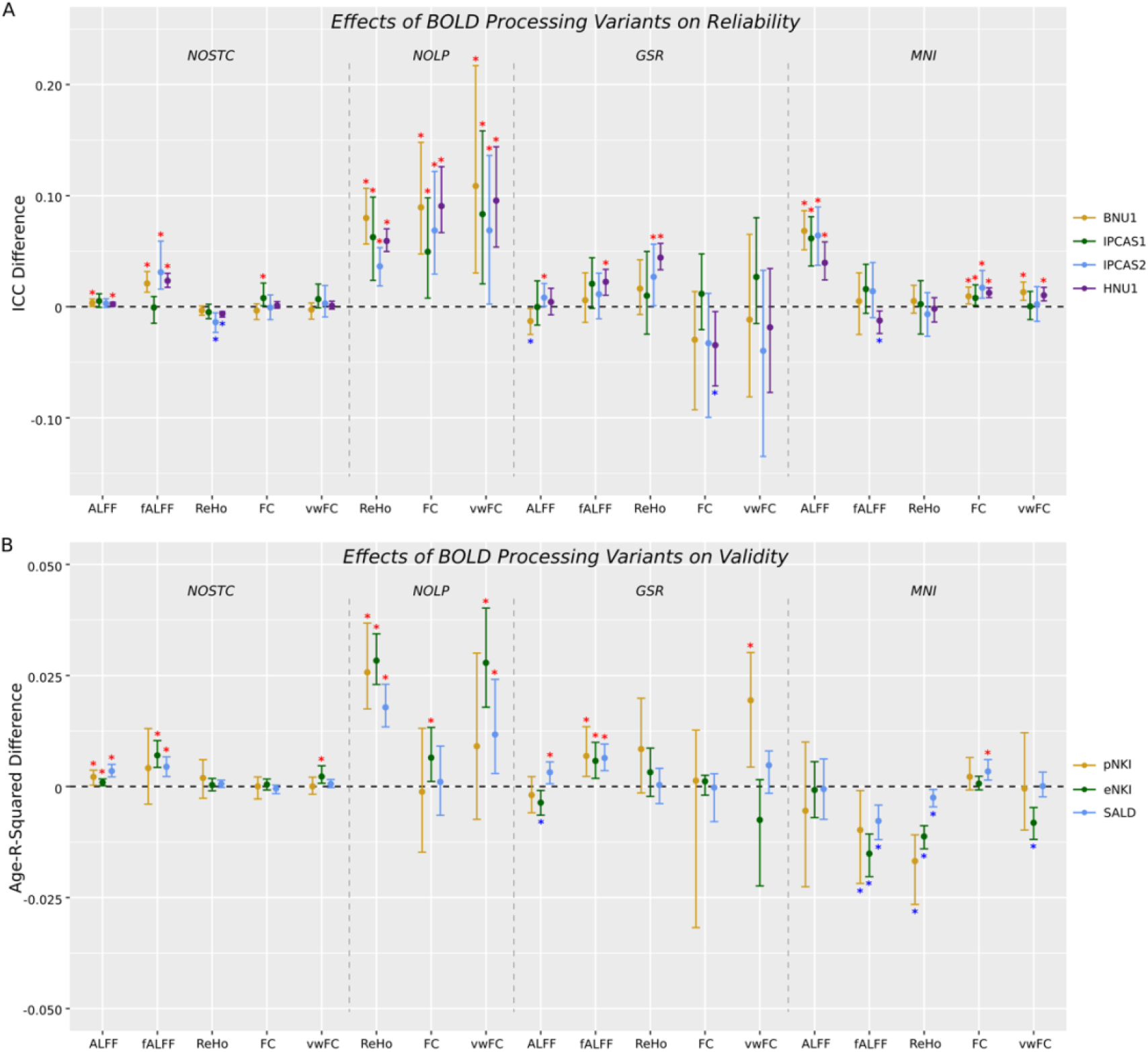
The effects of NOSTC, NOLP, GSR and MNI on the ICC and Age-R-squared for resting-state BOLD brain features across all datasets. The Y axis means the difference of mean ICC or Age-R-Squared between the results using default and alternative parameters and settings. The data are presented as (alternative - default), so a positive value means increased reliability and validity using alternative options compared with the default settings. The error bars represent 95% CIs. If the CI does not contain zero, the difference is statistically significant. The red/blue asterisks are used to indicate significant increase or decrease.

### 3.3 Comparisons with DPARSF and PANDA

The comparisons of reliability and validity between PhiPipe and DPARSF/PANDA were presented in Figure 8. In order to summarize results, we defined that if the significant differences were detected in all datasets between PhiPipe and DPARSF/PANDA, we could say that the PhiPipe had better or worse reliability and validity. Otherwise, the results of different pipelines were treated as comparable. In most cases, PhiPipe showed comparable results with DPARSF/PANDA. PhiPipe showed better validity for ALFF than DPARSF, and better reliability and validity for SC than PANDA. More numerical details could be found in Supplementary Table S29-S30. However, we should be careful about the comparison results among different pipelines, as there were also processing variants in DPARSF and PANDA, which would influence the reliability and validity results. Also we only compared the overall reliability and validity between pipelines, and the parcel-wise or edge-wise differences were unpredictable.

**Figure 8.**
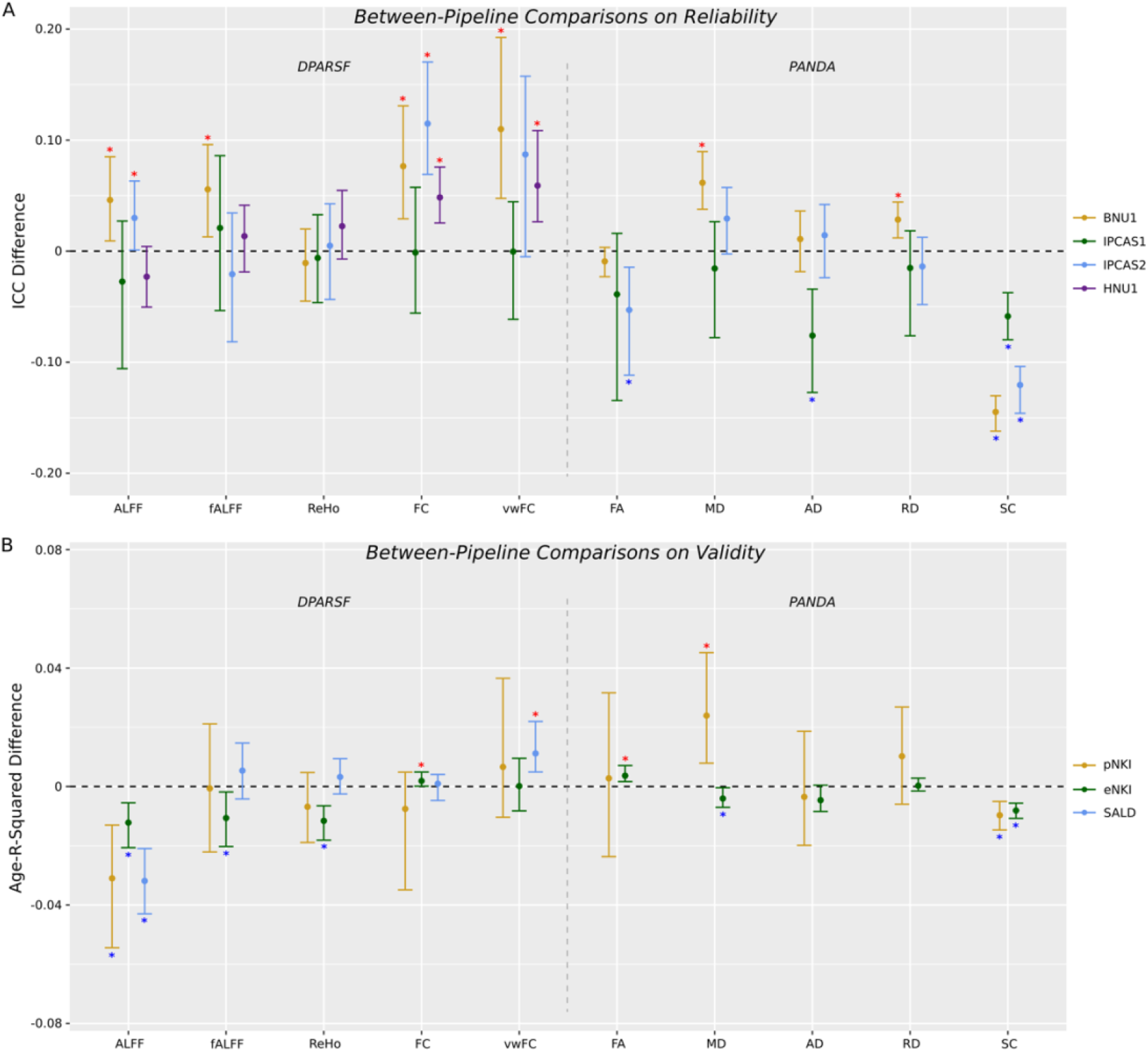
The comparison between PhiPipe and DPARSF/PANDA on the ICC and Age-R-Squared for resting-state BOLD and DWI brain features across all datasets. The Y axis means the difference of mean ICC or Age-R-Squared between the results using PhiPipe and DPARSF/PANDA. The data are presented as (DPARSF/PANDA - PhiPipe), so a positive value means increased reliability and validity using DPARSF/PANDA compared with PhiPipe. The error bars represent 95% CIs. If the CI does not contain zero, the difference is statistically significant. The red/blue asterisks are used to indicate significant increase or decrease.

### 3.4 The multivariate reliability and validity of brain features

Figure 9A presented the dbICC results for all brain features and datasets. Similar to the univariate reliability, T1 and DWI brain features showed higher reliability than the resting-state BOLD brain features. Figure 9B presented the CV-R results for all brain features and datasets. Different from the univariate validity results, all brain features except CA showed comparable multivariate predicative validity. More numerical details were presented in Supplementary Table S31-S32.

**Figure 9.**
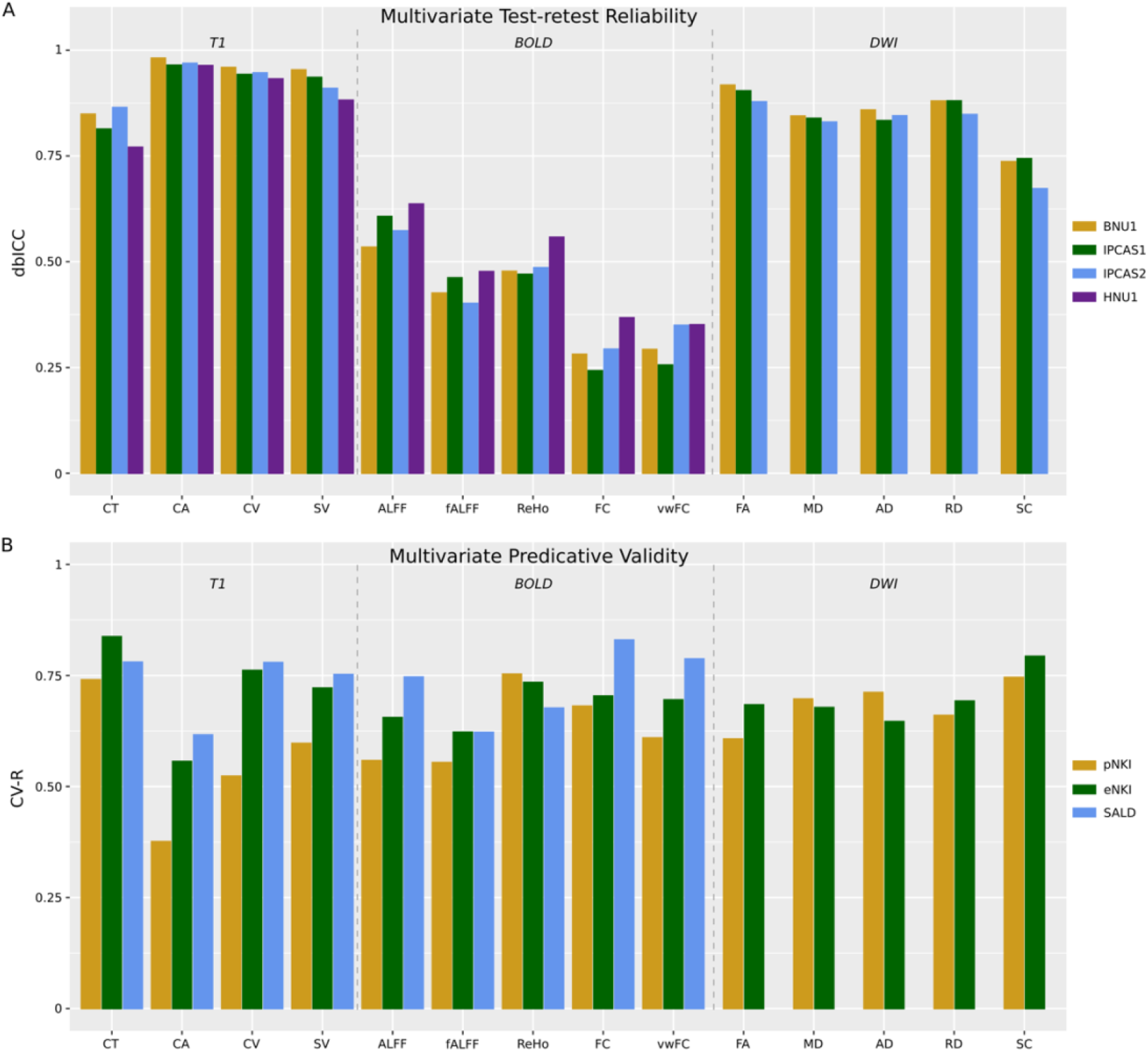
The dbICC and CV-R (the correlation between real age and predicated age) for all brain features across all datasets.

## 4. Discussions

We introduced PhiPipe for multi-modal MRI data processing. PhiPipe could generate common brain features for T1-weighted, resting-state BOLD and DWI data. We validated the PhiPipe with multiple public datasets from the perspectives of test-retest reliability and predicative validity. Compared with other pipelines, PhiPipe showed comparable or better reliability and validity.

The reliability and validity assessment of brain features could help researchers estimate sample size in experimental design or choose the appropriate brain features in statistical analysis. As an example, Figure 10 showed the effects of reliability and validity on the sample size needed to achieve an alpha level of 0.05 and a statistical power of 0.8. In Figure 10A, when we assumed the true correlation between a brain property and another variable of interest (for instance, memory function), the sample size needed was moderated by the reliability of the specific feature representing the brain property. We should note that the Figure 10A does not mean which brain feature is the best for a certain study, because different brain features represent different properties of human brain. In Figure 10B, we presented the relationship between sample size needed and observed Age-R. When we make an experiment design, we usually refer to previous studies to estimate the effect size. We could use the observed Age-R as an effect size estimate for future studies and consider whether the effect size of interest should be larger or smaller than the age effect. Again, the Figure 10B does not mean which brain feature is the best for a certain study. The sample size estimation was performed using *pwr* package (v1.3-0) of R. To make the access to the reliability and validity measures easier, an online database (https://yangzhi-psy.shinyapps.io/NeuroImageFeatureQualityViewer/) was provided so that researchers could query the reliability and validity measures for specific brain regions and image features.

**Figure 10.**
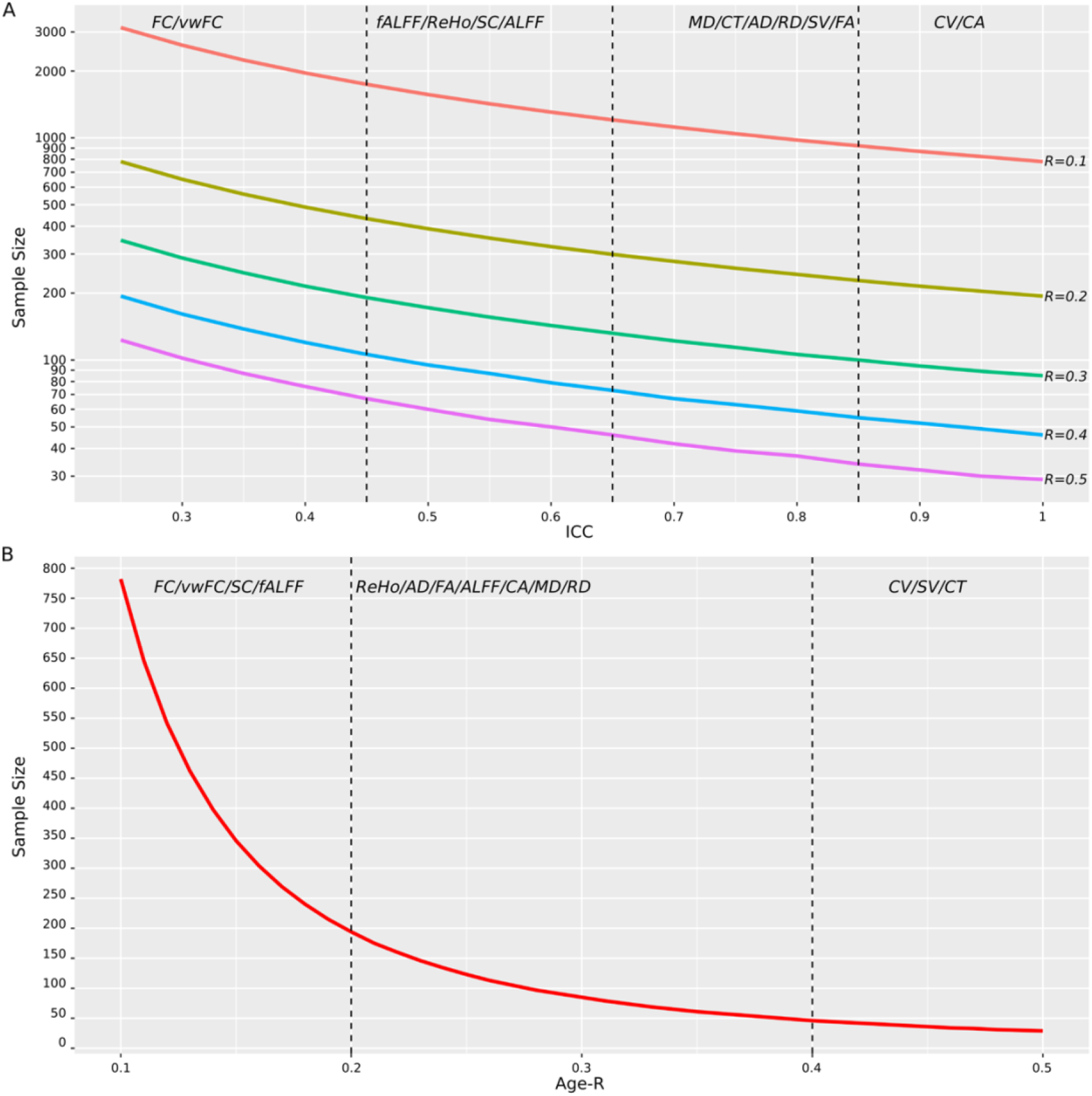
The relationship between ICC/Age-R and sample size needed to achieve an alpha level of 0.05 and a statistical power of 0.8. (A) For a specified true correlation between two given measures (e.g., brain feature and memory function), the sample size needed to detect this effect was moderated by the reliability of brain feature (here we simply assume the other measure is of perfect reliability). (B) The sample size needed to detect a specified observed Age-R. The brain features at the top of the panel were ordered by the average of ICC or Age-R across all datasets weighted by the sample size and scans.

PhiPipe provided an easy-to-use solution for multi-modal MRI data processing. Besides the three core MRI data processing softwares (i.e., FreeSurfer, AFNI and FSL) and Bash shell, other dependencies are minimum or optional, which make the installation and usage simple and robust in most computation environments. Qualitative and quantitative quality control pictures and measures were created to ensure the accuracy of processing results. The PhiPipe generates parcel-wise or edge-wise brain features characterizing the regional or inter-regional properties of human brain, which could be directly used for downstream statistical analysis. Future versions of PhiPipe would include more types of brain features after careful selection and testing.

PhiPipe is designed to be a simple-to-use image processing pipeline with validated results. This nature trades off the flexibility of processing steps and parameters for easy usage and confidence in the reliability and validity of the results. Compared with PhiPipe, there are many image processing pipelines featuring high flexibility. For instance, AFNI’s afni_proc.py allows users to combine processing blocks to arrange desired order of the processing steps. C-PAC provides a lot of options to implement parallel processing flows for fMRI data with little effort. These pipelines are suitable for users who need a high degree of freedom for processing steps and parameters. In contrast, PhiPipe is more suitable for researchers who need an imaging processing tool that is simple to use and extracts multimodal brain features with expectable reliability and validity.

In PhiPipe, we introduced CAT12 for skull-stripping and TIV estimation. For skull-stripping, CAT12 led to worse validity than FreeSurfer in the current datasets. For TIV estimation, CAT12 showed higher reliability and validity. For BOLD brain features, only the temporal filtering had the consistent and biggest influences on the reliability and validity. For the disputed global signal regression, the effect was not consistent. We should note that the results of processing variants do not mean which processing variant is the best, but should be used as estimates of potential reliability and validity changes when we choose to use a different processing step. We observed apparent dataset-dependent variability in most results, which indicate that we must carefully choose the datasets in any comparisons and validations.

This work also presents a general framework for evaluating reliability and validity of image processing pipelines. In recent years, there has been many concerns about the reproducibility of neuroimaging studies (Masouleh et al., 2019; Noble et al., 2019; Botvinik-Nezer et al., 2020; Elliott et al., 2020; Marek et al., 2022). Some researchers proposed that the standardization of pipelines could improve the reproducibility across studies, because using different pipelines would lead to different results and conclusions. In reality, almost all labs have their own custom pipelines, whether the pipelines are private or publicly available. We believe that the existing variability of processing pipelines was due to the fact that the brain features commonly used were not reliable or valid. For instance, the brain features of resting-state BOLD data had poor reliability and consequently low validity. As a result, various processing steps were adopted in different studies. Even for the cortical thickness measure, its reliability was far from perfect and the validity has not been fully established (Cardinale et al., 2014). Therefore, we propose that, instead of standardization of pipelines, improving the reliability and validity of brain features is the more urgent problem to solve in this field. Along the same line, we found the brain features from PhiPipe and two other pipelines showed similar reliability and validity. In other words, the low reliability and validity of MRI brain features is probably common to all pipelines, as all pipelines almost relies on the same sets of atomic softwares. We also share the core code and detailed information about the public datasets to evaluate the reliability and validity of pipelines, and hopefully more evaluations of commonly used image processing pipelines will emerge.

Most quality control procedures for key processing results relied on visual check in PhiPipe, which were the same in other pipelines. Visual check is time-consuming and subjective-biased, and how to perform quantitative quality control is the future direction of PhiPipe. The current version of PhiPipe was T1-centered and the results of T1 were used for BOLD/DWI processing. However, this common practice does not exploit the full possibilities of multi-modal MRI data. How to use BOLD/DWI data to optimize the accuracy of T1 processing results is another goal in future development of PhiPipe.

## 5. Conclusion

We presented the PhiPipe to facilitate multi-modal MRI data processing. The accompanying test-retest reliability and predicative validity assessment could help researchers make informed decisions in conducting experimental design and statistical analysis.

## Acknowledgements

This work was supported by the National Key R&D Program of China (2018YFC2001600); National Natural Science Foundation of China (81971682, 81571756, 81270023); Natural Science Foundation of Shanghai (20ZR1472800); Shanghai Municipal Commission of Education-Gaofeng Clinical Medicine Grant Support (20171929); Hundred-Talent Fund from Shanghai Municipal Commission of Health (2018BR17); Shanghai Mental Health Center Clinical Research Center (CRC2018DSJ01-5; CRC2019ZD04); Research Funds from Shanghai Mental Health Center (13dz2260500)

## Data Availability Statement

All datasets used in this study were publicly available and detailed information could be found in Supplementary Materials.

