## Supplementary Materials for "PhiPipe: a multi-modal MRI data processing pipeline with test-retest reliability and predicative validity assessments"

1. The demographic characteristics and subject IDs of public datasets used in the study.

The BNU1 dataset (http://fcon_1000.projects.nitrc.org/indi/CoRR/html/bnu_1.html) included 57 heathy subjects (male/female: 30/27, age range: 19-30 years) who completed two scans at an average interval of 41 days (range: 33-55 days). After quality control, 45 subjects were involved in the final analysis.

The IPCAS1 dataset (http://fcon_1000.projects.nitrc.org/indi/CoRR/html/ipcas_1.html) included 30 subjects (male/female: 9/21, age range: 18-24) who completed two scans at an average interval of 14 days (range: 5-24 days). After quality control, 18 subjects were involved in the final analysis.

The IPCAS2 dataset (http://fcon_1000.projects.nitrc.org/indi/CoRR/html/ipcas_2.html) included 35 healthy subjects (male/female: 12/23, age range: 11-15) who completed two scans at an average interval of 33 days (range: 7-59 days). After quality control, 27 subjects were involved in the final analysis.

The HNU1 dataset (http://fcon_1000.projects.nitrc.org/indi/CoRR/html/hnu_1.html) included 30 healthy subjects (male/female: 15/15, age range: 20-30) who completed 10 scans at an average interval of 3 days (range: 2-10 days). After quality control, 22 subjects were involved in the final analysis.

The pNKI dataset (http://fcon_1000.projects.nitrc.org/indi/pro/nki.html) included 207 subjects (male/female: 120/87, age range: 4-85). In this study, only adult subjects (age >=18, 151 subjects) were included for data analysis. After quality control, 113 subjects were involved in the final analysis. The age distribution of subjects involved in this study was shown below:

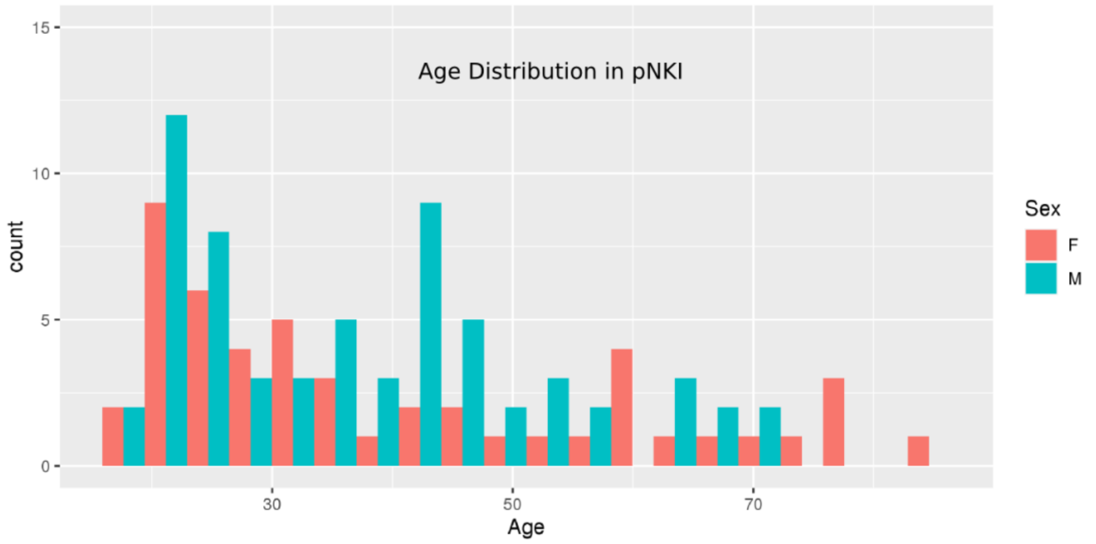


The eNKI dataset (http://fcon_1000.projects.nitrc.org/indi/enhanced/neurodata.html) included 1305 subjects (male/female: 791/514, age range: 6-85). Each subject had three resting-state BOLD fMRI scans with different scanning parameters (two with multi-band and one with single-band), and the single-band BOLD data was used in this study. In this study, only adult subjects (age >=18, 997 subjects) were included for data analysis. After quality control, 764 subjects were eligible for further analysis. To match the age distribution between male and female subjects, 502 subjects were selected for the final analysis. The age distribution of subjects involved in this study was shown below:

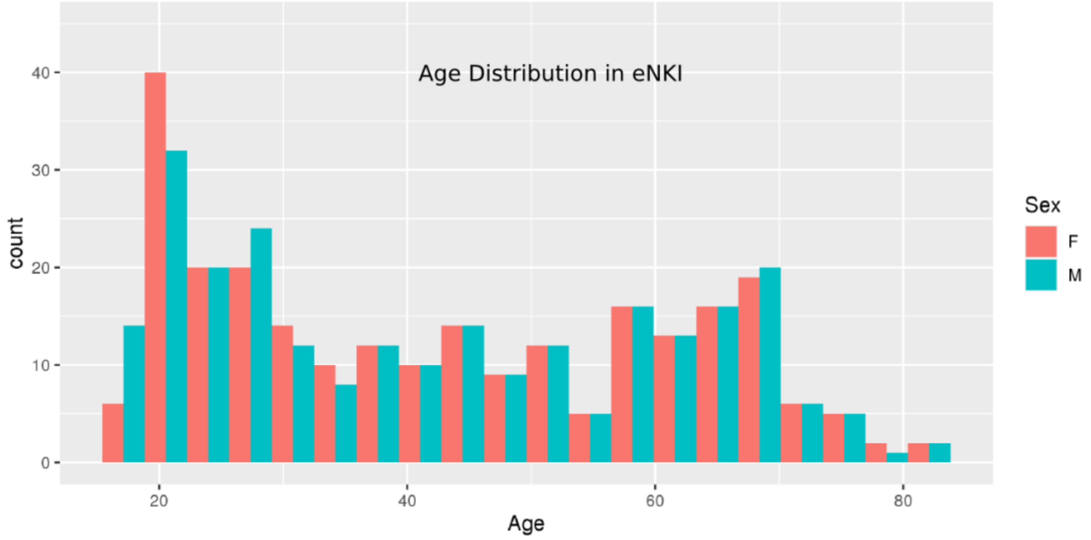


The SALD dataset (http://fcon_1000.projects.nitrc.org/indi/retro/sald.html) included 494 subjects (male/female: 187/307, age range: 19-80 years). After quality control, 414 subjects were included in the final analysis. The age distribution of subjects involved in this study was shown below:

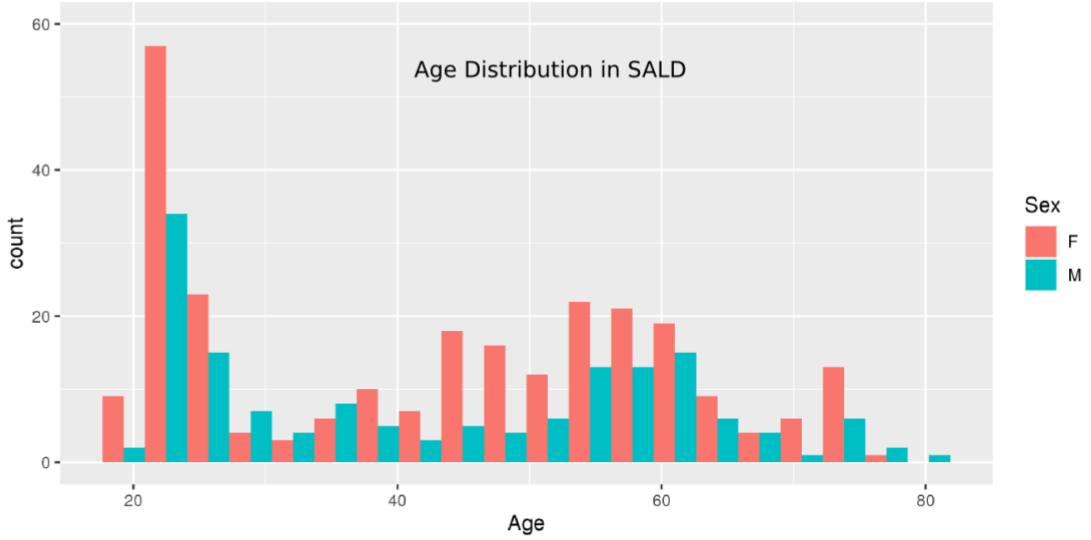


The IDs of finally involved subjects for all datasets were shown below:

*BNU1*

0025864 0025865 0025866 0025867 0025868 0025869 0025870 0025871 0025872 0025873 0025874 0025876 0025877 0025878 0025879 0025880 0025881 0025882 0025883 0025884

0025886 0025887 0025888 0025889 0025891 0025892 0025893 0025894 0025895 0025896 0025897 0025898 0025899 0025900 0025901 0025902 0025903 0025904 0025905 0025906 0025908 0025909 0025910 0025911 0025912

*IPCAS1*

0025485 0025487 0025488 0025490 0025492 0025493 0025495 0025496 0025497 0025499 0025501 0025502 0025504 0025505 0025508 0025509 0025510 0025514

*IPCAS2*

0025515 0025516 0025517 0025518 0025519 0025520 0025522 0025523 0025524 0025525 0025527 0025528 0025530 0025531 0025532 0025533 0025534 0025536 0025537 0025538 0025539 0025540 0025542 0025543 0025546 0025548 0025549

*HNU1*

0025427 0025428 0025429 0025432 0025435 0025436 0025437 0025438 0025439 0025440 0025443 0025444 0025445 0025446 0025447 0025448 0025449 0025450 0025451 0025452 0025453 0025456

*pNKI*

1013090 1097782 1150497 1216620 1264721 1271401 1285465 1322220 1334556 1339484 1366839 1382333 1427581 1508861 1522653 1709141 1713513 1734822 1834799 1903493 1915832 1931386 1935851 1961098 2009256 2021454 2071045 2132437 2139745 2160826 2166659 2176073 2221971 2258252 2286053 2286816 2328270 2357976 2362594 2436415 2457957 2475376 2483617 2505567 2513310 2524225 2531270 2541244 2589215 2631208 2731081 2788776 2799329 2842950 2861923 2986140 3033715 3070385 3209959 3304648 3313349 3315657 3329569 3366302 3409284 3431940 3466763 3486362 3606220 3611212 3695138 3701005 3729976 3755111 3804254 3808535 3811036 3815065 3827479 3911767 3927656 3934325 3951328 3965194 4015919 4077433 4119751 4126393 4154799 4188346 4277600 4288245 4290056 4323037 4487215 4791943 5205221 5279162 5461463 6471972 6539040 6709880 6913939 7055197 8317157 8574662 8691891 8735778 9006154 9100911 9536886 9630905 9645370

*eNKI*

00008326 00013809 00019903 00027544 00027651 00028150 00028152 00028184 00028185 00028192 00028207 00028246 00028287 00028352 00028389 00028399 00028400 00028429 00028552 00028606 00028625 00028656 00028678 00028691 00028753 00028754 00028766 00028784 00028822 00028842 00028844 00028845 00028929 00028994 00029075 00029076 00029092 00029104 00029126 00029127 00029219 00029231 00029303 00029304 00029979 00030912 00030913 00030947 00030989 00030990 00031167 00031216 00031217 00031452

00031459 00031549 00031605 00031625 00031794 00031871 00031872 00031893 00031894
00032007 00032010 00032862 00032875 00032876 00033011 00033180 00033231 00033232
00033247 00033248 00033589 00033609 00033640 00033643 00033647 00033747 00033748
00033834 00033849 00033903 00033963 00033998 00034093 00034193 00034385 00034400
00034827 00034854 00034987 00035072 00035440 00035504 00035535 00035553 00035562
00035606 00035608 00035625 00035699 00035744 00035765 00035798 00035827 00035840
00035881 00035926 00035943 00035960 00037081 00037110 00037117 00037118 00037122
00037160 00037170 00037266 00037377 00037378 00037386 00037387 00037396 00037439
00037445 00037457 00037476 00037492 00037510 00037511 00037522 00037531 00037545
00037582 00037588 00037635 00037646 00037647 00037725 00037783 00037784 00037817
00037866 00037877 00038159 00038189 00038285 00038411 00038424 00038522 00038605
00038616 00038623 00038731 00038831 00038832 00038957 00038998 00039074 00039084
00039143 00039159 00039257 00039391 00039431 00039432 00039461 00039463 00039490
00039500 00039559 00039593 00039607 00039635 00039636 00039639 00039640 00039655
00039657 00039669 00039686 00039699 00039700 00039755 00039758 00039783 00039819
00039820 00039895 00039916 00039951 00039952 00039966 00039988 00040116 00040151
00040152 00040225 00040226 00040257 00040285 00040286 00040301 00040324 00040351
00040462 00040493 00040494 00040500 00040517 00040524 00040525 00040543 00040573
00040594 00040599 00040604 00040623 00040628 00040629 00040678 00040741 00040806
00040827 00040890 00040891 00040915 00043283 00043299 00043384 00043417 00043450
00043462 00043466 00043520 00043635 00043649 00043677 00043704 00043721 00043722
00043998 00044012 00044130 00044131 00044171 00044291 00044344 00044408 00045554
00045590 00050720 00050795 00050911 00051063 00051064 00051455 00051457 00051477
00051513 00051514 00051528 00051529 00051539 00051603 00051604 00051638 00051676
00051679 00051726 00051774 00051927 00052070 00052117 00052118 00052125 00052126
00052181 00052197 00052235 00052307 00052340 00052500 00052502 00053202 00053455
00053473 00053474 00053475 00053546 00053576 00053577 00053578 00053850 00053851
00053854 00053901 00053927 00053949 00054019 00054153 00054173 00054441 00054482
00054532 00054533 00054534 00054621 00054857 00055121 00055215 00055373 00055446
00055447 00055542 00055763 00055806 00056097 00056164 00056306 00056627 00056898
00056919 00056949 00057005 00057183 00057372 00057444 00057786 00057965 00058214
00058503 00058644 00058667 00058952 00058998 00059361 00059845 00059911 00059990
00060006 00060093 00060169 00060170 00060279 00060302 00060372 00060383 00060407
00060431 00060471 00060480 00060516 00060574 00060576 00060581 00060582 00060602
00060630 00060632 00060773 00060923 00060925 00061204 00061210 00061274 00061275
00061284 00061387 00061598 00061600 00061656 00061666 00061711 00061725 00061727
00061767 00061881 00062248 00062288 00062351 00062411 00062923 00062929 00062934
00062937 00062982 00063103 00063326 00063450 00063455 00063458 00063473 00063589
00064058 00064415 00065197 00065574 00065717 00065722 00065749 00065995 00066087
00066091 00066153 00066232 00066236 00066246 00066282 00066389 00066539 00066735
00066781 00066812 00066926 00073296 00073600 00073611 00073953 00074000 00074425
00074609 00074687 00074711 00074793 00074802 00074836 00074956 00074985 00075220
00075267 00075407 00075674 00075757 00075964 00076381 00076798 00077091 00077099
00077134 00077181 00077204 00077252 00077391 00077622 00077637 00077758 00078262

00079302 00079353 00079422 00079724 00079730 00079747 00079777 00079850 00080052 00080447 00080535 00080819 00081071 00081098 00081112 00081132 00081142 00081149 00081185 00081243 00081253 00081402 00081544 00081616 00081729 00081735 00081785 00081846 00082215 00082532 00082554 00082556 00082619 00082804 00082942 00083109 00083138 00083171 00083419 00083631 00083923 00083982 00083983 00084921 00085057 00085095 00085391 00085680 00085828 00085907 00086748 00088338

*SALD*

031274 031275 031276 031277 031278 031279 031280 031281 031282 031283 031285 031287 031288 031289 031290 031291 031292 031295 031296 031297 031298 031299 031301 031302 031303 031305 031306 031307 031308 031309 031310 031311 031312 031313 031314 031315 031316 031317 031321 031322 031323 031324 031325 031326 031327 031328 031330 031331 031332 031333 031336 031337 031338 031339 031340 031342 031343 031344 031345 031349 031350 031351 031352 031353 031354 031357 031358 031359 031360 031361 031363 031365 031366 031367 031368 031370 031371 031372 031374 031375 031376 031377 031379 031380 031381 031382 031383 031384 031385 031386 031387 031388 031389 031390 031391 031392 031393 031398 031399 031400 031402 031403 031404 031407 031409 031410 031411 031412 031414 031415 031416 031419 031420 031422 031423 031424 031426 031427 031428 031431 031432 031434 031436 031437 031439 031440 031441 031442 031443 031444 031445 031446 031447 031449 031450 031451 031453 031454 031457 031458 031459 031462 031463 031464 031465 031466 031467 031468 031469 031470 031471 031472 031473 031474 031475 031476 031477 031478 031479 031480 031481 031482 031484 031485 031486 031487 031489 031490 031491 031492 031493 031494 031495 031496 031497 031498 031499 031500 031501 031502 031504 031505 031506 031507 031508 031509 031510 031511 031513 031514 031515 031516 031517 031518 031519 031520 031522 031523 031524 031525 031526 031527 031528 031529 031530 031531 031533 031534 031535 031537 031538 031539 031540 031541 031542 031543 031544 031545 031546 031547 031548 031549 031551 031552 031553 031554 031555 031556 031558 031559 031560 031561 031562 031563 031566 031567 031568 031569 031570 031571 031572 031573 031574 031575 031576 031579 031580 031581 031582 031583 031585 031587 031588 031589 031590 031591 031592 031593 031594 031595 031596 031597 031598 031599 031600 031601 031602 031603 031604 031605 031606 031607 031609 031610 031611 031612 031613 031614 031615 031617 031618 031619 031620 031621 031622 031623 031624 031625 031626 031627 031628 031629 031630 031631 031632 031634 031635 031636 031637 031638 031641 031643 031644 031645 031646 031647 031650 031651 031652 031653 031654 031655 031656 031657 031658 031659 031660 031662 031663 031664 031665 031666 031667 031668 031669 031671 031672 031673 031674 031675 031676 031677 031678 031679 031680 031681 031682 031683 031684 031685 031686 031687 031689 031691 031692 031693 031694 031695 031696 031697

031698 031699 031700 031701 031702 031703 031704 031706 031710 031711
031712 031713 031714 031715 031716 031718 031720 031721 031722 031723
031724 031725 031726 031727 031728 031729 031730 031731 031732 031733
031734 031735 031736 031737 031738 031739 031740 031741 031742 031743
031744 031745 031746 031747 031748 031749 031750 031751 031752 031753
031754 031755 031756 031757 031758 031759 031760 031761 031762 031763
031764 031765 031766 031767

2. The codes to invoke PhiPipe to process MRI data (using the BNU1 dataset as an example)

```
## make the PhiPipe globally accessible
export PhiPipe=/home/huyang/PhiPipe_v1.2
## T1 processing
## ${T1} means the T1 raw data with absolute path
## ${OUTDIR} means the folder to store the T1 processing results
bash ${PhiPipe}/t1_process.sh -a ${T1} -b ${OUTDIR} -c t1
## BOLD processing
## ${BOLD} means the BOLD raw data with absolute path
bash ${PhiPipe}/bold_process.sh -a ${OUTDIR} -b t1 -c ${BOLD} -d ${OUTDIR} -e bold -f 5 -h 2 -i 3 -
k 0.01 -l 0.1
## DWI processing
## ${DWI} means the DWI raw data with absolute path
## ${BVAL}/${BVEC} means the b-value and b-vector files with absolute paths
bash ${PhiPipe}/dwi_process.sh -a ${OUTDIR} -b t1 -c ${DWI} -d ${OUTDIR} -e dwi -f ${BVAL} -g
${BVEC} -h 2 -i 1
```

3. The data processing and quality control procedures using DPARSF

Resting-state BOLD fMRI data were processed using default parameters (except *Image Reorient*) of DPARSF. The major processing steps included: (1) removing the first 10 volumes; (2) slice timing correction; (3) motion correction; (4) nuisance regression with mean WM/CSF signals, Friston's 24-parameter motion model and linear trends as nuisance covariates; (5) spatial normalization to the sample-specific template (in MNI space) created by DARTEL; (6) bandpass filtering at 0.01-0.1 Hz. By default, DPARSF requires user to manually reorient the image origin into the anterior commissure. As this step is time-consuming and has little influence on the final results (based on our experiences), we skipped this step.

The fALFF/ALFF/ReHo were calculated and Z-scored in DPARSF. The mean fALFF/ALFF/ReHo were extracted based on the Schaefer+Aseg Atlases and compared with PhiPipe's results. The mean time series were extracted from the DPARSF's output based on the Schaefer+Aseg Atlases, and the FC/vwFC matrices were computed and compared with PhiPipe's results.

The spatial normalization pictures were visually checked and no subjects were excluded due to bad registrations. No other quality control pictures were available (as far as we know).

The full parameters (excluding SubjectID variable) of DPARSF were shown below (using BNU dataset as an example):

```
Cfg.DPARSFVersion = 'V5.2_210501';
Cfg.WorkingDir = ['/home/huyang/BNU1/PROCDATA/REST_DPABI'];
Cfg.DataProcessDir = ['/home/huyang/BNU1/PROCDATA/REST_DPABI'];
Cfg.TimePoints = 200;
Cfg.TR = 2;
Cfg.IsNeedConvertFunDCM2IMG = 0;
Cfg.IsRemoveFirstTimePoints = 1;
Cfg.RemoveFirstTimePoints = 10;
Cfg.IsSliceTiming = 1;
Cfg.SliceTiming = struct;
Cfg.SliceTiming.SliceNumber = 33;
Cfg.SliceTiming.SliceOrder = [1 3 5 7 9 11 13 15 17 19 21 23 25 27 29 31 33 2 4 6 8 10 12 14 16 18
20 22 24 26 28 30 32];
Cfg.SliceTiming.ReferenceSlice = 33;
Cfg.IsRealign = 1;
Cfg.IsCalVoxelSpecificHeadMotion = 0;
Cfg.IsNeedReorientFunImgInteractively = 0;
Cfg.IsNeedConvertT1DCM2IMG = 0;
Cfg.IsNeedReorientCropT1Img = 0;
Cfg.IsNeedReorientT1ImgInteractively = 0;
Cfg.IsNeedT1CoregisterToFun = 1;
Cfg.IsNeedReorientInteractivelyAfterCoreg = 0;
```

```
Cfg.IsSegment = 2;
Cfg.Segment = struct;
Cfg.Segment.AffineRegularisationInSegmentation = 'mni';
Cfg.IsDARTEL = 1;
Cfg.IsCovremove = 1;
Cfg.Covremove = struct;
Cfg.Covremove.Timing = 'AfterRealign';
Cfg.Covremove.PolynomialTrend = 1;
Cfg.Covremove.HeadMotion = 4;
Cfg.Covremove.IsHeadMotionScrubbingRegressors = 0;
Cfg.Covremove.HeadMotionScrubbingRegressors = struct;
Cfg.Covremove.HeadMotionScrubbingRegressors.FDThreshold = 0.2;
Cfg.Covremove.HeadMotionScrubbingRegressors.PreviousPoints = 0;
Cfg.Covremove.HeadMotionScrubbingRegressors.LaterPoints = 0;
Cfg.Covremove.HeadMotionScrubbingRegressors.FDType = 'FD_Jenkinson';
Cfg.Covremove.WholeBrain = struct;
Cfg.Covremove.WholeBrain.IsRemove = 0;
Cfg.Covremove.WholeBrain.Mask = 'SPM';
Cfg.Covremove.WholeBrain.Method = 'Mean';
Cfg.Covremove.WholeBrain.IsBothWithWithoutGSR = 0;
Cfg.Covremove.CSF = struct;
Cfg.Covremove.CSF.IsRemove = 1;
Cfg.Covremove.CSF.Mask = 'SPM';
Cfg.Covremove.CSF.MaskThreshold = 0.99;
Cfg.Covremove.CSF.Method = 'Mean';
Cfg.Covremove.CSF.CompCorPCNum = 5;
Cfg.Covremove.OtherCovariatesROI = [];
Cfg.Covremove.WM = struct;
Cfg.Covremove.WM.IsRemove = 1;
Cfg.Covremove.WM.Mask = 'SPM';
Cfg.Covremove.WM.MaskThreshold = 0.99;
Cfg.Covremove.WM.Method = 'Mean';
Cfg.Covremove.WM.CompCorPCNum = 5;
Cfg.Covremove.IsAddMeanBack = 0;
Cfg.IsFilter = 1;
Cfg.Filter = struct;
Cfg.Filter.Timing = 'AfterNormalize';
Cfg.Filter.ALowPass_HighCutoff = 0.1;
Cfg.Filter.AHighPass_LowCutoff = 0.01;
Cfg.Filter.AAddMeanBack = 'Yes';
Cfg.IsNormalize = 3;
Cfg.Normalize = struct;
Cfg.Normalize.Timing = 'OnFunctionalData';
Cfg.Normalize.BoundingBox = [-90 -126 -72;90 90 108];
```

```
Cfg.Normalize.VoxSize = [3 3 3];
Cfg.IsSmooth = 0;
Cfg.Smooth = struct;
Cfg.Smooth.Timing = 'OnResults';
Cfg.Smooth.FWHM = [4 4 4];
Cfg.MaskFile = 'Default';
Cfg.IsWarpMasksIntoIndividualSpace = 0;
Cfg.IsDetrend = 0;
Cfg.IsCalALFF = 1;
Cfg.CalALFF = struct;
Cfg.CalALFF.AHighPass_LowCutoff = 0.01;
Cfg.CalALFF.ALowPass_HighCutoff = 0.1;
Cfg.IsScrubbing = 0;
Cfg.Scrubbing = struct;
Cfg.Scrubbing.Timing = 'AfterPreprocessing';
Cfg.Scrubbing.FDThreshold = 0.2;
Cfg.Scrubbing.PreviousPoints = 0;
Cfg.Scrubbing.LaterPoints = 0;
Cfg.Scrubbing.ScrubbingMethod = 'cut';
Cfg.Scrubbing.FDType = 'FD_Jenkinson';
Cfg.IsCalReHo = 1;
Cfg.CalReHo = struct;
Cfg.CalReHo.ClusterNVoxel = 27;
Cfg.CalReHo.SmoothReHo = 0;
Cfg.IsCalDegreeCentrality = 0;
Cfg.CalDegreeCentrality = struct;
Cfg.CalDegreeCentrality.rThreshold = 0.25;
Cfg.IsCalFC = 0;
Cfg.IsExtractROISignals = 0;
Cfg.CalFC = struct;
Cfg.CalFC.ROIDef = cell(10, 1);
Cfg.CalFC.ROIDef{1,1}
=['/home/huyang/matlab/DPABI_V6.0/Templates/aal.nii'];
Cfg.CalFC.ROIDef{2,1}
=['/home/huyang/matlab/DPABI_V6.0/Templates/HarvardOxford-cort-maxprob-thr25-
2mm_YCG.nii'];
Cfg.CalFC.ROIDef{3,1}
=['/home/huyang/matlab/DPABI_V6.0/Templates/HarvardOxford-sub-maxprob-thr25-
2mm_YCG.nii'];
Cfg.CalFC.ROIDef{4,1}
=['/home/huyang/matlab/DPABI_V6.0/Templates/CC200ROI_tcorr05_2level_all.nii'];
Cfg.CalFC.ROIDef{5,1}
=['/home/huyang/matlab/DPABI_V6.0/Templates/Zalesky_980_parcellated_compact.nii'];
Cfg.CalFC.ROIDef{6,1}
```

```
=['/home/huyang/matlab/DPABI_V6.0/Templates/Dosenbach_Science_160ROIs_Radius5_Mask.n
ii'];
Cfg.CalFC.ROIDef{7,1}
=['/home/huyang/matlab/DPABI_V6.0/Templates/BrainMask_05_91x109x91.img'];
Cfg.CalFC.ROIDef{8,1}
=['/home/huyang/matlab/DPABI_V6.0/Templates/Power_Neuron_264ROIs_Radius5_Mask.nii'];
Cfg.CalFC.ROIDef{9,1}
=['/home/huyang/matlab/DPABI_V6.0/Templates/Schaefer2018_400Parcels_7Networks_order_F
SLMNI152_1mm.nii'];
Cfg.CalFC.ROIDef{10,1}
=['/home/huyang/matlab/DPABI_V6.0/Templates/Tian2020_Subcortex_Atlas/Tian_Subcortex_S4
_3T.nii'];
Cfg.CalFC.IsMultipleLabel = 1;
Cfg.IsDefineROIInteractively = 0;
Cfg.IsExtractAALTC = 0;
Cfg.IsCalVMHC = 0;
Cfg.IsCWAS = 0;
Cfg.CWAS = struct;
Cfg.CWAS.Regressors = [];
Cfg.CWAS.iter = 0;
Cfg.FunctionalSessionNumber = 1;
Cfg.StartingDirName = 'FunImg';
Cfg.ParallelWorkersNumber = 10;
Cfg.IsNormalizeToSymmetricGroupT1Mean = 0;
Cfg.IsApplyDownloadedReorientMats = 0;
Cfg.IsBet = 1;
Cfg.IsAutoMask = 1;
Cfg.IsSmoothBeforeVMHC = 0;
Cfg.IsBIDStoDPARSF = 0;
```

4. The data processing and quality control procedures using PANDA

DWI data were processed using default parameters of PANDA. The major processing steps included: (1) cropping the image to save space; (2) eddy/motion correction; (3) diffusion tensor fitting to calculate FA/MD/AD/RD measures; (4) register FA maps into the FA58_FMRIB template; (5) extract the mean FA/MD/AD/RD based on the JHU Tract Atlas, which were compared with PhiPipe's results; (6) register the FA image into the T1 brain image (after skull-stripping); (7) register T1 brain image into the MNI152 template and transform the Schaefer+Aseg Atlases into the individual's native space; (8) probabilistic tractography based on the Schaefer+Aseg Atlases and the SC matrix was compared with PhiPipe's results.

The FA-FA58_FMRIB registration, T1 skull-stripping, FA-T1 registration, T1-MNI152 registration and parcellation results at the native space were visually checked. Two subjects in the IPCAS1 dataset were excluded due to bad T1-MNI152 registration results. Two subjects in the IPCAS2 datasets were excluded due to bad T1-MNI152 registration results. No subjects were excluded in other datasets. As T1-MNI152 registration only had influence on the SC matrix, so when compare PANDA with PhiPipe on the SC matrix, the excluded subjects in PANDA were also removed in PhiPipe.

The full parameters (excluding the data paths) of PANDA were shown below (using the BNU1 dataset as an example):

```
SubjectIDArray = [1 2 3 4 5 6 7 8 9 10 11 12 13 14 15 16 17 18 19 20 21 22 23 24 25 26 27 28 29
30 31 32 33 34 35 36 37 38 39 40 41 42 43 44 45 46 47 48 49 50 51 52 53 54 55 56 57 58 59 60 61
62 63 64 65 66 67 68 69 70 71 72 73 74 75 76 77 78 79 80 81 82 83 84 85 86 87 88 89 90];
DestinationPath_Edit
= ['/home/huyang/BNU1/PROCDATA/DWI_PANDA/OUTPUT'];
TensorPrefixEdit = 'dwi';
pipeline_opt = struct;
pipeline_opt.mode = 'qsub';
pipeline_opt.max_queued = 40;
pipeline_opt.flag_verbose = 0;
pipeline_opt.flag_pause = 0;
pipeline_opt.qsub_options = '-cwd -V -q new.q';
dti_opt = struct;
dti_opt.BET_1_f = 0.25;
dti_opt.BET_2_f = 0.25;
dti_opt.Delete_Flag = 0;
dti_opt.Cropping_Flag = 1;
dti_opt.NIIcrop_slice_gap = 3;
dti_opt.RawDataResample_Flag = 0;
dti_opt.RawDataResampleResolution = [2 2 2];
dti_opt.Inversion = 'No Inversion';
dti_opt.Swap = 'No Swap';
```

```
dti_opt.LDH_Flag = 0;
dti_opt.LDH_Neighborhood = 7;
dti_opt.Normalizing_Flag = 1;
dti_opt.FAnormalize_target =
['/home/huyang/matlab/PANDA_1.3.1_64/data/Templates/FMRIB58_FA_1mm.nii.gz'];
dti_opt.applywarp_1_ref_fileName = 1;
dti_opt.applywarp_3_ref_fileName = 1;
dti_opt.applywarp_5_ref_fileName = 1;
dti_opt.applywarp_7_ref_fileName = 1;
dti_opt.Resampling_Flag = 1;
dti_opt.applywarp_2_ref_fileName = 2;
dti_opt.applywarp_4_ref_fileName = 2;
dti_opt.applywarp_6_ref_fileName = 2;
dti_opt.applywarp_8_ref_fileName = 2;
dti_opt.Smoothing_Flag = 1;
dti_opt.smoothNII_1_kernel_size = 6;
dti_opt.smoothNII_2_kernel_size = 6;
dti_opt.smoothNII_3_kernel_size = 6;
dti_opt.smoothNII_4_kernel_size = 6;
dti_opt.Atlas_Flag = 1;
dti_opt.WM_Label_Atlas =
['/home/huyang/matlab/PANDA_1.3.1_64/data/atlases/rICBM_DTI/rICBM_DTI_81_WMPM_FMR
IB58.nii.gz'];
dti_opt.WM_Probtract_Atlas =
['/home/huyang/matlab/PANDA_1.3.1_64/data/atlases/rICBM_DTI/JHU_ICBM_tracts_maxprob_
thr25_1mm.nii.gz'];
dti_opt.TBSS_Flag = 0;
tracking_opt = struct;
tracking_opt.DterminFiberTracking = 0;
tracking_opt.NetworkNode = 1;
tracking_opt.PartitionOfSubjects = 0;
tracking_opt.T1 = 1;
tracking_opt.DeterministicNetwork = 0;
tracking_opt.BedpostxProbabilisticNetwork = 1;
tracking_opt.ImageOrientation = 'Auto';
tracking_opt.Inversion = 'Invert Z';
tracking_opt.Swap = 'No Swap';
tracking_opt.PartitionTemplate =
['/home/huyang/Atlases/SchaeferAseg_MNI2mm.nii.gz'];
tracking_opt.T1Template =
['/home/huyang/matlab/PANDA_1.3.1_64/data/Templates/MNI152_T1_2mm_brain'];
tracking_opt.T1Bet_Flag = 1;
tracking_opt.T1BetF = 0.5;
tracking_opt.T1Cropping_Flag = 1;
```

tracking_opt.T1CroppingGap = 3;
tracking_opt.T1Resample_Flag = 1;
tracking_opt.T1ResampleResolution = [1 1 1];
tracking_opt.Fibers = 2;
tracking_opt.Weight = 1;
tracking_opt.Burnin = 1000;
tracking_opt.ProbabilisticTrackingType = 'OPD';
tracking_opt.LabelIdVector = [10 11 12 13 17 18 26 49 50 51 52 53 54 58 1001 1002 1003 1004 1005 1006 1007 1008 1009 1010 1011 1012 1013 1014 1015 1016 1017 1018 1019 1020 1021 1022 1023 1024 1025 1026 1027 1028 1029 1030 1031 1032 1033 1034 1035 1036 1037 1038 1039 1040 1041 1042 1043 1044 1045 1046 1047 1048 1049 1050 2001 2002 2003 2004 2005 2006 2007 2008 2009 2010 2011 2012 2013 2014 2015 2016 2017 2018 2019 2020 2021 2022 2023 2024 2025 2026 2027 2028 2029 2030 2031 2032 2033 2034 2035 2036 2037 2038 2039 2040 2041 2042 2043 2044 2045 2046 2047 2048 2049 2050];
tracking_opt.LabelIdVectorText = '[10 11 12 13 17 18 26 49:54 58 1001:1050 2001:2050]';
PANDAPath = '/home/huyang/matlab/PANDA_1.3.1_64';

5. The codes to perform ICC, bootstrapping and Bca confidence interval calculation

```
## ICC calculation
## The dat variable contains the brain feature of a parcel or edge, in which each row represents a subject and each column represents the scan session
library(irr)
stats <- icc(dat, model='two way', type='agreement', unit='single')
ICC <- stats$value

## bootstrap resampling
## NBOOT means the number of resampling
## NSUB means the number of subjects
set.seed(100)
NBOOT <- 2000
sub_idx <- c(1:NSUB)
boot_samples <- replicate(NBOOT, sample(sub_idx, NSUB, replace=TRUE))
boot_ICC <- matrix(0, nrow=NBOOT, ncol=1)
for (boot_idx in 1:NBOOT){
    boot_dat <- dat[boot_samples[,bootidx],]
    stats <- icc(boot_dat, model='two way', type='agreement', unit='single')
    boot_ICC[boot_idx,] <- stats$value
}

## jackknife resampling (used in CI calculation)
jk_ICC <- matrix(0, nrow=NSUB, ncol=1)
for (sub_idx in 1:NSUB){
    jk_dat <- dat[-sub_idx,]
    stats <- icc(jk_dat, model='two way', type='agreement', unit='single')
    jk_ICC[sub_idx,] <- stats$value
}

## Bca confidence interval
## ICCmat is the ICC of all parcels or edges, where the column represents the parcel or edge and there is only one row.
## boot_ICCmat is the ICC of all parcels or edges for bootstrap samples, where the column represents the parcel or edge and the row represents bootstrap samples.
## jk_ICCmat is the ICC of all parcels or edges for jackknife samples, where the column represents the parcel or edge and the row represents jackknife samples.
## Of note, here we present the CI for mean ICC, while in the manuscript, we calculated the CI for the difference of mean ICC between two methods or pipelines.

meanICC <- rowMeans(ICCmat[1,])
boot_meanICC <- rowMeans(boot_ICCmat)
jk_meanICC <- rowMeans(jk_ICCmat)
```

```
alpha <- 0.05
z0 <- qnorm(mean(boot_meanICC < meanICC))
U <- meanICC - jk_meanICC
a <- sum(U^3)/(6*sum(U^2)^(3/2))
lb <- pnorm(z0+(z0+qnorm(alpha/2))/(1-a*(z0+qnorm(alpha/2))))
ub <- pnorm(z0+(z0+qnorm(1-alpha/2))/(1-a*(z0+qnorm(1-alpha/2))))
CI <- quantile(boot_meanICC, c(lb, ub))
```

6. The codes to perform dbICC calculation and age predication using SVR and CPM models

```
## dbICC calculation
## The dat variable contains the brain feature of a parcel or edge in each column, and the rows of dat are ordered by subjects
## NSUB means the number of subjects, NREP means the number of sessions for each subject
library(dbicc)
distdat <- as.matrix(dist(dat, method= 'euclidean'))
dbICC <- dm2icc(distdat, NSUB, NREP)
CI <- dm2icc.bt(distdat, NSUB, NREP, 2000)

## Age predication using SVR
## ------------------------------------------------
## custom functions used in SVR
## K-fold CV
SVR_CV <- function(x, y, K, C, eps){
  N <- nrow(x)
  y_predict <- numeric(length = N)
  ## randomly split the data into K folds
  set.seed(100)
  rand_idx <- sample(N)
  folds <- cut(c(1:N), breaks=K, labels=FALSE)
  for (curr_fold in c(1:K)){
    fold_idx <- which(folds == curr_fold, arr.ind=TRUE)
    x_train <- x[-rand_idx[fold_idx],]
    y_train <- y[-rand_idx[fold_idx]]
    x_test <- x[rand_idx[fold_idx],,drop=FALSE]
    ## fit SVR model
    model_train <- svm(x_train, y_train, kernel='linear', cost=C, epsilon=eps)
    ## predict
    y_predict[rand_idx[fold_idx]] <- predict(model_train, x_test)
  }
  ## evaluate performance
  acc <- cor(y, y_predict)
  return(acc)
}
## hyper-parameter tuning
SVR_Best <- function(x, y, K){
  C <- 10^(-2:2)
  eps <- seq(0.1,1,0.2)
  hp_all <- expand.grid(C, eps)
  N <- nrow(hp_all)
  acc_all <- NULL
  for (curr_eps in eps){
```

```
        for (curr_C in C){
            curr_acc <- SVR_CV(x, y, K, C=curr_C, eps=curr_eps)
            acc_all <- rbind(acc_all, data.frame(C=curr_C, eps=curr_eps, acc=curr_acc))
        }
    }
    max_idx <- which.max(acc_all$acc)
    return(c(acc_all$C[max_idx], acc_all$eps[max_idx]))
}
## nested CV
SVR_nestCV <- function(x, y, K_out, K_in){
    N <- nrow(x)
    y_predict <- numeric(length = N)
    ## randomly split the data into K_out folds
    set.seed(100)
    rand_idx <- sample(N)
    folds <- cut(c(1:N), breaks=K_out, labels=FALSE)
    for (curr_fold in c(1:K_out)){
        fold_idx <- which(folds == curr_fold, arr.ind=TRUE)
        x_train <- x[-rand_idx[fold_idx],]
        y_train <- y[-rand_idx[fold_idx]]
        x_test <- x[rand_idx[fold_idx],,drop=FALSE]
        ## hyper-parameter tuning
        best_hp <- SVR_Best(x_train, y_train, K_in)
        ## fit SVR model
        model_train <- svm(x_train, y_train, kernel='linear', cost=best_hp[1], epsilon=best_hp[2])
        ## predict
        y_predict[rand_idx[fold_idx]] <- predict(model_train, x_test)
    }
    ## evaluate performance
    acc <- cor(y, y_predict)
    return(acc)
}
## -------------------------------------------------
## invoke custom functions on real data
## the columns of dat contains all parcels or edges of a brain feature, and the rows contains all subjects
## Age means the age of all subjects
library(e1071)
CVR <- SVR_nestCV(as.matrix(dat), Age, 10, 10)

## Age predication using CPM
## -------------------------------------------------
## custom functions used in CPM
## calculate correlation matrix
```

```
calcCORR <- function(x,y){
  N <- nrow(x)
  R <- cor(y, x)
  Tvalue <- R/sqrt((1-R^2)/(N-2))
  Pvalue <- 2*pt(abs(Tvalue), N-2, lower.tail=FALSE)
  CORR <- rbind(R, Pvalue)
  return(CORR)
}
## create feature selection mask
createMask <- function(CORR, pthr){
  R <- CORR[1,]
  Pvalue <- CORR[2,]
  posmask <- (R > 0) * (Pvalue < pthr)
  negmask <- (R < 0) * (Pvalue < pthr)
  mask <- rbind(posmask,negmask)
  return(mask)
}
## summarize features by summing selected features
sumFeature <- function(x, mask){
  posmask <- mask[1,]
  negmask <- mask[2,]
  posSum <- rowSums(x[,posmask == 1, drop=FALSE])
  negSum <- rowSums(x[,negmask == 1, drop=FALSE])
  allSum <- data.frame(Pos=posSum, Neg=negSum)
  return(allSum)
}
## K-fold CV
CPM_CV <- function(x, y, K, pthr){
  N <- nrow(x)
  y_predict <- numeric(length = N)
  ## randomly split the data into K folds
  set.seed(100)
  rand_idx <- sample(N)
  folds <- cut(c(1:N), breaks=K, labels=FALSE)
  for (curr_fold in c(1:K)){
    fold_idx <- which(folds == curr_fold, arr.ind=TRUE)
    x_train <- x[-rand_idx[fold_idx],]
    y_train <- y[-rand_idx[fold_idx]]
    x_test <- x[rand_idx[fold_idx],,drop=FALSE]
    ## fit CPM model
    CORR_train <- calcCORR(x_train, y_train)
    mask_train <- createMask(CORR_train, pthr)
    sum_train <- sumFeature(x_train, mask_train)
    model_train <- lm(y_train ~ Pos + Neg, data = sum_train)
```

```r
        ## predict
        sum_test <- sumFeature(x_test, mask_train)
        y_predict[rand_idx[fold_idx]] <- predict(model_train, sum_test)
    }
    ## evaluate performance
    acc <- cor(y, y_predict)
    return(acc)
}
## hyper-parameter tuning
CPM_Best <- function(x, y, K){
    pthr <- c(0.05, 0.01, 0.005, 0.001)
    acc_all <- NULL
    for (curr_pthr in pthr){
            curr_acc <- CPM_CV(x, y, K, pthr=curr_pthr)
            acc_all <- rbind(acc_all, data.frame(pthr=curr_pthr, acc=curr_acc))
    }
    max_idx <- which.max(acc_all$acc)
    return(acc_all$pthr[max_idx])
}
## nested CV
CPM_nestCV <- function(x, y, K_out, K_in){
    N <- nrow(x)
    y_predict <- numeric(length = N)
    ## randomly split the data into K_out folds
    set.seed(100)
    rand_idx <- sample(N)
    folds <- cut(c(1:N), breaks=K_out, labels=FALSE)
    for (curr_fold in c(1:K_out)){
        fold_idx <- which(folds == curr_fold, arr.ind=TRUE)
        x_train <- x[-rand_idx[fold_idx],]
        y_train <- y[-rand_idx[fold_idx]]
        x_test <- x[rand_idx[fold_idx],,drop=FALSE]
        ## hyper-parameter tuning
        best_hp <- CPM_Best(x_train, y_train, K_in)
        ## fit CPM model
        CORR_train <- calcCORR(x_train, y_train)
        mask_train <- createMask(CORR_train, pthr=best_hp)
        sum_train <- sumFeature(x_train, mask_train)
        model_train <- lm(y_train ~ Pos + Neg, data = sum_train)
        ## predict
        sum_test <- sumFeature(x_test, mask_train)
        y_predict[rand_idx[fold_idx]] <- predict(model_train, sum_test)
    }
    ## evaluate performance
```

```
  acc <- cor(y, y_predict)
  return(acc)
}
## -------------------------------------------------
## invoke custom functions on real data
## the columns of dat contains all parcels or edges of a brain feature, and the rows contains all subjects
## Age means the age of all subjects
CVR <- CPM_nestCV(dat, Age, 10, 10)
```

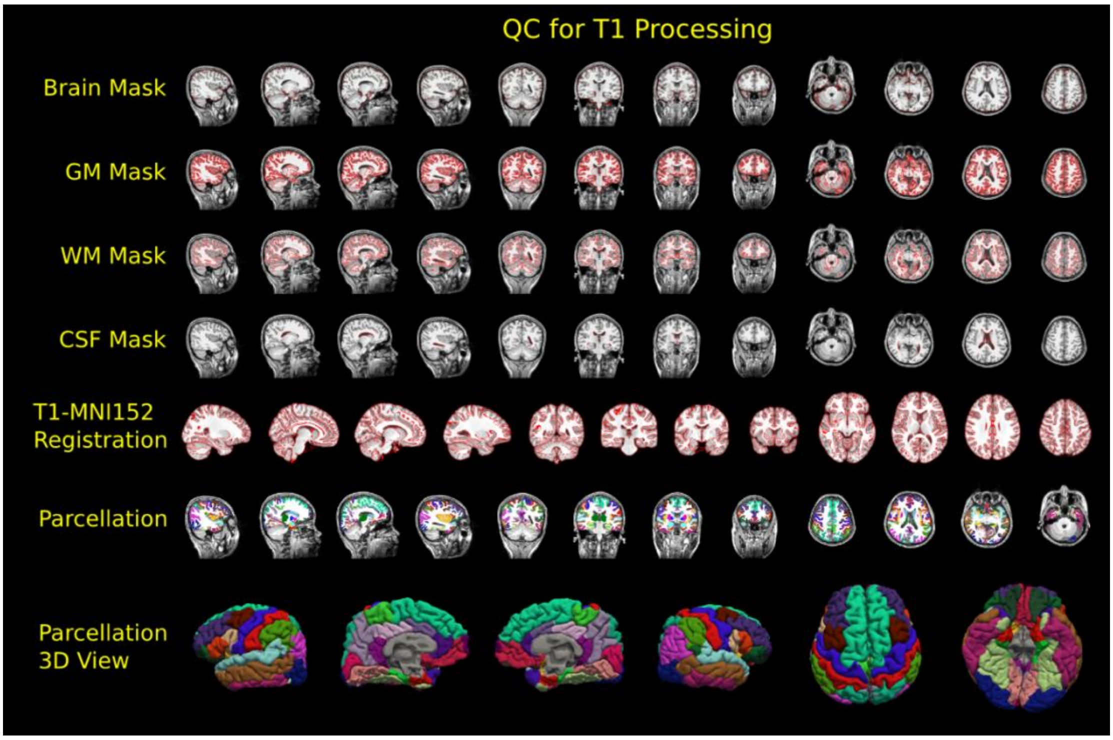


Figure S1. The quality control pictures of T1 processing for one example subject

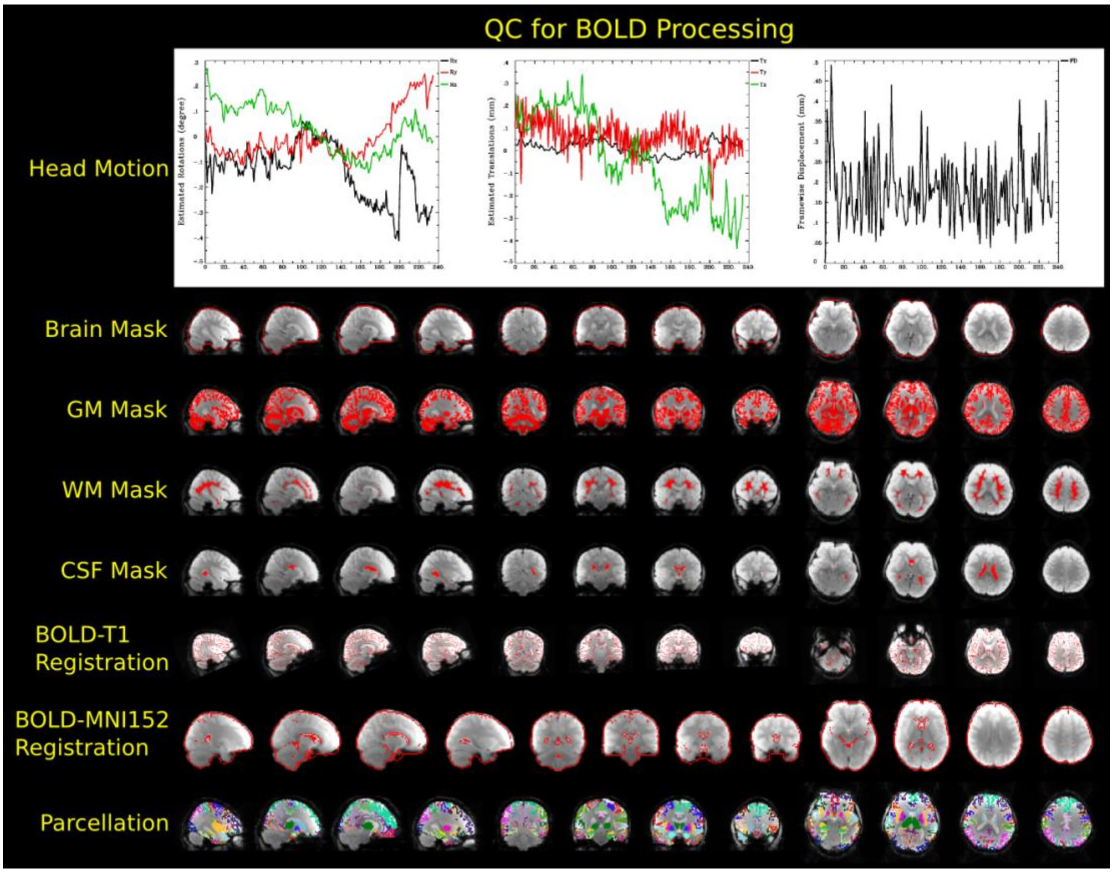


Figure S2. The quality control pictures of BOLD processing for one example subject

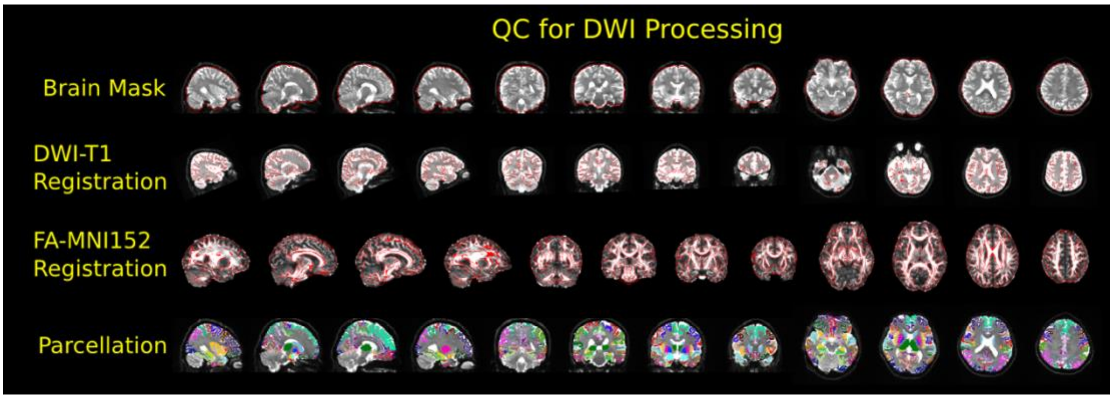


Figure S3. The quality control pictures of DWI processing for one example subject

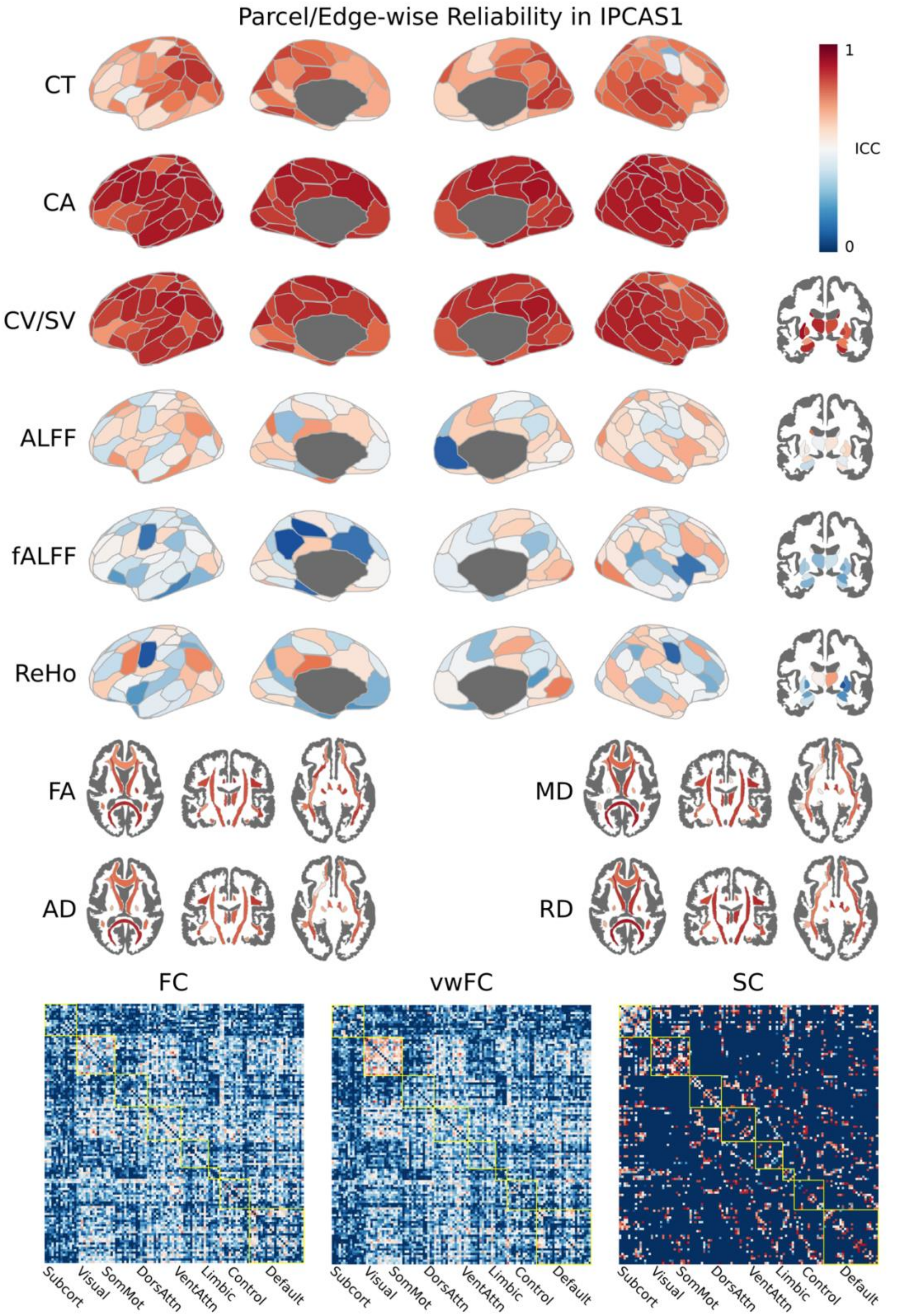


Figure S4. The parcel-wise or edge-wise ICC for all features in the IPCAS1 dataset using Schaefer+Aseg Atlases and JHU Tract Atlas.

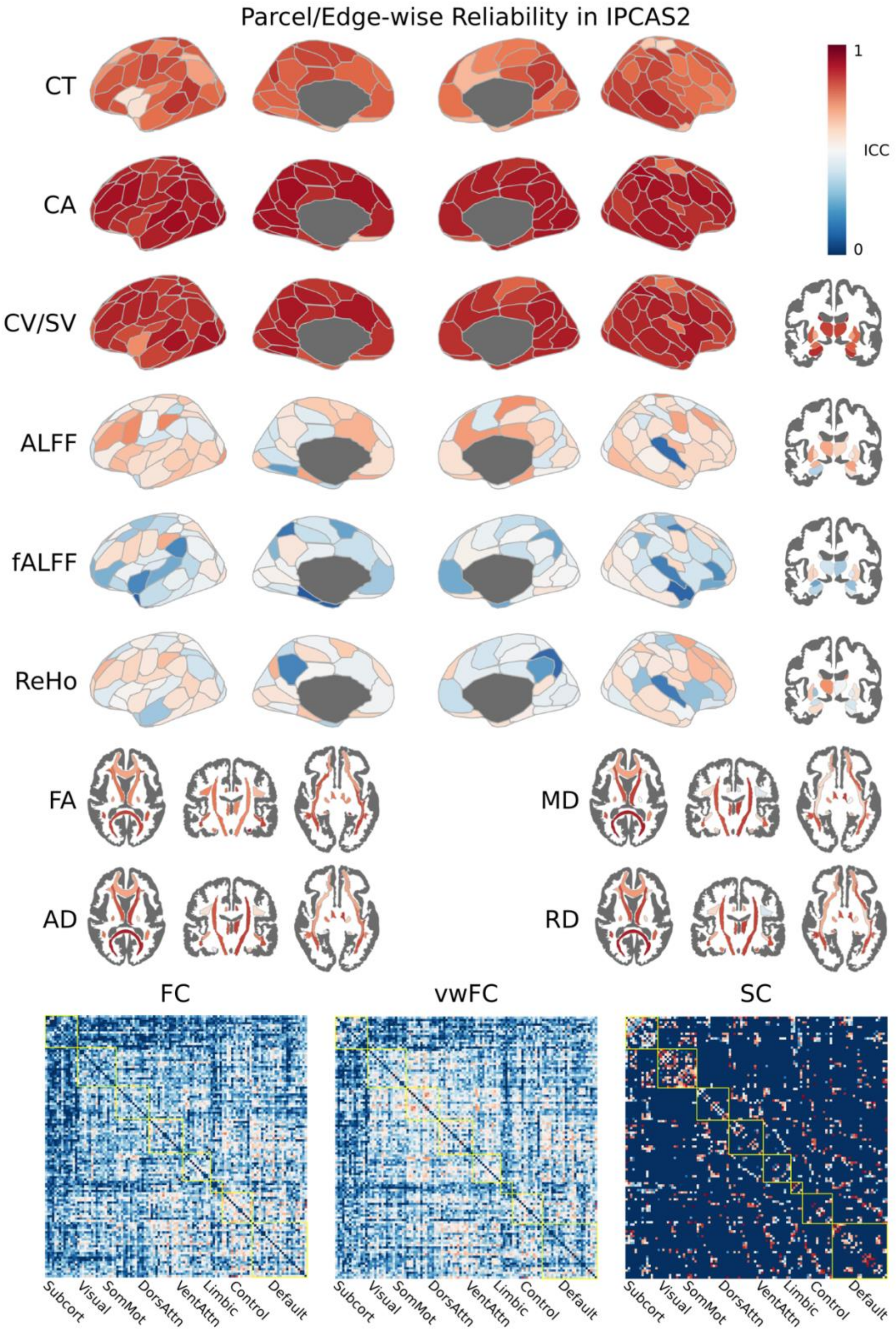


Figure S5. The parcel-wise or edge-wise ICC for all features in the IPCAS2 dataset using Schaefer+Aseg Atlases and JHU Tract Atlas.

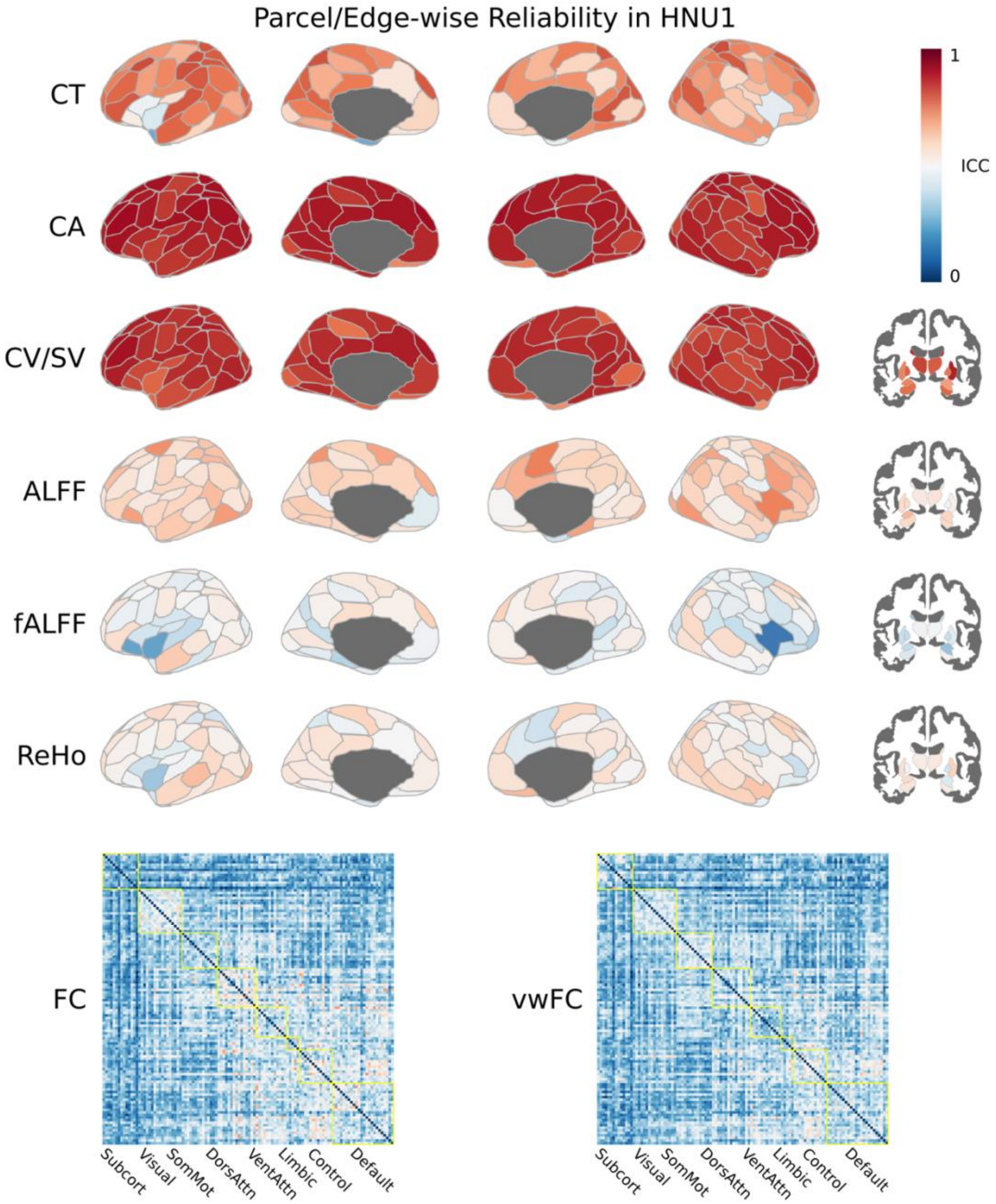


Figure S6. The parcel-wise or edge-wise ICC for all features in the HNU1 dataset using Schaefer+Aseg Atlases and JHU Tract Atlas.

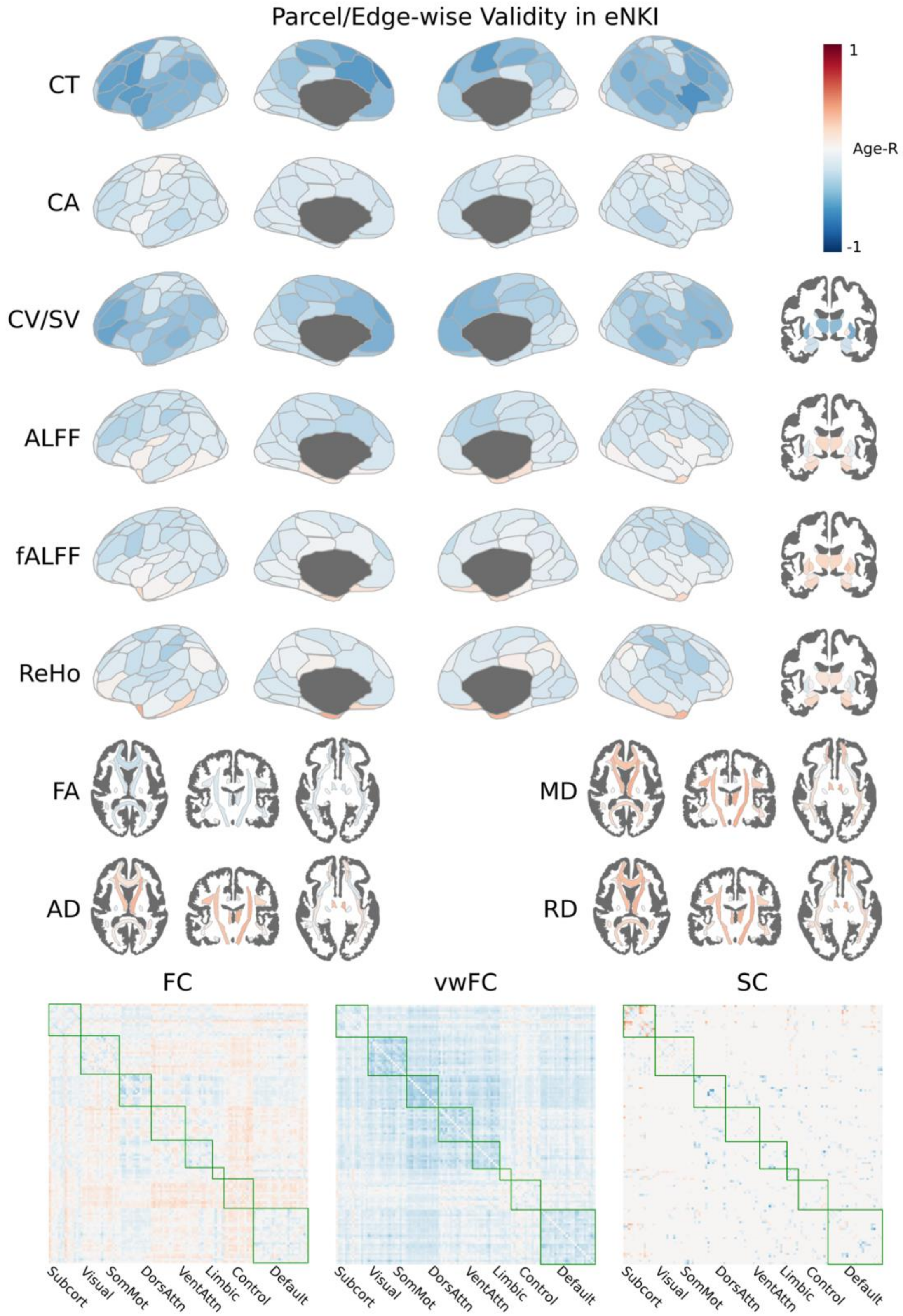


Figure S7. The parcel-wise or edge-wise Age-R for all features in the eNKI dataset using Schaefer+Aseg Atlases and JHU Tract Atlas.

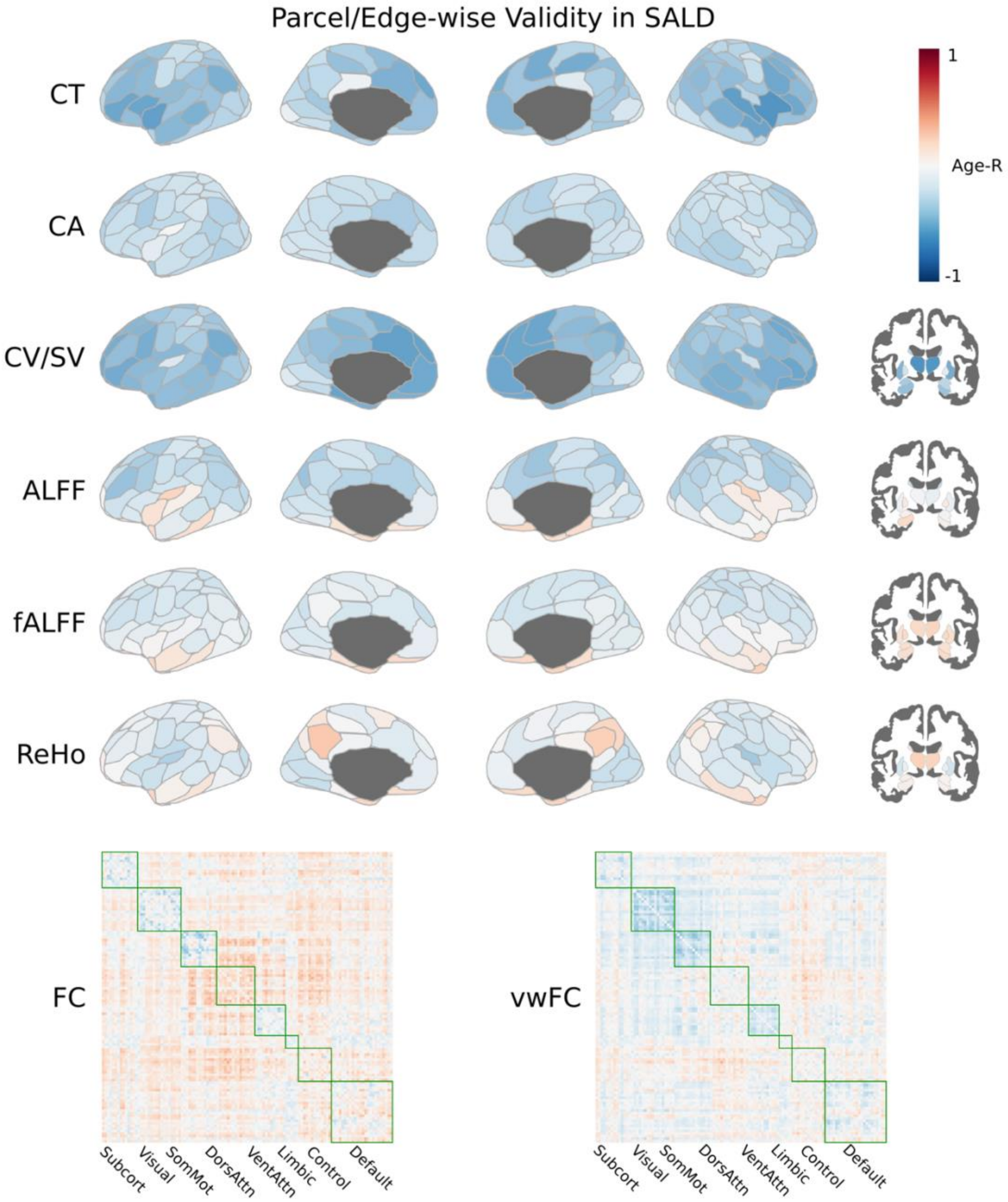


Figure S8. The parcel-wise or edge-wise Age-R for all features in the SALD dataset using Schaefer+Aseg Atlases and JHU Tract Atlas.

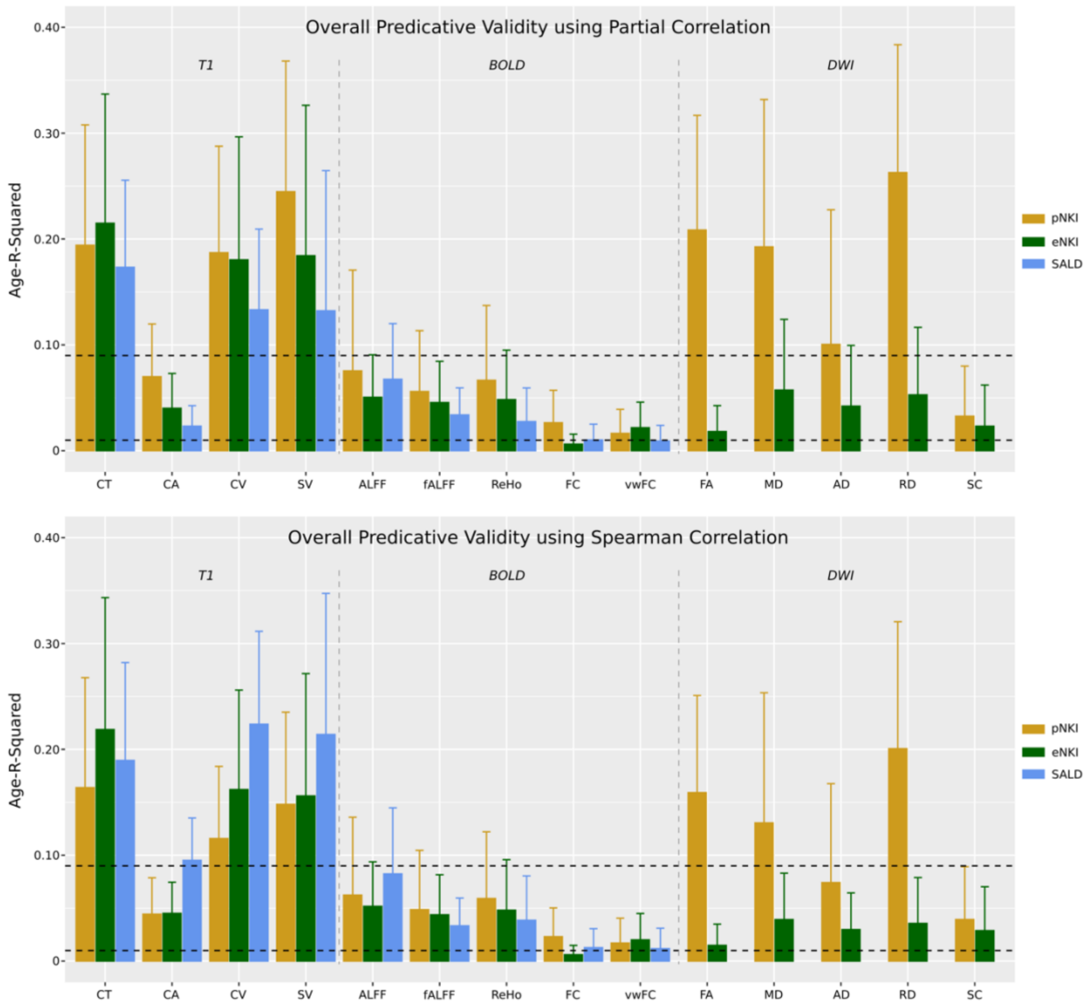


Figure S9. The mean and standard deviation of Age-R-Squared for all brain features across all datasets using Schaefer+Aseg Atlases and JHU Tract Atlas. (A) The Age-R was computed using partial correlation with sex and TIV as covariates; (B) The Age-R was computed using Spearman correlation.

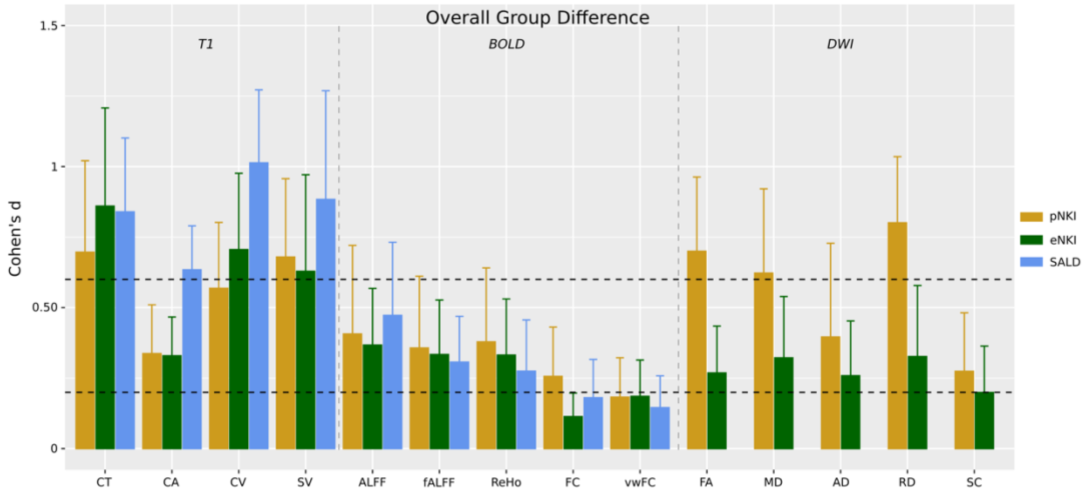


Figure S10. The mean and standard deviation of absolute Cohen's d for all brain features across all datasets using Schaefer+Aseg Atlases and JHU Tract Atlas. The dashed horizontal lines mean d = 0.2 and d = 0.6 separately.

Table S1. A list of pipelines (ordered by publication year) we investigated during the development of PhiPipe

| Pipeline | Publication Year[a] | Citation[b] | URL |
|---|---|---|---|
| DPARSF | 2010 | 2816 | http://www.rfmri.org/DPARSF |
| CONN | 2012 | 3001 | https://www.nitrc.org/projects/conn/ |
| HCP-Pipelines | 2013 | 3171 | https://github.com/Washington-University/HCPpipelines |
| PANDA | 2013 | 509 | https://www.nitrc.org/projects/panda/ |
| C-PAC | 2013 | 149 | https://fcp-indi.github.io/docs/latest/user/index.html |
| GRETNA | 2015 | 758 | https://www.nitrc.org/projects/gretna/ |
| CCS | 2015 | 129 | https://github.com/zuoxinian/CCS |
| volBrain | 2016 | 366 | https://www.volbrain.upv.es/ |
| Biobank Pipeline | 2018 | 605 | https://git.fmrib.ox.ac.uk/falmagro/UK_biobank_pipeline_v_1 |
| Brant | 2018 | 40 | https://sphinx-doc-brant.readthedocs.io/en/latest/ |
| fMRIPrep | 2019 | 1044 | https://fmriprep.org/en/stable/ |
| Ciftify | 2019 | 57 | https://github.com/edickie/ciftify |
| m2g | 2021 | 1 | https://neurodata.io/mri/ |
| micapipe | 2022 | 3 | https://micapipe.readthedocs.io/en/latest/index.html |
| CAT12[c] | 2022 | 295 | http://www.neuro.uni-jena.de/cat12-html/cat.html |
| YeoLab pipeline | *NA* | *NA* | https://github.com/ThomasYeoLab/CBIG/tree/master/stable_projects/preprocessing |

[a] published in a peer-reviewed journal or as a preprint

[b] the citation was based on Google scholar results on June 19, 2022

[c] CAT12 has long been widely used before a standard reference was available

Table S2. A summary of runtime and peak memory usage for processing each modality of a typical subject from each dataset. The processing was run on the CentOS 7 and the CPU is Intel(R) Xeon(R) Silver 4110 CPU @ 2.10GHz.

| Datasets | T1 | REST | DTI[b] | TRACT[c] |
|---|---|---|---|---|
| BNU1 | 9.40 h/4.35 GB[a] | 0.13 h/1.36 GB | 0.35 h/3.95 GB | 8.52 h/2.66 GB |
| IPCAS1 | 8.68 h/4.37 GB | 0.12 h/1.38 GB | 0.47 h/3.50 GB | 12.68 h/2.50 GB |
| IPCAS2 | 9.78 h/4.40 GB | 0.13 h/1.42 GB | 0.32 h/3.18 GB | 9.03 h/2.41 GB |
| HNU1 | 9.88 h/4.35 GB | 0.20 h/2.15 GB | / | / |
| pNKI | 9.47 h/4.40 GB | 0.18 h/1.96 GB | 0.67 h/4.22 GB | 14.08 h/2.63 GB |
| eNKI | 7.55 h/4.38 GB | 0.12 h/0.98 GB | 1.05 h/3.24 GB | 20.55 h/2.27 GB |
| SALD | 8.80 h/4.37 GB | 0.13 h/1.60 GB | / | / |

[a] the runtime (in hour)/peak memory usage (in GB)

[b] the DWI pipeline without tractography (to obtain diffusion tensor measures only)

[c] the tractography processing (to obtain SC measure only) in the DWI pipeline

Table S3. A summary of head motion metrics for resting-state BOLD fMRI data in the finally involved subjects

| Datasets | N[a] | meanFD[b] | Outlier Ratio[c] |
|---|---|---|---|
| BNU1 | 45 (2) | 0.11 (0.033) | 0 (0.051) |
| IPCAS1 | 18 (2) | 0.14 (0.042) | 0 (0.045) |
| IPCAS2 | 27 (2) | 0.11 (0.038) | 0 (0.068) |
| HNU1 | 22 (10) | 0.070 (0.022) | 0 (0.027) |
| pNKI | 113 (1) | 0.16 (0.080) | 0.016 (0.17) |
| eNKI | 502 (1) | 0.17 (0.077) | 0.017 (0.17) |
| SALD | 414 (1) | 0.18 (0.078) | 0.008 (0.19) |

[a] the number of subjects (the number of scans for each subject)

[b] the mean (standard deviation) of meanFD (in millimeter)

[c] the median (maximum) of motion outlier ratio

Table S4. The demographics of younger and older groups used in the group difference detection analysis.

| Dataset | Group | N[a] | Age[b] | Sex[c] |
|---|---|---|---|---|
| pNKI | younger | 63 | 25.86 (5.55) | 34/29 |
| | older | 50 | 54.74 (12.50) | 30/20 |
| eNKI | younger | 244 | 25.92 (5.91) | 122/122 |
| | older | 258 | 58.49 (10.63) | 129/129 |
| SALD | younger | 182 | 25.79 (4.90) | 72/110 |
| | older | 232 | 56.87 (9.44) | 82/150 |

[a] sample size

[b] mean (standard deviation) of age

[c] sample sizes for male/female

Table S5. The summary of reliability for all brain features in the BNU1 dataset

| Modality | Feature | Atlas | N[a] | ICC[b] | Sig[c] | Poor[d] | Moderate[d] | Good[d] |
|---|---|---|---|---|---|---|---|---|
| T1 | CT | Schaefer | 100 | 0.84 (0.065) | 100 | 0 | 12 | 88 |
| | | DK | 68 | 0.81 (0.11) | 100 | 3 | 16 | 81 |
| | CA | Schaefer | 100 | 0.98 (0.010) | 100 | 0 | 0 | 100 |
| | | DK | 68 | 0.95 (0.064) | 100 | 0 | 4 | 96 |
| | CV | Schaefer | 100 | 0.96 (0.019) | 100 | 0 | 0 | 100 |
| | | DK | 68 | 0.94 (0.061) | 100 | 0 | 3 | 97 |
| | SV | Aseg | 14 | 0.93 (0.050) | 100 | 0 | 0 | 100 |
| BOLD | ALFF | Schaefer+Aseg | 114 | 0.53 (0.13) | 99 | 38 | 59 | 3 |
| | | DK+Aseg | 82 | 0.53 (0.14) | 99 | 44 | 52 | 4 |
| | fALFF | Schaefer+Aseg | 114 | 0.42 (0.14) | 88 | 68 | 32 | 0 |
| | | DK+Aseg | 82 | 0.40 (0.14) | 84 | 77 | 23 | 0 |
| | ReHo | Schaefer+Aseg | 114 | 0.46 (0.14) | 90 | 52 | 48 | 0 |
| | | DK+Aseg | 82 | 0.45 (0.16) | 91 | 61 | 38 | 1 |
| | FC | Schaefer+Aseg | 6441 | 0.27 (0.15) | 56 | 94 | 6 | 0 |
| | | DK+Aseg | 3321 | 0.26 (0.15) | 55 | 95 | 5 | 0 |
| | vwFC | Schaefer+Aseg | 6441 | 0.28 (0.16) | 58 | 92 | 8 | 0 |
| | | DK+Aseg | 3321 | 0.27 (0.15) | 59 | 94 | 6 | 0 |
| DWI | FA | JHU-Tract | 20 | 0.92 (0.028) | 100 | 0 | 0 | 100 |
| | | JHU-Label | 48 | 0.88 (0.082) | 100 | 0 | 6 | 94 |
| | MD | JHU-Tract | 20 | 0.81 (0.11) | 100 | 0 | 35 | 65 |
| | | JHU-Label | 48 | 0.77 (0.15) | 100 | 6 | 33 | 61 |
| | AD | JHU-Tract | 20 | 0.82 (0.087) | 100 | 0 | 20 | 80 |
| | | JHU-Label | 48 | 0.80 (0.14) | 100 | 4 | 23 | 73 |
| | RD | JHU-Tract | 20 | 0.87 (0.061) | 100 | 0 | 5 | 95 |
| | | JHU-Label | 48 | 0.83 (0.11) | 100 | 0 | 25 | 75 |
| | SC | Schaefer+Aseg | 854 | 0.65 (0.19) | 97 | 21 | 46 | 33 |
| | | DK+Aseg | 761 | 0.61 (0.22) | 92 | 29 | 37 | 34 |

[a] the number of parcels or edges for a brain feature

[b] the mean (standard deviation) of ICC

[c] the percentage of significant ICCs. The significance level was $p<0.05$ (uncorrected for multiple comparisons).

[d] the percentage of ICCs belonging to different levels of reliability (Poor/Moderate/Good)

Table S6. The summary of reliability for all brain features in the IPCAS1 dataset

| Modality | Feature | Atlas | N[a] | ICC[b] | Sig[c] | Poor[d] | Moderate[d] | Good[d] |
|---|---|---|---|---|---|---|---|---|
| T1 | CT | Schaefer | 100 | 0.79 (0.13) | 99 | 3 | 27 | 70 |
| | | DK | 68 | 0.75 (0.16) | 97 | 10 | 28 | 62 |
| | CA | Schaefer | 100 | 0.96 (0.033) | 100 | 0 | 0 | 100 |
| | | DK | 68 | 0.92 (0.12) | 99 | 3 | 4 | 93 |
| | CV | Schaefer | 100 | 0.93 (0.049) | 100 | 0 | 2 | 98 |
| | | DK | 68 | 0.91 (0.087) | 100 | 0 | 4 | 96 |
| | SV | Aseg | 14 | 0.91 (0.070) | 100 | 0 | 0 | 100 |
| BOLD | ALFF | Schaefer+Aseg | 114 | 0.57 (0.14) | 89 | 27 | 65 | 8 |
| | | DK+Aseg | 82 | 0.55 (0.16) | 88 | 33 | 56 | 11 |
| | fALFF | Schaefer+Aseg | 114 | 0.45 (0.18) | 68 | 59 | 38 | 3 |
| | | DK+Aseg | 82 | 0.43 (0.19) | 66 | 60 | 36 | 4 |
| | ReHo | Schaefer+Aseg | 114 | 0.45 (0.20) | 63 | 59 | 37 | 4 |
| | | DK+Aseg | 82 | 0.46 (0.20) | 65 | 49 | 47 | 4 |
| | FC | Schaefer+Aseg | 6441 | 0.24 (0.20) | 25 | 90 | 10 | 0 |
| | | DK+Aseg | 3321 | 0.22 (0.18) | 20 | 92 | 8 | 0 |
| | vwFC | Schaefer+Aseg | 6441 | 0.23 (0.19) | 23 | 90 | 10 | 0 |
| | | DK+Aseg | 3321 | 0.20 (0.18) | 18 | 93 | 7 | 0 |
| DWI | FA | JHU-Tract | 20 | 0.89 (0.050) | 100 | 0 | 0 | 100 |
| | | JHU-Label | 48 | 0.83 (0.13) | 98 | 2 | 19 | 79 |
| | MD | JHU-Tract | 20 | 0.81 (0.13) | 100 | 0 | 30 | 70 |
| | | JHU-Label | 48 | 0.73 (0.22) | 92 | 17 | 27 | 56 |
| | AD | JHU-Tract | 20 | 0.82 (0.10) | 100 | 0 | 20 | 80 |
| | | JHU-Label | 48 | 0.78 (0.16) | 98 | 6 | 31 | 63 |
| | RD | JHU-Tract | 20 | 0.87 (0.069) | 100 | 0 | 10 | 90 |
| | | JHU-Label | 48 | 0.79 (0.17) | 96 | 10 | 17 | 73 |
| | SC | Schaefer+Aseg | 1149 | 0.60 (0.25) | 79 | 32 | 35 | 33 |
| | | DK+Aseg | 957 | 0.57 (0.26) | 76 | 36 | 34 | 30 |

[a] the number of parcels or edges for a brain feature

[b] the mean (standard deviation) of ICC

[c] the percentage of significant ICCs. The significance level was $p<0.05$ (uncorrected for multiple comparisons).

[d] the percentage of ICCs belonging to different levels of reliability (Poor/Moderate/Good)

Table S7. The summary of reliability for all brain features in the IPCAS2 dataset

| Modality | Feature | Atlas | N[a] | ICC[b] | Sig[c] | Poor[d] | Moderate[d] | Good[d] |
|---|---|---|---|---|---|---|---|---|
| T1 | CT | Schaefer | 100 | 0.85 (0.074) | 100 | 0 | 11 | 89 |
| | | DK | 68 | 0.82 (0.12) | 100 | 4 | 15 | 81 |
| | CA | Schaefer | 100 | 0.96 (0.040) | 100 | 0 | 1 | 99 |
| | | DK | 68 | 0.92 (0.095) | 100 | 0 | 7 | 93 |
| | CV | Schaefer | 100 | 0.94 (0.037) | 100 | 0 | 0 | 100 |
| | | DK | 68 | 0.92 (0.10) | 100 | 1 | 6 | 93 |
| | SV | Aseg | 14 | 0.88 (0.073) | 100 | 0 | 7 | 93 |
| BOLD | ALFF | Schaefer+Aseg | 114 | 0.57 (0.15) | 96 | 26 | 68 | 6 |
| | | DK+Aseg | 82 | 0.58 (0.15) | 96 | 24 | 71 | 5 |
| | fALFF | Schaefer+Aseg | 114 | 0.40 (0.16) | 73 | 70 | 30 | 0 |
| | | DK+Aseg | 82 | 0.39 (0.17) | 70 | 71 | 29 | 0 |
| | ReHo | Schaefer+Aseg | 114 | 0.51 (0.14) | 91 | 45 | 53 | 2 |
| | | DK+Aseg | 82 | 0.50 (0.15) | 87 | 49 | 47 | 4 |
| | FC | Schaefer+Aseg | 6441 | 0.29 (0.18) | 45 | 87 | 13 | 0 |
| | | DK+Aseg | 3321 | 0.26 (0.17) | 41 | 91 | 9 | 0 |
| | vwFC | Schaefer+Aseg | 6441 | 0.32 (0.17) | 52 | 84 | 16 | 0 |
| | | DK+Aseg | 3321 | 0.30 (0.17) | 49 | 88 | 12 | 0 |
| DWI | FA | JHU-Tract | 20 | 0.85 (0.070) | 100 | 0 | 10 | 90 |
| | | JHU-Label | 48 | 0.82 (0.10) | 100 | 2 | 19 | 79 |
| | MD | JHU-Tract | 20 | 0.72 (0.16) | 100 | 10 | 45 | 45 |
| | | JHU-Label | 48 | 0.63 (0.22) | 90 | 29 | 38 | 33 |
| | AD | JHU-Tract | 20 | 0.78 (0.13) | 100 | 5 | 35 | 60 |
| | | JHU-Label | 48 | 0.71 (0.17) | 100 | 17 | 37 | 46 |
| | RD | JHU-Tract | 20 | 0.79 (0.13) | 100 | 5 | 25 | 70 |
| | | JHU-Label | 48 | 0.71 (0.18) | 100 | 15 | 39 | 46 |
| | SC | Schaefer+Aseg | 943 | 0.58 (0.24) | 85 | 33 | 41 | 26 |
| | | DK+Aseg | 802 | 0.55 (0.24) | 81 | 40 | 33 | 27 |

[a] the number of parcels or edges for a brain feature

[b] the mean (standard deviation) of ICC

[c] the percentage of significant ICCs. The significance level was $p<0.05$ (uncorrected for multiple comparisons).

[d] the percentage of ICCs belonging to different levels of reliability (Poor/Moderate/Good)

Table S8. The summary of reliability for all brain features in the HNU1 dataset

| Modality | Feature | Atlas | N[a] | ICC[b] | Sig[c] | Poor[d] | Moderate[d] | Good[d] |
|---|---|---|---|---|---|---|---|---|
| T1 | CT | Schaefer | 100 | 0.76 (0.12) | 100 | 5 | 29 | 66 |
| | | DK | 68 | 0.73 (0.14) | 100 | 8 | 29 | 63 |
| | CA | Schaefer | 100 | 0.96 (0.033) | 100 | 0 | 0 | 100 |
| | | DK | 68 | 0.93 (0.073) | 100 | 0 | 3 | 97 |
| | CV | Schaefer | 100 | 0.94 (0.037) | 100 | 0 | 0 | 100 |
| | | DK | 68 | 0.91 (0.096) | 100 | 0 | 10 | 90 |
| | SV | Aseg | 14 | 0.85 (0.077) | 100 | 0 | 14 | 86 |
| BOLD | ALFF | Schaefer+Aseg | 114 | 0.63 (0.082) | 100 | 6 | 85 | 9 |
| | | DK+Aseg | 82 | 0.63 (0.090) | 100 | 6 | 84 | 10 |
| | fALFF | Schaefer+Aseg | 114 | 0.47 (0.10) | 100 | 60 | 40 | 0 |
| | | DK+Aseg | 82 | 0.45 (0.12) | 100 | 62 | 38 | 0 |
| | ReHo | Schaefer+Aseg | 114 | 0.54 (0.085) | 100 | 22 | 78 | 0 |
| | | DK+Aseg | 82 | 0.56 (0.089) | 100 | 22 | 78 | 0 |
| | FC | Schaefer+Aseg | 6441 | 0.34 (0.11) | 100 | 92 | 8 | 0 |
| | | DK+Aseg | 3321 | 0.32 (0.11) | 100 | 95 | 5 | 0 |
| | vwFC | Schaefer+Aseg | 6441 | 0.33 (0.10) | 100 | 95 | 5 | 0 |
| | | DK+Aseg | 3321 | 0.32 (0.099) | 100 | 97 | 3 | 0 |

[a] the number of parcels or edges for a brain feature

[b] the mean (standard deviation) of ICC

[c] the percentage of significant ICCs. The significance level was $p<0.05$ (uncorrected for multiple comparisons).

[d] the percentage of ICCs belonging to different levels of reliability (Poor/Moderate/Good)

Table S9. The summary of predicative validity for all brain features in the pNKI dataset

| Modality | Feature | Atlas | N[a] | Age-R-Squared[b] | Neg[c] | Sig[d] | Low[e] | Moderate[e] | High[e] |
|---|---|---|---|---|---|---|---|---|---|
| T1 | CT | Schaefer | 100 | 0.17 (0.11) | 97 | 88 | 3 | 27 | 70 |
| | | DK | 68 | 0.15 (0.12) | 94 | 79 | 10 | 22 | 68 |
| | CA | Schaefer | 100 | 0.041 (0.033) | 95 | 59 | 25 | 66 | 9 |
| | | DK | 68 | 0.041 (0.030) | 99 | 47 | 16 | 75 | 9 |
| | CV | Schaefer | 100 | 0.11 (0.068) | 100 | 83 | 1 | 44 | 55 |
| | | DK | 68 | 0.10 (0.062) | 97 | 87 | 6 | 44 | 50 |
| | SV | Aseg | 14 | 0.17 (0.095) | 100 | 100 | 0 | 21 | 79 |
| BOLD | ALFF | Schaefer+Aseg | 114 | 0.072 (0.089) | 82 | 50 | 35 | 39 | 26 |
| | | DK+Aseg | 82 | 0.063 (0.087) | 74 | 45 | 40 | 35 | 25 |
| | fALFF | Schaefer+Aseg | 114 | 0.053 (0.053) | 76 | 55 | 25 | 55 | 20 |
| | | DK+Aseg | 82 | 0.054 (0.060) | 67 | 45 | 30 | 44 | 26 |
| | ReHo | Schaefer+Aseg | 114 | 0.062 (0.063) | 82 | 53 | 27 | 45 | 28 |
| | | DK+Aseg | 82 | 0.064 (0.075) | 68 | 50 | 32 | 40 | 28 |
| | FC | Schaefer+Aseg | 6441 | 0.024 (0.028) | 23 | 27 | 43 | 53 | 4 |
| | | DK+Aseg | 3321 | 0.021 (0.024) | 23 | 22 | 47 | 51 | 2 |
| | vwFC | Schaefer+Aseg | 6441 | 0.015 (0.020) | 53 | 14 | 57 | 42 | 1 |
| | | DK+Aseg | 3321 | 0.011 (0.016) | 47 | 9 | 67 | 33 | 0 |
| DWI | FA | JHU-Tract | 20 | 0.22 (0.11) | 100 | 95 | 0 | 15 | 85 |
| | | JHU-Label | 48 | 0.20 (0.12) | 96 | 92 | 6 | 15 | 79 |
| | MD | JHU-Tract | 20 | 0.20 (0.14) | 0 | 90 | 0 | 20 | 80 |
| | | JHU-Label | 48 | 0.17 (0.13) | 8 | 79 | 10 | 23 | 67 |
| | AD | JHU-Tract | 20 | 0.11 (0.13) | 30 | 55 | 25 | 45 | 30 |
| | | JHU-Label | 48 | 0.10 (0.098) | 27 | 73 | 19 | 37 | 44 |
| | RD | JHU-Tract | 20 | 0.27 (0.12) | 0 | 100 | 0 | 0 | 100 |
| | | JHU-Label | 48 | 0.23 (0.12) | 4 | 92 | 4 | 8 | 88 |
| | SC | Schaefer+Aseg | 830 | 0.032 (0.047) | 73 | 30 | 43 | 48 | 9 |
| | | DK+Aseg | 705 | 0.044 (0.064) | 75 | 39 | 35 | 52 | 13 |

[a] the number of parcels or edges for a brain feature

[b] the mean (standard deviation) of Age-R-Squared

[c] the percentage of negative Age-Rs.

[d] the percentage of significant Age-Rs. The significance level was $p<0.05$ (uncorrected for multiple comparisons).

[e] the percentage of Age-Rs belonging to different levels of validity (Low/Moderate/High)

Table S10. The summary of predicative validity for all brain features in the eNKI dataset

| Modality | Feature | Atlas | N[a] | Age-R-Squared[b] | Neg[c] | Sig[d] | Low[e] | Moderate[e] | High[e] |
|---|---|---|---|---|---|---|---|---|---|
| T1 | CT | Schaefer | 100 | 0.21 (0.12) | 98 | 96 | 4 | 15 | 81 |
| | | DK | 68 | 0.20 (0.14) | 97 | 94 | 7 | 19 | 74 |
| | CA | Schaefer | 100 | 0.044 (0.029) | 97 | 92 | 9 | 85 | 6 |
| | | DK | 68 | 0.045 (0.030) | 99 | 84 | 16 | 78 | 6 |
| | CV | Schaefer | 100 | 0.16 (0.089) | 100 | 98 | 2 | 27 | 71 |
| | | DK | 68 | 0.14 (0.086) | 96 | 94 | 6 | 22 | 72 |
| | SV | Aseg | 14 | 0.16 (0.11) | 93 | 93 | 7 | 36 | 57 |
| BOLD | ALFF | Schaefer+Aseg | 114 | 0.051 (0.041) | 82 | 80 | 22 | 61 | 17 |
| | | DK+Aseg | 82 | 0.043 (0.043) | 71 | 72 | 34 | 50 | 16 |
| | fALFF | Schaefer+Aseg | 114 | 0.047 (0.040) | 80 | 78 | 25 | 57 | 18 |
| | | DK+Aseg | 82 | 0.042 (0.045) | 68 | 65 | 40 | 43 | 17 |
| | ReHo | Schaefer+Aseg | 114 | 0.048 (0.047) | 80 | 79 | 25 | 59 | 16 |
| | | DK+Aseg | 82 | 0.047 (0.044) | 72 | 79 | 25 | 57 | 18 |
| | FC | Schaefer+Aseg | 6441 | 0.0063 (0.0094) | 48 | 27 | 80 | 20 | 0 |
| | | DK+Aseg | 3321 | 0.0060 (0.0096) | 57 | 24 | 82 | 18 | 0 |
| | vwFC | Schaefer+Aseg | 6441 | 0.020 (0.023) | 92 | 63 | 43 | 55 | 2 |
| | | DK+Aseg | 3321 | 0.019 (0.020) | 90 | 65 | 42 | 57 | 1 |
| DWI | FA | JHU-Tract | 20 | 0.018 (0.024) | 80 | 50 | 50 | 45 | 5 |
| | | JHU-Label | 48 | 0.025 (0.048) | 75 | 44 | 65 | 27 | 8 |
| | MD | JHU-Tract | 20 | 0.054 (0.064) | 25 | 70 | 35 | 40 | 25 |
| | | JHU-Label | 48 | 0.061 (0.080) | 40 | 69 | 33 | 36 | 31 |
| | AD | JHU-Tract | 20 | 0.040 (0.053) | 45 | 75 | 40 | 45 | 15 |
| | | JHU-Label | 48 | 0.052 (0.057) | 40 | 81 | 21 | 58 | 21 |
| | RD | JHU-Tract | 20 | 0.050 (0.061) | 25 | 60 | 40 | 40 | 20 |
| | | JHU-Label | 48 | 0.054 (0.077) | 33 | 56 | 44 | 31 | 25 |
| | SC | Schaefer+Aseg | 797 | 0.022 (0.037) | 65 | 52 | 53 | 42 | 5 |
| | | DK+Aseg | 736 | 0.027 (0.044) | 65 | 56 | 50 | 43 | 7 |

[a] the number of parcels or edges for a brain feature

[b] the mean (standard deviation) of Age-R-Squared

[c] the percentage of negative Age-Rs.

[d] the percentage of significant Age-Rs. The significance level was $p<0.05$ (uncorrected for multiple comparisons).

[e] the percentage of Age-Rs belonging to different levels of validity (Low/Moderate/High)

Table S11. The summary of predicative validity for all brain features in the SALD dataset

| Modality | Feature | Atlas | N[a] | Age-R-Squared[b] | Neg[c] | Sig[d] | Low[e] | Moderate[e] | High[e] |
|---|---|---|---|---|---|---|---|---|---|
| T1 | CT | Schaefer | 100 | 0.20 (0.091) | 100 | 98 | 2 | 9 | 89 |
| | | DK | 68 | 0.20 (0.11) | 99 | 97 | 3 | 15 | 82 |
| | CA | Schaefer | 100 | 0.10 (0.040) | 99 | 99 | 1 | 42 | 57 |
| | | DK | 68 | 0.10 (0.048) | 100 | 100 | 0 | 46 | 54 |
| | CV | Schaefer | 100 | 0.23 (0.085) | 100 | 100 | 0 | 7 | 93 |
| | | DK | 68 | 0.22 (0.088) | 100 | 100 | 0 | 10 | 90 |
| | SV | Aseg | 14 | 0.22 (0.13) | 100 | 100 | 0 | 14 | 86 |
| BOLD | ALFF | Schaefer+Aseg | 114 | 0.080 (0.061) | 82 | 83 | 18 | 42 | 40 |
| | | DK+Aseg | 82 | 0.070 (0.068) | 67 | 79 | 22 | 45 | 33 |
| | fALFF | Schaefer+Aseg | 114 | 0.037 (0.028) | 79 | 80 | 21 | 75 | 4 |
| | | DK+Aseg | 82 | 0.038 (0.033) | 57 | 76 | 25 | 65 | 10 |
| | ReHo | Schaefer+Aseg | 114 | 0.036 (0.039) | 69 | 64 | 36 | 52 | 12 |
| | | DK+Aseg | 82 | 0.038 (0.036) | 59 | 65 | 35 | 55 | 10 |
| | FC | Schaefer+Aseg | 6441 | 0.013 (0.019) | 30 | 40 | 62 | 37 | 1 |
| | | DK+Aseg | 3321 | 0.0098 (0.015) | 40 | 31 | 71 | 29 | 0 |
| | vwFC | Schaefer+Aseg | 6441 | 0.010 (0.017) | 60 | 33 | 69 | 30 | 1 |
| | | DK+Aseg | 3321 | 0.0087 (0.014) | 66 | 29 | 73 | 27 | 0 |

[a] the number of parcels or edges for a brain feature

[b] the mean (standard deviation) of Age-R-Squared

[c] the percentage of negative Age-Rs.

[d] the percentage of significant Age-Rs. The significance level was $p<0.05$ (uncorrected for multiple comparisons).

[e] the percentage of Age-Rs belonging to different levels of validity (Low/Moderate/High)

Table S12. The summary of predicative validity for all brain features in the pNKI dataset using partial correlation with sex and TIV as covariates

| Modality | Feature | Atlas | N[a] | Age-R-Squared[b] | Neg[c] | Sig[d] | Low[e] | Moderate[e] | High[e] |
|---|---|---|---|---|---|---|---|---|---|
| T1 | CT | Schaefer | 100 | 0.19 (0.11) | 97 | 95 | 5 | 14 | 81 |
| | | DK | 68 | 0.18 (0.12) | 96 | 85 | 7 | 24 | 69 |
| | CA | Schaefer | 100 | 0.070 (0.050) | 99 | 69 | 13 | 56 | 31 |
| | | DK | 68 | 0.072 (0.048) | 100 | 74 | 4 | 68 | 28 |
| | CV | Schaefer | 100 | 0.19 (0.10) | 100 | 97 | 0 | 23 | 77 |
| | | DK | 68 | 0.17 (0.098) | 97 | 90 | 5 | 13 | 82 |
| | SV | Aseg | 14 | 0.25 (0.12) | 100 | 100 | 0 | 7 | 93 |
| BOLD | ALFF | Schaefer+Aseg | 114 | 0.075 (0.095) | 81 | 52 | 31 | 41 | 28 |
| | | DK+Aseg | 82 | 0.065 (0.091) | 74 | 41 | 37 | 38 | 26 |
| | fALFF | Schaefer+Aseg | 114 | 0.056 (0.058) | 76 | 55 | 22 | 57 | 21 |
| | | DK+Aseg | 82 | 0.057 (0.066) | 66 | 48 | 33 | 41 | 26 |
| | ReHo | Schaefer+Aseg | 114 | 0.067 (0.071) | 84 | 54 | 28 | 40 | 32 |
| | | DK+Aseg | 82 | 0.068 (0.084) | 67 | 49 | 34 | 37 | 29 |
| | FC | Schaefer+Aseg | 6441 | 0.026 (0.031) | 23 | 28 | 41 | 54 | 5 |
| | | DK+Aseg | 3321 | 0.022 (0.025) | 24 | 24 | 45 | 53 | 3 |
| | vwFC | Schaefer+Aseg | 6441 | 0.016 (0.023) | 53 | 14 | 55 | 43 | 2 |
| | | DK+Aseg | 3321 | 0.012 (0.017) | 46 | 10 | 65 | 34 | 1 |
| DWI | FA | JHU-Tract | 20 | 0.21 (0.11) | 100 | 95 | 0 | 15 | 85 |
| | | JHU-Label | 48 | 0.19 (0.11) | 98 | 90 | 6 | 21 | 73 |
| | MD | JHU-Tract | 20 | 0.19 (0.14) | 0 | 90 | 0 | 25 | 75 |
| | | JHU-Label | 48 | 0.16 (0.12) | 13 | 79 | 10 | 27 | 63 |
| | AD | JHU-Tract | 20 | 0.10 (0.13) | 35 | 55 | 25 | 40 | 35 |
| | | JHU-Label | 48 | 0.10 (0.095) | 29 | 69 | 17 | 46 | 38 |
| | RD | JHU-Tract | 20 | 0.26 (0.12) | 0 | 100 | 0 | 0 | 100 |
| | | JHU-Label | 48 | 0.21 (0.12) | 4 | 92 | 4 | 15 | 81 |
| | SC | Schaefer+Aseg | 830 | 0.033 (0.047) | 71 | 31 | 42 | 48 | 10 |
| | | DK+Aseg | 705 | 0.044 (0.066) | 74 | 37 | 35 | 51 | 13 |

[a] the number of parcels or edges for a brain feature

[b] the mean (standard deviation) of Age-R-Squared

[c] the percentage of negative Age-Rs.

[d] the percentage of significant Age-Rs. The significance level was $p<0.05$ (uncorrected for multiple comparisons).

[e] the percentage of Age-Rs belonging to different levels of validity (Low/Moderate/High)

Table S13. The summary of predicative validity for all brain features in the eNKI dataset using partial correlation with sex and TIV as covariates

| Modality | Feature | Atlas | N[a] | Age-R-Squared[b] | Neg[c] | Sig[d] | Low[e] | Moderate[e] | High[e] |
|---|---|---|---|---|---|---|---|---|---|
| T1 | CT | Schaefer | 100 | 0.21 (0.12) | 98 | 96 | 4 | 16 | 80 |
| | | DK | 68 | 0.20 (0.14) | 96 | 93 | 7 | 19 | 74 |
| | CA | Schaefer | 100 | 0.040 (0.033) | 94 | 88 | 14 | 77 | 9 |
| | | DK | 68 | 0.042 (0.036) | 93 | 78 | 23 | 68 | 9 |
| | CV | Schaefer | 100 | 0.18 (0.12) | 99 | 97 | 3 | 24 | 73 |
| | | DK | 68 | 0.16 (0.11) | 96 | 96 | 6 | 23 | 71 |
| | SV | Aseg | 14 | 0.18 (0.14) | 93 | 93 | 7 | 29 | 64 |
| BOLD | ALFF | Schaefer+Aseg | 114 | 0.050 (0.040) | 81 | 79 | 21 | 62 | 17 |
| | | DK+Aseg | 82 | 0.043 (0.043) | 71 | 71 | 33 | 51 | 16 |
| | fALFF | Schaefer+Aseg | 114 | 0.046 (0.039) | 81 | 79 | 24 | 59 | 17 |
| | | DK+Aseg | 82 | 0.041 (0.044) | 71 | 66 | 39 | 44 | 17 |
| | ReHo | Schaefer+Aseg | 114 | 0.048 (0.047) | 80 | 78 | 25 | 60 | 15 |
| | | DK+Aseg | 82 | 0.047 (0.043) | 72 | 78 | 24 | 59 | 17 |
| | FC | Schaefer+Aseg | 6441 | 0.0063 (0.0094) | 50 | 27 | 80 | 20 | 0 |
| | | DK+Aseg | 3321 | 0.0062 (0.0096) | 58 | 25 | 81 | 19 | 0 |
| | vwFC | Schaefer+Aseg | 6441 | 0.022 (0.024) | 93 | 65 | 42 | 56 | 2 |
| | | DK+Aseg | 3321 | 0.020 (0.020) | 91 | 67 | 40 | 59 | 1 |
| DWI | FA | JHU-Tract | 20 | 0.018 (0.024) | 80 | 50 | 50 | 45 | 5 |
| | | JHU-Label | 48 | 0.026 (0.050) | 75 | 42 | 65 | 27 | 8 |
| | MD | JHU-Tract | 20 | 0.057 (0.067) | 25 | 70 | 30 | 45 | 25 |
| | | JHU-Label | 48 | 0.065 (0.083) | 42 | 69 | 33 | 36 | 31 |
| | AD | JHU-Tract | 20 | 0.042 (0.057) | 45 | 75 | 40 | 45 | 15 |
| | | JHU-Label | 48 | 0.056 (0.060) | 42 | 81 | 19 | 60 | 21 |
| | RD | JHU-Tract | 20 | 0.053 (0.064) | 20 | 65 | 40 | 40 | 20 |
| | | JHU-Label | 48 | 0.056 (0.081) | 33 | 58 | 44 | 29 | 27 |
| | SC | Schaefer+Aseg | 797 | 0.023 (0.039) | 65 | 52 | 52 | 42 | 6 |
| | | DK+Aseg | 736 | 0.028 (0.046) | 65 | 55 | 50 | 42 | 8 |

[a] the number of parcels or edges for a brain feature

[b] the mean (standard deviation) of Age-R-Squared

[c] the percentage of negative Age-Rs.

[d] the percentage of significant Age-Rs. The significance level was $p<0.05$ (uncorrected for multiple comparisons).

[e] the percentage of Age-Rs belonging to different levels of validity (Low/Moderate/High)

Table S14. The summary of predicative validity for all brain features in the SALD dataset using partial correlation with sex and TIV as covariates

| Modality | Feature | Atlas | N[a] | Age-R-Squared[b] | Neg[c] | Sig[d] | Low[e] | Moderate[e] | High[e] |
|---|---|---|---|---|---|---|---|---|---|
| T1 | CT | Schaefer | 100 | 0.17 (0.082) | 100 | 98 | 2 | 11 | 87 |
| | | DK | 68 | 0.18 (0.095) | 97 | 97 | 3 | 19 | 78 |
| | CA | Schaefer | 100 | 0.023 (0.019) | 96 | 71 | 30 | 70 | 0 |
| | | DK | 68 | 0.023 (0.023) | 93 | 59 | 41 | 57 | 2 |
| | CV | Schaefer | 100 | 0.13 (0.076) | 99 | 96 | 4 | 30 | 66 |
| | | DK | 68 | 0.13 (0.076) | 100 | 96 | 4 | 28 | 68 |
| | SV | Aseg | 14 | 0.13 (0.13) | 93 | 86 | 14 | 29 | 57 |
| BOLD | ALFF | Schaefer+Aseg | 114 | 0.068 (0.053) | 82 | 81 | 19 | 46 | 35 |
| | | DK+Aseg | 82 | 0.059 (0.060) | 67 | 74 | 27 | 44 | 29 |
| | fALFF | Schaefer+Aseg | 114 | 0.034 (0.026) | 75 | 76 | 25 | 72 | 4 |
| | | DK+Aseg | 82 | 0.034 (0.031) | 55 | 74 | 26 | 66 | 8 |
| | ReHo | Schaefer+Aseg | 114 | 0.028 (0.032) | 74 | 57 | 43 | 53 | 4 |
| | | DK+Aseg | 82 | 0.030 (0.029) | 62 | 65 | 35 | 62 | 2 |
| | FC | Schaefer+Aseg | 6441 | 0.010 (0.015) | 31 | 33 | 69 | 31 | 0 |
| | | DK+Aseg | 3321 | 0.0073 (0.012) | 44 | 25 | 77 | 23 | 0 |
| | vwFC | Schaefer+Aseg | 6441 | 0.0092 (0.015) | 65 | 30 | 72 | 27 | 1 |
| | | DK+Aseg | 3321 | 0.0080 (0.012) | 72 | 27 | 75 | 25 | 0 |

[a] the number of parcels or edges for a brain feature

[b] the mean (standard deviation) of Age-R-Squared

[c] the percentage of negative Age-Rs.

[d] the percentage of significant Age-Rs. The significance level was $p<0.05$ (uncorrected for multiple comparisons).

[e] the percentage of Age-Rs belonging to different levels of validity (Low/Moderate/High)

Table S15. The summary of predicative validity for all brain features in the pNKI dataset using Spearman correlation

| Modality | Feature | Atlas | N[a] | Age-R-Squared[b] | Neg[c] | Sig[d] | Low[e] | Moderate[e] | High[e] |
|---|---|---|---|---|---|---|---|---|---|
| T1 | CT | Schaefer | 100 | 0.16 (0.10) | 97 | 88 | 3 | 24 | 73 |
| | | DK | 68 | 0.15 (0.11) | 93 | 84 | 9 | 22 | 69 |
| | CA | Schaefer | 100 | 0.044 (0.034) | 96 | 61 | 24 | 63 | 13 |
| | | DK | 68 | 0.041 (0.033) | 99 | 49 | 22 | 66 | 12 |
| | CV | Schaefer | 100 | 0.12 (0.068) | 100 | 85 | 2 | 41 | 57 |
| | | DK | 68 | 0.11 (0.063) | 96 | 87 | 9 | 34 | 57 |
| | SV | Aseg | 14 | 0.15 (0.087) | 100 | 100 | 0 | 29 | 71 |
| BOLD | ALFF | Schaefer+Aseg | 114 | 0.062 (0.073) | 87 | 48 | 29 | 44 | 27 |
| | | DK+Aseg | 82 | 0.055 (0.074) | 78 | 41 | 32 | 44 | 24 |
| | fALFF | Schaefer+Aseg | 114 | 0.049 (0.056) | 75 | 45 | 33 | 48 | 19 |
| | | DK+Aseg | 82 | 0.051 (0.062) | 67 | 43 | 38 | 43 | 19 |
| | ReHo | Schaefer+Aseg | 114 | 0.059 (0.063) | 79 | 49 | 31 | 42 | 27 |
| | | DK+Aseg | 82 | 0.058 (0.075) | 67 | 43 | 40 | 33 | 27 |
| | FC | Schaefer+Aseg | 6441 | 0.023 (0.027) | 23 | 24 | 45 | 52 | 3 |
| | | DK+Aseg | 3321 | 0.019 (0.023) | 24 | 20 | 50 | 48 | 2 |
| | vwFC | Schaefer+Aseg | 6441 | 0.017 (0.024) | 55 | 16 | 54 | 44 | 2 |
| | | DK+Aseg | 3321 | 0.012 (0.017) | 50 | 9 | 66 | 33 | 1 |
| DWI | FA | JHU-Tract | 20 | 0.16 (0.092) | 100 | 90 | 5 | 15 | 80 |
| | | JHU-Label | 48 | 0.15 (0.11) | 96 | 75 | 15 | 21 | 64 |
| | MD | JHU-Tract | 20 | 0.13 (0.12) | 5 | 75 | 15 | 35 | 50 |
| | | JHU-Label | 48 | 0.093 (0.10) | 19 | 58 | 25 | 33 | 42 |
| | AD | JHU-Tract | 20 | 0.074 (0.094) | 50 | 50 | 25 | 45 | 30 |
| | | JHU-Label | 48 | 0.061 (0.093) | 40 | 33 | 23 | 58 | 19 |
| | RD | JHU-Tract | 20 | 0.20 (0.12) | 0 | 95 | 5 | 10 | 85 |
| | | JHU-Label | 48 | 0.14 (0.11) | 8 | 77 | 8 | 23 | 69 |
| | SC | Schaefer+Aseg | 830 | 0.039 (0.050) | 78 | 37 | 37 | 50 | 13 |
| | | DK+Aseg | 705 | 0.050 (0.066) | 79 | 42 | 29 | 55 | 16 |

[a] the number of parcels or edges for a brain feature

[b] the mean (standard deviation) of Age-R-Squared

[c] the percentage of negative Age-Rs.

[d] the percentage of significant Age-Rs. The significance level was $p<0.05$ (uncorrected for multiple comparisons).

[e] the percentage of Age-Rs belonging to different levels of validity (Low/Moderate/High)

Table S16. The summary of predicative validity for all brain features in the eNKI dataset using Spearman correlation

| Modality | Feature | Atlas | N[a] | Age-R-Squared[b] | Neg[c] | Sig[d] | Low[e] | Moderate[e] | High[e] |
|---|---|---|---|---|---|---|---|---|---|
| T1 | CT | Schaefer | 100 | 0.22 (0.12) | 99 | 96 | 4 | 15 | 81 |
| | | DK | 68 | 0.20 (0.14) | 97 | 93 | 9 | 16 | 75 |
| | CA | Schaefer | 100 | 0.045 (0.029) | 97 | 93 | 10 | 84 | 6 |
| | | DK | 68 | 0.046 (0.031) | 97 | 88 | 16 | 77 | 7 |
| | CV | Schaefer | 100 | 0.16 (0.094) | 100 | 98 | 2 | 25 | 73 |
| | | DK | 68 | 0.15 (0.090) | 96 | 94 | 6 | 22 | 72 |
| | SV | Aseg | 14 | 0.16 (0.12) | 93 | 86 | 14 | 29 | 57 |
| BOLD | ALFF | Schaefer+Aseg | 114 | 0.052 (0.042) | 82 | 80 | 24 | 58 | 18 |
| | | DK+Aseg | 82 | 0.043 (0.043) | 71 | 71 | 38 | 44 | 18 |
| | fALFF | Schaefer+Aseg | 114 | 0.044 (0.038) | 81 | 77 | 27 | 60 | 13 |
| | | DK+Aseg | 82 | 0.039 (0.042) | 66 | 60 | 45 | 39 | 16 |
| | ReHo | Schaefer+Aseg | 114 | 0.048 (0.048) | 81 | 77 | 24 | 60 | 16 |
| | | DK+Aseg | 82 | 0.046 (0.044) | 68 | 77 | 23 | 59 | 18 |
| | FC | Schaefer+Aseg | 6441 | 0.0061 (0.0088) | 48 | 26 | 80 | 20 | 0 |
| | | DK+Aseg | 3321 | 0.0058 (0.0089) | 58 | 24 | 82 | 18 | 0 |
| | vwFC | Schaefer+Aseg | 6441 | 0.020 (0.025) | 89 | 60 | 47 | 50 | 3 |
| | | DK+Aseg | 3321 | 0.018 (0.021) | 91 | 60 | 47 | 52 | 1 |
| DWI | FA | JHU-Tract | 20 | 0.015 (0.020) | 80 | 50 | 55 | 45 | 0 |
| | | JHU-Label | 48 | 0.021 (0.039) | 71 | 44 | 63 | 33 | 4 |
| | MD | JHU-Tract | 20 | 0.039 (0.044) | 35 | 75 | 25 | 55 | 20 |
| | | JHU-Label | 48 | 0.046 (0.065) | 42 | 71 | 37 | 44 | 19 |
| | AD | JHU-Tract | 20 | 0.030 (0.035) | 45 | 65 | 45 | 45 | 10 |
| | | JHU-Label | 48 | 0.044 (0.049) | 40 | 73 | 31 | 52 | 17 |
| | RD | JHU-Tract | 20 | 0.036 (0.043) | 30 | 60 | 45 | 40 | 15 |
| | | JHU-Label | 48 | 0.040 (0.064) | 40 | 60 | 46 | 37 | 17 |
| | SC | Schaefer+Aseg | 797 | 0.029 (0.042) | 73 | 61 | 44 | 49 | 7 |
| | | DK+Aseg | 736 | 0.033 (0.047) | 71 | 63 | 41 | 49 | 10 |

[a] the number of parcels or edges for a brain feature

[b] the mean (standard deviation) of Age-R-Squared

[c] the percentage of negative Age-Rs.

[d] the percentage of significant Age-Rs. The significance level was $p<0.05$ (uncorrected for multiple comparisons).

[e] the percentage of Age-Rs belonging to different levels of validity (Low/Moderate/High)

Table S17. The summary of predicative validity for all brain features in the SALD dataset using Spearman correlation

| Modality | Feature | Atlas | N[a] | Age-R-Squared[b] | Neg[c] | Sig[d] | Low[e] | Moderate[e] | High[e] |
|---|---|---|---|---|---|---|---|---|---|
| T1 | CT | Schaefer | 100 | 0.19 (0.093) | 100 | 100 | 2 | 9 | 89 |
| | | DK | 68 | 0.19 (0.10) | 97 | 97 | 3 | 19 | 78 |
| | CA | Schaefer | 100 | 0.095 (0.040) | 99 | 99 | 1 | 47 | 52 |
| | | DK | 68 | 0.096 (0.048) | 100 | 100 | 0 | 49 | 51 |
| | CV | Schaefer | 100 | 0.22 (0.088) | 100 | 100 | 0 | 8 | 92 |
| | | DK | 68 | 0.21 (0.088) | 100 | 100 | 0 | 10 | 90 |
| | SV | Aseg | 14 | 0.21 (0.13) | 100 | 100 | 0 | 14 | 86 |
| BOLD | ALFF | Schaefer+Aseg | 114 | 0.082 (0.062) | 81 | 83 | 18 | 40 | 42 |
| | | DK+Aseg | 82 | 0.072 (0.069) | 66 | 79 | 22 | 44 | 34 |
| | fALFF | Schaefer+Aseg | 114 | 0.033 (0.026) | 80 | 80 | 23 | 74 | 3 |
| | | DK+Aseg | 82 | 0.034 (0.031) | 60 | 74 | 27 | 66 | 7 |
| | ReHo | Schaefer+Aseg | 114 | 0.039 (0.042) | 68 | 64 | 37 | 48 | 15 |
| | | DK+Aseg | 82 | 0.041 (0.038) | 59 | 68 | 32 | 57 | 11 |
| | FC | Schaefer+Aseg | 6441 | 0.013 (0.018) | 31 | 38 | 63 | 36 | 1 |
| | | DK+Aseg | 3321 | 0.0094 (0.015) | 41 | 30 | 72 | 28 | 0 |
| | vwFC | Schaefer+Aseg | 6441 | 0.012 (0.019) | 59 | 37 | 65 | 34 | 1 |
| | | DK+Aseg | 3321 | 0.0099 (0.016) | 65 | 32 | 70 | 29 | 1 |

[a] the number of parcels or edges for a brain feature

[b] the mean (standard deviation) of Age-R-Squared

[c] the percentage of negative Age-Rs.

[d] the percentage of significant Age-Rs. The significance level was $p<0.05$ (uncorrected for multiple comparisons).

[e] the percentage of Age-Rs belonging to different levels of validity (Low/Moderate/High)

Table S18. The summary of quadratic age-effect for all brain features in the pNKI dataset

| Modality | Feature | Atlas | N[a] | Increased R-Squared[b] | Sig[c] | Better[d] |
|---|---|---|---|---|---|---|
| T1 | CT | Schaefer | 100 | 0.012 (0.022) | 15 | 25 |
| | | DK | 68 | 0.015 (0.020) | 18 | 31 |
| | CA | Schaefer | 100 | 0.012 (0.016) | 11 | 21 |
| | | DK | 68 | 0.010 (0.014) | 9 | 21 |
| | CV | Schaefer | 100 | 0.015 (0.014) | 14 | 39 |
| | | DK | 68 | 0.015 (0.019) | 16 | 32 |
| | SV | Aseg | 14 | 0.012 (0.017) | 14 | 21 |
| BOLD | ALFF | Schaefer+Aseg | 114 | 0.018 (0.022) | 18 | 42 |
| | | DK+Aseg | 82 | 0.017 (0.021) | 17 | 37 |
| | fALFF | Schaefer+Aseg | 114 | 0.017 (0.021) | 22 | 32 |
| | | DK+Aseg | 82 | 0.019 (0.018) | 24 | 43 |
| | ReHo | Schaefer+Aseg | 114 | 0.025 (0.026) | 29 | 46 |
| | | DK+Aseg | 82 | 0.025 (0.028) | 29 | 48 |
| | FC | Schaefer+Aseg | 6441 | 0.016 (0.019) | 16 | 33 |
| | | DK+Aseg | 3321 | 0.015 (0.018) | 14 | 32 |
| | vwFC | Schaefer+Aseg | 6441 | 0.012 (0.014) | 9 | 27 |
| | | DK+Aseg | 3321 | 0.012 (0.014) | 8 | 28 |
| DWI | FA | JHU-Tract | 20 | 0.040 (0.035) | 60 | 70 |
| | | JHU-Label | 48 | 0.044 (0.053) | 44 | 63 |
| | MD | JHU-Tract | 20 | 0.084 (0.054) | 95 | 95 |
| | | JHU-Label | 48 | 0.091 (0.062) | 83 | 90 |
| | AD | JHU-Tract | 20 | 0.067 (0.042) | 75 | 85 |
| | | JHU-Label | 48 | 0.080 (0.064) | 73 | 79 |
| | RD | JHU-Tract | 20 | 0.069 (0.052) | 80 | 95 |
| | | JHU-Label | 48 | 0.075 (0.058) | 75 | 92 |
| | SC | Schaefer+Aseg | 830 | 0.016 (0.025) | 14 | 28 |
| | | DK+Aseg | 705 | 0.016 (0.028) | 13 | 25 |

[a] the number of parcels or edges for a brain feature

[b] the increased R-Squared using quadratic regression. For instance, a value of 0.05 means the model could explain 5% more variances when including the quadratic term of age.

[c] the percentage of significant quadratic term of age. The significance level was $p<0.05$ (uncorrected for multiple comparisons).

[d] the percentage of quadratic models which was the better than simple linear regression models based on the AIC. A better model means the model had a lower AIC.

Table S19. The summary of quadratic age-effect for all brain features in the eNKI dataset

| Modality | Feature | Atlas | N[a] | Increased R-Squared[b] | Sig[c] | Better[d] |
|---|---|---|---|---|---|---|
| T1 | CT | Schaefer | 100 | 0.0051 (0.0074) | 25 | 41 |
| | | DK | 68 | 0.0063 (0.0079) | 31 | 56 |
| | CA | Schaefer | 100 | 0.0016 (0.0022) | 2 | 15 |
| | | DK | 68 | 0.0020 (0.0029) | 4 | 16 |
| | CV | Schaefer | 100 | 0.0073 (0.0076) | 41 | 61 |
| | | DK | 68 | 0.0086 (0.0094) | 47 | 62 |
| | SV | Aseg | 14 | 0.0076 (0.0065) | 50 | 64 |
| BOLD | ALFF | Schaefer+Aseg | 114 | 0.0043 (0.0057) | 18 | 37 |
| | | DK+Aseg | 82 | 0.0050 (0.0080) | 21 | 34 |
| | fALFF | Schaefer+Aseg | 114 | 0.0030 (0.0037) | 11 | 29 |
| | | DK+Aseg | 82 | 0.0025 (0.0023) | 6 | 28 |
| | ReHo | Schaefer+Aseg | 114 | 0.0040 (0.0042) | 21 | 39 |
| | | DK+Aseg | 82 | 0.0046 (0.0046) | 22 | 49 |
| | FC | Schaefer+Aseg | 6441 | 0.0031 (0.0042) | 12 | 27 |
| | | DK+Aseg | 3321 | 0.0026 (0.0039) | 9 | 22 |
| | vwFC | Schaefer+Aseg | 6441 | 0.0021 (0.0026) | 5 | 19 |
| | | DK+Aseg | 3321 | 0.0019 (0.0026) | 5 | 15 |
| DWI | FA | JHU-Tract | 20 | 0.0025 (0.0027) | 10 | 25 |
| | | JHU-Label | 48 | 0.0042 (0.0073) | 23 | 27 |
| | MD | JHU-Tract | 20 | 0.054 (0.035) | 100 | 100 |
| | | JHU-Label | 48 | 0.046 (0.036) | 79 | 85 |
| | AD | JHU-Tract | 20 | 0.076 (0.054) | 95 | 95 |
| | | JHU-Label | 48 | 0.056 (0.044) | 81 | 88 |
| | RD | JHU-Tract | 20 | 0.031 (0.025) | 75 | 90 |
| | | JHU-Label | 48 | 0.028 (0.027) | 71 | 77 |
| | SC | Schaefer+Aseg | 797 | 0.0070 (0.011) | 27 | 44 |
| | | DK+Aseg | 736 | 0.0068 (0.012) | 28 | 41 |

[a] the number of parcels or edges for a brain feature

[b] the increased R-Squared using quadratic regression. For instance, a value of 0.05 means the model could explain 5% more variances on average when including the quadratic term of age.

[c] the percentage of significant quadratic term of age. The significance level was $p<0.05$ (uncorrected for multiple comparisons).

[d] the percentage of quadratic models which was the better than simple linear regression models based on the AIC. A better model means the model had a lower AIC.

Table S20. The summary of quadratic age-effect for all brain features in the SALD dataset

| Modality | Feature | Atlas | N[a] | Increased R-Squared[b] | Sig[c] | Better[d] |
|---|---|---|---|---|---|---|
| T1 | CT | Schaefer | 100 | 0.0027 (0.0033) | 10 | 26 |
| | | DK | 68 | 0.0039 (0.0051) | 15 | 31 |
| | CA | Schaefer | 100 | 0.0023 (0.0031) | 4 | 19 |
| | | DK | 68 | 0.0019 (0.0020) | 1 | 10 |
| | CV | Schaefer | 100 | 0.0046 (0.0053) | 26 | 41 |
| | | DK | 68 | 0.0042 (0.0056) | 18 | 41 |
| | SV | Aseg | 14 | 0.0083 (0.0089) | 43 | 43 |
| BOLD | ALFF | Schaefer+Aseg | 114 | 0.0056 (0.0070) | 23 | 38 |
| | | DK+Aseg | 82 | 0.0064 (0.0076) | 23 | 48 |
| | fALFF | Schaefer+Aseg | 114 | 0.0035 (0.0040) | 11 | 26 |
| | | DK+Aseg | 82 | 0.0037 (0.0053) | 11 | 29 |
| | ReHo | Schaefer+Aseg | 114 | 0.0071 (0.0092) | 25 | 44 |
| | | DK+Aseg | 82 | 0.0071 (0.0081) | 29 | 48 |
| | FC | Schaefer+Aseg | 6441 | 0.0039 (0.0053) | 12 | 27 |
| | | DK+Aseg | 3321 | 0.0038 (0.0050) | 11 | 27 |
| | vwFC | Schaefer+Aseg | 6441 | 0.0028 (0.0035) | 6 | 21 |
| | | DK+Aseg | 3321 | 0.0029 (0.0036) | 6 | 21 |

[a] the number of parcels or edges for a brain feature

[b] the increased R-Squared using quadratic regression. For instance, a value of 0.05 means the model could explain 5% more variances when including the quadratic term of age.

[c] the percentage of significant quadratic term of age. The significance level was $p<0.05$ (uncorrected for multiple comparisons).

[d] the percentage of quadratic models which was the better than simple linear regression models based on the AIC. A better model means the model had a lower AIC.

Table S21. The reproducibility of ICC and Age-R for all brain features across all datasets.

| Modality | Feature | Atlas | ICC_Corr[a] | ICC_MAD[a] | Age-R_Corr[b] | Age-R_MAD[b] |
|---|---|---|---|---|---|---|
| T1 | CT | Schaefer | 0.36 (0.15) | 0.081 (0.025) | 0.82 | 0.082 |
| | | DK | 0.52 (0.039) | 0.095 (0.024) | 0.87 | 0.089 |
| | CA | Schaefer | 0.41 (0.023) | 0.024 (0.0030) | 0.70 | 0.055 |
| | | DK | 0.69 (0.053) | 0.039 (0.0055) | 0.75 | 0.047 |
| | CV | Schaefer | 0.33 (0.084) | 0.032 (0.0065) | 0.87 | 0.076 |
| | | DK | 0.64 (0.045) | 0.039 (0.0039) | 0.89 | 0.075 |
| | SV | Aseg | 0.61 (0.19) | 0.054 (0.0068) | 0.85 | 0.090 |
| BOLD | ALFF | Schaefer+Aseg | 0.058 (0.14) | 0.15 (0.017) | 0.74 | 0.12 |
| | | DK+Aseg | 0.13 (0.21) | 0.15 (0.024) | 0.74 | 0.12 |
| | fALFF | Schaefer+Aseg | 0.084 (0.12) | 0.17 (0.0033) | 0.80 | 0.079 |
| | | DK+Aseg | 0.15 (0.057) | 0.18 (0.011) | 0.85 | 0.089 |
| | ReHo | Schaefer+Aseg | 0.052 (0.12) | 0.18 (0.017) | 0.74 | 0.094 |
| | | DK+Aseg | 0.015 (0.090) | 0.19 (0.0091) | 0.85 | 0.099 |
| | FC | Schaefer+Aseg | 0.15 (0.028) | 0.18 (0.011) | 0.54 | 0.11 |
| | | DK+Aseg | 0.080 (0.032) | 0.19 (0.0085) | 0.53 | 0.11 |
| | vwFC | Schaefer+Aseg | 0.083 (0.078) | 0.20 (0.0076) | 0.66 | 0.11 |
| | | DK+Aseg | 0.026 (0.076) | 0.20 (0.013) | 0.58 | 0.13 |
| DWI | FA | JHU-Tract | 0.0065 (0.086) | 0.067 (0.021) | 0.77 | 0.35 |
| | | JHU-Label | 0.47 (0.14) | 0.086 (0.013) | 0.81 | 0.34 |
| | MD | JHU-Tract | 0.39 (0.20) | 0.12 (0.036) | 0.80 | 0.28 |
| | | JHU-Label | 0.63 (0.12) | 0.16 (0.035) | 0.70 | 0.24 |
| | AD | JHU-Tract | 0.39 (0.075) | 0.089 (0.011) | 0.82 | 0.14 |
| | | JHU-Label | 0.54 (0.071) | 0.11 (0.021) | 0.71 | 0.16 |
| | RD | JHU-Tract | 0.19 (0.16) | 0.091 (0.028) | 0.75 | 0.35 |
| | | JHU-Label | 0.57 (0.14) | 0.13 (0.041) | 0.71 | 0.31 |
| | SC | Schaefer+Aseg | 0.38 (0.056) | 0.18 (0.018) | 0.68 | 0.091 |
| | | DK+Aseg | 0.51 (0.054) | 0.18 (0.015) | 0.75 | 0.092 |

[a] the mean (standard deviation) of correlations and MADs across three datasets for ICC results

[b] the correlation and MAD between two datasets for Age-R results

Table S22. The summary of group difference for all brain features in the pNKI dataset

| Modality | Feature | Atlas | N[a] | Cohen's d[b] | Neg[c] | Sig[d] | Small[e] | Moderate[e] | Large[e] |
|---|---|---|---|---|---|---|---|---|---|
| T1 | CT | Schaefer | 100 | 0.70 (0.32) | 3 | 79 | 5 | 30 | 65 |
| | | DK | 68 | 0.67 (0.32) | 6 | 78 | 10 | 25 | 65 |
| | CA | Schaefer | 100 | 0.34 (0.17) | 4 | 45 | 27 | 67 | 6 |
| | | DK | 68 | 0.34 (0.16) | 0 | 43 | 25 | 69 | 6 |
| | CV | Schaefer | 100 | 0.57 (0.23) | 0 | 78 | 6 | 44 | 50 |
| | | DK | 68 | 0.55 (0.21) | 4 | 78 | 6 | 51 | 43 |
| | SV | Aseg | 14 | 0.68 (0.28) | 0 | 86 | 7 | 29 | 64 |
| BOLD | ALFF | Schaefer+Aseg | 114 | 0.41 (0.31) | 18 | 43 | 38 | 35 | 27 |
| | | DK+Aseg | 82 | 0.37 (0.32) | 24 | 38 | 41 | 37 | 22 |
| | fALFF | Schaefer+Aseg | 114 | 0.36 (0.25) | 28 | 39 | 33 | 48 | 19 |
| | | DK+Aseg | 82 | 0.36 (0.28) | 38 | 44 | 35 | 45 | 20 |
| | ReHo | Schaefer+Aseg | 114 | 0.38 (0.26) | 18 | 40 | 31 | 46 | 23 |
| | | DK+Aseg | 82 | 0.37 (0.31) | 33 | 40 | 40 | 33 | 27 |
| | FC | Schaefer+Aseg | 6441 | 0.26 (0.17) | 80 | 24 | 43 | 53 | 4 |
| | | DK+Aseg | 3321 | 0.23 (0.16) | 78 | 19 | 49 | 50 | 1 |
| | vwFC | Schaefer+Aseg | 6441 | 0.18 (0.14) | 52 | 10 | 62 | 37 | 1 |
| | | DK+Aseg | 3321 | 0.15 (0.12) | 59 | 6 | 70 | 30 | 0 |
| DWI | FA | JHU-Tract | 20 | 0.70 (0.26) | 0 | 85 | 5 | 25 | 70 |
| | | JHU-Label | 48 | 0.63 (0.32) | 6 | 73 | 15 | 29 | 56 |
| | MD | JHU-Tract | 20 | 0.62 (0.30) | 100 | 75 | 5 | 45 | 50 |
| | | JHU-Label | 48 | 0.53 (0.30) | 90 | 63 | 19 | 31 | 50 |
| | AD | JHU-Tract | 20 | 0.40 (0.33) | 55 | 35 | 35 | 40 | 25 |
| | | JHU-Label | 48 | 0.42 (0.28) | 65 | 46 | 17 | 60 | 23 |
| | RD | JHU-Tract | 20 | 0.80 (0.23) | 100 | 100 | 0 | 25 | 75 |
| | | JHU-Label | 48 | 0.66 (0.29) | 94 | 77 | 8 | 27 | 65 |
| | SC | Schaefer+Aseg | 830 | 0.27 (0.21) | 29 | 27 | 43 | 50 | 7 |
| | | DK+Aseg | 705 | 0.32 (0.25) | 26 | 34 | 38 | 51 | 11 |

[a] the number of parcels or edges for a brain feature

[b] the mean (standard deviation) of absolute Cohen's d

[c] the percentage of the negative group difference (i.e., younger group < older group).

[d] the percentage of significant group difference. The significance level was $p<0.05$ (uncorrected for multiple comparisons).

[e] the percentage of Cohen's d belonging to different levels of effect size (Small/Moderate/Large)

Table S23. The summary of group difference for all brain features in the eNKI dataset

| Modality | Feature | Atlas | N[a] | Cohen's d[b] | Neg[c] | Sig[d] | Small[e] | Moderate[e] | Large[e] |
|---|---|---|---|---|---|---|---|---|---|
| T1 | CT | Schaefer | 100 | 0.86 (0.35) | 1 | 96 | 4 | 20 | 76 |
| | | DK | 68 | 0.81 (0.40) | 3 | 94 | 7 | 27 | 66 |
| | CA | Schaefer | 100 | 0.33 (0.14) | 3 | 86 | 17 | 79 | 4 |
| | | DK | 68 | 0.33 (0.15) | 6 | 79 | 21 | 76 | 3 |
| | CV | Schaefer | 100 | 0.71 (0.27) | 0 | 97 | 3 | 35 | 62 |
| | | DK | 68 | 0.67 (0.27) | 4 | 94 | 6 | 41 | 53 |
| | SV | Aseg | 14 | 0.63 (0.34) | 7 | 93 | 7 | 36 | 57 |
| BOLD | ALFF | Schaefer+Aseg | 114 | 0.37 (0.20) | 19 | 77 | 27 | 63 | 10 |
| | | DK+Aseg | 82 | 0.32 (0.21) | 29 | 67 | 40 | 51 | 9 |
| | fALFF | Schaefer+Aseg | 114 | 0.33 (0.19) | 20 | 72 | 30 | 62 | 8 |
| | | DK+Aseg | 82 | 0.29 (0.22) | 34 | 55 | 49 | 39 | 12 |
| | ReHo | Schaefer+Aseg | 114 | 0.33 (0.20) | 18 | 74 | 30 | 61 | 10 |
| | | DK+Aseg | 82 | 0.33 (0.20) | 31 | 77 | 26 | 62 | 12 |
| | FC | Schaefer+Aseg | 6441 | 0.11 (0.084) | 55 | 21 | 85 | 15 | 0 |
| | | DK+Aseg | 3321 | 0.11 (0.086) | 48 | 20 | 85 | 15 | 0 |
| | vwFC | Schaefer+Aseg | 6441 | 0.19 (0.13) | 13 | 47 | 59 | 40 | 1 |
| | | DK+Aseg | 3321 | 0.18 (0.11) | 15 | 46 | 62 | 38 | 0 |
| DWI | FA | JHU-Tract | 20 | 0.27 (0.17) | 0 | 65 | 45 | 50 | 5 |
| | | JHU-Label | 48 | 0.27 (0.20) | 13 | 63 | 40 | 52 | 8 |
| | MD | JHU-Tract | 20 | 0.32 (0.22) | 75 | 70 | 40 | 40 | 20 |
| | | JHU-Label | 48 | 0.33 (0.26) | 73 | 65 | 42 | 42 | 16 |
| | AD | JHU-Tract | 20 | 0.26 (0.19) | 55 | 55 | 50 | 45 | 5 |
| | | JHU-Label | 48 | 0.31 (0.21) | 58 | 69 | 36 | 58 | 6 |
| | RD | JHU-Tract | 20 | 0.33 (0.25) | 85 | 60 | 40 | 40 | 20 |
| | | JHU-Label | 48 | 0.31 (0.28) | 85 | 54 | 50 | 33 | 17 |
| | SC | Schaefer+Aseg | 797 | 0.20 (0.16) | 36 | 46 | 60 | 38 | 2 |
| | | DK+Aseg | 736 | 0.22 (0.18) | 39 | 50 | 56 | 40 | 4 |

[a] the number of parcels or edges for a brain feature

[b] the mean (standard deviation) of absolute Cohen's d

[c] the percentage of the negative group difference (i.e., younger group < older group).

[d] the percentage of significant group difference. The significance level was $p<0.05$ (uncorrected for multiple comparisons).

[e] the percentage of Cohen's d belonging to different levels of effect size (Small/Moderate/Large)

Table S24. The summary of group difference for all brain features in the SALD dataset

| Modality | Feature | Atlas | N[a] | Cohen's d[b] | Neg[c] | Sig[d] | Small[e] | Moderate[e] | Large[e] |
|---|---|---|---|---|---|---|---|---|---|
| T1 | CT | Schaefer | 100 | 0.84 (0.26) | 0 | 98 | 2 | 15 | 83 |
| | | DK | 68 | 0.84 (0.29) | 3 | 96 | 4 | 18 | 78 |
| | CA | Schaefer | 100 | 0.63 (0.16) | 0 | 99 | 1 | 40 | 59 |
| | | DK | 68 | 0.63 (0.18) | 0 | 100 | 0 | 44 | 56 |
| | CV | Schaefer | 100 | 1.01 (0.26) | 0 | 100 | 0 | 6 | 94 |
| | | DK | 68 | 0.98 (0.28) | 0 | 100 | 0 | 10 | 90 |
| | SV | Aseg | 14 | 0.88 (0.39) | 0 | 93 | 7 | 7 | 86 |
| BOLD | ALFF | Schaefer+Aseg | 114 | 0.47 (0.26) | 21 | 80 | 21 | 42 | 37 |
| | | DK+Aseg | 82 | 0.43 (0.28) | 39 | 77 | 23 | 52 | 24 |
| | fALFF | Schaefer+Aseg | 114 | 0.31 (0.16) | 25 | 69 | 31 | 66 | 3 |
| | | DK+Aseg | 82 | 0.31 (0.17) | 43 | 77 | 26 | 68 | 6 |
| | ReHo | Schaefer+Aseg | 114 | 0.27 (0.18) | 28 | 63 | 40 | 54 | 6 |
| | | DK+Aseg | 82 | 0.30 (0.17) | 40 | 65 | 37 | 58 | 5 |
| | FC | Schaefer+Aseg | 6441 | 0.18 (0.14) | 72 | 39 | 62 | 37 | 1 |
| | | DK+Aseg | 3321 | 0.15 (0.12) | 63 | 31 | 71 | 29 | 0 |
| | vwFC | Schaefer+Aseg | 6441 | 0.15 (0.11) | 44 | 27 | 74 | 26 | 0 |
| | | DK+Aseg | 3321 | 0.14 (0.10) | 38 | 24 | 77 | 23 | 0 |

[a] the number of parcels or edges for a brain feature

[b] the mean (standard deviation) of absolute Cohen's d

[c] the percentage of the negative group difference (i.e., younger group < older group).

[d] the percentage of significant group difference. The significance level was $p<0.05$ (uncorrected for multiple comparisons).

[e] the percentage of Cohen's d belonging to different levels of effect size (Small/Moderate/Large)

Table S25. The reliability changes caused by CAT12's skull stripping for all brain features across all datasets.

| Modality | Feature | Atlas | BNU1 | IPCAS1 | IPCAS2 | HNU1 |
|---|---|---|---|---|---|---|
| T1 | CT | Schaefer | -0.0095 (-0.023, 0.0031)[a] | -0.0047 (-0.038, 0.025) | -0.0074 (-0.044, 0.020) | **-0.016 (-0.027, -0.0019)** |
| | | DK | -0.013 (-0.026, 0.00025) | -0.0074 (-0.051, 0.033) | 0.0021 (-0.032, 0.029) | -0.012 (-0.024, 0.0033) |
| | CA | Schaefer | -0.0014 (-0.0058, 0.00041) | 0.0037 (-0.0023, 0.0083) | 9e-4 (-0.0027, 0.0056) | -0.0016 (-0.0043, 0.00065) |
| | | DK | -0.0033 (-0.0095, 0.0028) | 0.0031 (-0.012, 0.014) | -0.014 (-0.029, 0.0021) | -0.00038 (-0.0048, 0.0047) |
| | CV | Schaefer | 0.00054 (-0.0034, 0.0042) | 0.0020 (-0.010, 0.018) | -0.0051 (-0.019, 0.0028) | -0.0018 (-0.0084, 0.0038) |
| | | DK | 0.0011 (-0.0058, 0.0083) | -7e-5 (-0.015, 0.018) | **-0.016 (-0.037, -0.00076)** | 0.00058 (-0.0069, 0.0068) |
| | SV | Aseg | -0.00086 (-0.013, 0.010) | -0.011 (-0.054, 0.018) | -0.023 (-0.063, 0.022) | **0.022 (0.0069, 0.050)** |
| BOLD | ALFF | Schaefer+Aseg | 0.0051 (-7e-4, 0.011) | **-0.0098 (-0.021, -0.00016)** | 0.0053 (-0.0045, 0.015) | -0.0020 (-0.0075, 0.0011) |
| | | DK+Aseg | 0.0076 (-0.0015, 0.017) | -0.013 (-0.029, 0.0033) | 0.0030 (-0.0095, 0.016) | -0.00083 (-0.0043, 0.0027) |
| | fALFF | Schaefer+Aseg | -0.00069 (-0.0067, 0.0058) | **-0.013 (-0.029, -0.0020)** | 0.0019 (-0.0045, 0.010) | 0.00084 (-0.0024, 0.0041) |
| | | DK+Aseg | 0.0038 (-0.0038, 0.015) | -0.0085 (-0.024, 0.0039) | **-0.014 (-0.023, -0.0091)** | 0.00049 (-0.0025, 0.0039) |
| | ReHo | Schaefer+Aseg | 0.0031 (-8e-4, 0.0074) | -0.0019 (-0.010, 0.0091) | 0.0021 (-0.0036, 0.0093) | 0.0015 (-0.00062, 0.0038) |
| | | DK+Aseg | **0.0084 (0.0017, 0.018)** | -0.00086 (-0.013, 0.011) | -0.0057 (-0.015, 0.0049) | -0.0010 (-0.0041, 0.0014) |
| | FC | Schaefer+Aseg | 0.0022 (-0.0020, 0.0068) | -0.0010 (-0.0061, 0.0040) | **0.0073 (0.0010, 0.015)** | 0.00076 (-0.0018, 0.0030) |
| | | DK+Aseg | 0.0019 (-0.0031, 0.0077) | -0.0027 (-0.0098, 0.0025) | 0.0037 (-0.0030, 0.012) | 5.0e-5 (-0.0047, 0.0035) |
| | vwFC | Schaefer+Aseg | 0.0049 (-4e-5, 0.011) | 0.0014 (-0.0062, 0.0090) | 0.0047 (-0.0026, 0.013) | -0.00012 (-0.0037, 0.0034) |
| | | DK+Aseg | 0.0040 (-0.0021, 0.013) | -0.00073 (-0.0096, 0.0074) | 0.0063 (-0.0027, 0.017) | -0.0012 (-0.0072, 0.0045) |
| DWI | FA | JHU-Tract | 0.0011 (-0.0026, 0.0049) | -0.0040 (-0.015, 0.0071) | -0.0036 (-0.011, 0.0033) | / |
| | | JHU-Label | 0.0017 (-0.0025, 0.0077) | -0.0023 (-0.0080, 0.0033) | 0.0034 (-0.0031, 0.015) | / |
| | MD | JHU-Tract | 0.0046 (-0.0041, 0.018) | -0.013 (-0.033, 0.0080) | -0.00039 (-0.017, 0.010) | / |
| | | JHU-Label | 0.0031 (-0.0046, 0.011) | 5e-5 (-0.011, 0.012) | 0.0038 (-0.0052, 0.015) | / |
| | AD | JHU-Tract | 0.0022 (-0.0051, 0.011) | -0.019 (-0.039, 0.00059) | -0.0043 (-0.022, 0.0050) | / |
| | | JHU-Label | 0.0024 (-0.0051, 0.010) | -0.0028 (-0.013, 0.0083) | -0.00034 (-0.0084, 0.0098) | / |
| | RD | JHU-Tract | 0.0028 (-0.0023, 0.011) | -0.0091 (-0.022, 0.0027) | -0.0021 (-0.013, 0.0058) | / |
| | | JHU-Label | 0.0024 (-0.0030, 0.0086) | -0.00037 (-0.0083, 0.011) | **0.0076 (0.0011, 0.019)** | / |
| | SC | Schaefer+Aseg | 0.00152 (-0.0061, 0.0086) | 0.00059 (-0.0062, 0.015) | **-0.012 (-0.023, -0.0025)** | / |
| | | DK+Aseg | 0.0020 (-0.0097, 0.0097) | -0.0089 (-0.019, 0.0069) | **-0.025 (-0.036, -0.017)** | / |

[a] the ICC difference and its 95% CI using CAT12 for skull-stripping. Positive ICC difference means using CAT12 would improve the reliability of brain features. Numbers in bold means the difference was statistically significant.

Table S26. The validity changes caused by CAT12's skull stripping for all brain features across all datasets.

| Modality | Feature | Atlas | pNKI | eNKI | SALD |
|---|---|---|---|---|---|
| T1 | CT | Schaefer | -0.0048 (-0.015, 0.0041)[a] | **-0.026 (-0.034, -0.018)** | **-0.037 (-0.049, -0.027)** |
| | | DK | -0.0044 (-0.014, 0.0046) | **-0.027 (-0.035, -0.019)** | **-0.037 (-0.049, -0.027)** |
| | CA | Schaefer | 0.00036 (-0.0014, 0.0023) | **0.0013 (0.00055, 0.0022)** | **-0.0042 (-0.0069, -0.0022)** |
| | | DK | 0.0011 (-0.0016, 0.0042) | 0.00067 (-0.00036, 0.0019) | **-0.0035 (-0.0061, -0.0013)** |
| | CV | Schaefer | **-0.0051 (-0.0088, -0.0022)** | **-0.0077 (-0.011, -0.0048)** | **-0.016 (-0.021, -0.011)** |
| | | DK | **-0.0048 (-0.0093, -0.0015)** | **-0.0089 (-0.012, -0.0061)** | **-0.016 (-0.021, -0.011)** |
| | SV | Aseg | 0.00059 (-0.011, 0.012) | **-0.0082 (-0.013, -0.0026)** | -0.00061 (-0.0085, 0.0063) |
| BOLD | ALFF | Schaefer+Aseg | -0.00072 (-0.0023, 0.00047) | **-0.0023 (-0.0047, -0.0013)** | **-0.0043 (-0.0059, -0.0028)** |
| | | DK+Aseg | -1.0e-4 (-0.0016, 0.0013) | **-0.0020 (-0.0041, -0.0010)** | **-0.0031 (-0.0046, -0.0016)** |
| | fALFF | Schaefer+Aseg | -0.00033 (-0.0016, 0.00081) | -0.00067 (-0.0021, 0.00019) | -0.00063 (-0.0015, 0.00013) |
| | | DK+Aseg | -5.0e-4 (-0.0021, 0.00081) | **-0.00094 (-0.0022, -1.0e-4)** | **-0.0012 (-0.0021, -0.00038)** |
| | ReHo | Schaefer+Aseg | 8.0e-5 (-0.00094, 0.0012) | -0.00015 (-0.00097, 0.00049) | 0.00013 (-0.00032, 0.00063) |
| | | DK+Aseg | -0.00093 (-0.0022, 0.00019) | -1.0e-5 (-0.0012, 7.0e-4) | **0.00062 (2.0e-5, 0.0013)** |
| | FC | Schaefer+Aseg | -8.0e-5 (-0.001, 0.0022) | -0.00013 (-0.00039, 0.00016) | 0.00013 (-0.00019, 0.00075) |
| | | DK+Aseg | 0.00017 (-0.00092, 0.0018) | -0.00012 (-0.00052, 0.00023) | 0.00014 (-0.00024, 0.00056) |
| | vwFC | Schaefer+Aseg | -0.00018 (-0.0014, 0.00063) | 2.0e-5 (-0.0010, 0.00097) | -0.00023 (-0.00061, 3.0e-5) |
| | | DK+Aseg | 0.00013 (-0.0010, 0.0014) | -4.0e-5 (-0.0012, 9.0e-4) | -0.00028 (-0.00095, 7.0e-5) |
| DWI | FA | JHU-Tract | 0.00022 (-0.0015, 0.0020) | 7.0e-5 (-6.0e-5, 0.00027) | / |
| | | JHU-Label | 0.0013 (-0.0012, 0.0094) | 3.0e-5 (-9.0e-5, 2.0e-4) | / |
| | MD | JHU-Tract | -0.00043 (-0.0022, 0.0012) | -5.0e-5 (-3.0e-4, 0.00019) | / |
| | | JHU-Label | 0.00042 (-8.0e-4, 0.0045) | -1.0e-4 (-0.00028, 5.0e-5) | / |
| | AD | JHU-Tract | -0.00044 (-0.0036, 0.00085) | -4.0e-5 (-0.00029, 2.0e-4) | / |
| | | JHU-Label | 0.0017 (-0.00066, 0.011) | -2.0e-5 (-0.00022, 2.0e-4) | / |
| | RD | JHU-Tract | -0.00026 (-0.0022, 0.0015) | -3.0e-5 (-0.00026, 0.00016) | / |
| | | JHU-Label | 0.00026 (-0.0011, 0.003) | -1.0e-5 (-0.00016, 0.00013) | / |
| | SC | Schaefer+Aseg | -0.00068 (-0.0019, 5.0e-4) | 0.00034 (-0.00066, 0.0018) | / |
| | | DK+Aseg | -0.0016 (-0.0033, 0) | 0.00015 (-0.00085, 0.0017) | / |

[a] the Age-R-Squared difference and its 95% CI using CAT12 for skull-stripping. Positive Age-R-Squared difference means using CAT12 would improve the validity of brain features. Numbers in bold means the difference was statistically significant.

Table S27. The reliability changes caused by processing variants for resting-state BOLD brain features across all datasets.

| Variant | Feature | Atlas | BNU1 | IPCAS1 | IPCAS2 | HNU1 |
|---|---|---|---|---|---|---|
| NOSTC | ALFF | Schaefer+Aseg | **0.0033 (0.00021, 0.0070)**[a] | 0.0051 (-0.00062, 0.012) | 0.0028 (-0.00073, 0.0071) | **0.0022 (0.00027, 0.0041)** |
| | | DK+Aseg | **0.0044 (0.00078, 0.0082)** | **0.0078 (0.0019, 0.016)** | 0.0032 (-0.0014, 0.0092) | 0.0019 (-0.00018, 0.0037) |
| | fALFF | Schaefer+Aseg | **0.021 (0.013, 0.032)** | -0.00069 (-0.015, 0.0091) | **0.031 (0.016, 0.059)** | **0.023 (0.017, 0.030)** |
| | | DK+Aseg | **0.019 (0.010, 0.031)** | **-0.020 (-0.034, -0.012)** | **0.020 (0.00032, 0.049)** | **0.021 (0.016, 0.028)** |
| | ReHo | Schaefer+Aseg | -0.0034 (-0.0077, 0.00058) | -0.0049 (-0.011, 0.0022) | **-0.014 (-0.023, -0.0055)** | **-0.0063 (-0.0093, -0.0044)** |
| | | DK+Aseg | -0.0041 (-0.0095, 0.0016) | -0.0054 (-0.018, 0.0029) | **-0.012 (-0.024, -0.0024)** | **-0.0071 (-0.0096, -0.0054)** |
| | FC | Schaefer+Aseg | -0.0035 (-0.012, 0.0026) | **0.0078 (0.00078, 0.021)** | -0.00073 (-0.012, 0.011) | 0.0011 (-0.0013, 0.0046) |
| | | DK+Aseg | -0.0034 (-0.012, 0.0032) | **0.012 (0.0040, 0.026)** | -0.00066 (-0.013, 0.011) | -0.00019 (-0.0033, 0.0044) |
| | vwFC | Schaefer+Aseg | -0.0027 (-0.011, 0.0034) | 0.0068 (-0.0010, 0.020) | 0.0029 (-0.0092, 0.019) | 0.00082 (-0.0021, 0.0049) |
| | | DK+Aseg | -0.0051 (-0.015, 0.0022) | **0.011 (0.0011, 0.027)** | 0.0034 (-0.011, 0.020) | -0.00023 (-0.0043, 0.0052) |
| NOLP | ReHo | Schaefer+Aseg | **0.080 (0.056, 0.11)** | **0.063 (0.024, 0.099)** | **0.036 (0.019, 0.053)** | **0.059 (0.050, 0.070)** |
| | | DK+Aseg | **0.083 (0.057, 0.11)** | **0.060 (0.020, 0.10)** | **0.027 (0.0071, 0.046)** | **0.060 (0.044, 0.076)** |
| | FC | Schaefer+Aseg | **0.089 (0.047, 0.15)** | **0.049 (0.0077, 0.098)** | **0.069 (0.029, 0.12)** | **0.091 (0.067, 0.13)** |
| | | DK+Aseg | **0.080 (0.040, 0.13)** | 0.039 (-0.0063, 0.098) | **0.065 (0.025, 0.12)** | **0.095 (0.070, 0.14)** |
| | vwFC | Schaefer+Aseg | **0.11 (0.030, 0.22)** | **0.083 (0.021, 0.16)** | **0.069 (0.0024, 0.14)** | **0.096 (0.054, 0.14)** |
| | | DK+Aseg | **0.10 (0.020, 0.22)** | 0.075 (-0.0078, 0.17) | 0.068 (-0.0052, 0.15) | **0.10 (0.053, 0.15)** |
| GSR | ALFF | Schaefer+Aseg | **-0.013 (-0.025, -0.0017)** | -0.00026 (-0.017, 0.023) | **0.0082 (0.00041, 0.021)** | 0.0042 (-0.0074, 0.017) |
| | | DK+Aseg | -0.0080 (-0.019, 0.0029) | -0.0019 (-0.018, 0.020) | 0.0035 (-0.0052, 0.016) | -0.00052 (-0.0097, 0.011) |
| | fALFF | Schaefer+Aseg | 0.0058 (-0.014, 0.030) | 0.021 (-0.0015, 0.044) | 0.011 (-0.011, 0.030) | **0.022 (0.010, 0.034)** |
| | | DK+Aseg | 0.0050 (-0.016, 0.031) | 0.0083 (-0.013, 0.029) | 0.013 (-0.0071, 0.031) | 0.0094 (-0.00032, 0.019) |
| | ReHo | Schaefer+Aseg | 0.016 (-0.0071, 0.042) | 0.0099 (-0.025, 0.050) | **0.027 (0.00076, 0.056)** | **0.044 (0.033, 0.057)** |
| | | DK+Aseg | 0.012 (-0.013, 0.039) | 0.016 (-0.033, 0.060) | 0.024 (-0.0031, 0.054) | **0.035 (0.025, 0.048)** |
| | FC | Schaefer+Aseg | -0.030 (-0.093, 0.014) | 0.012 (-0.021, 0.048) | -0.033 (-0.10, 0.012) | **-0.035 (-0.071, -0.0044)** |
| | | DK+Aseg | **-0.050 (-0.11, -0.0091)** | 0.026 (-0.019, 0.075) | **-0.040 (-0.10, -0.0030)** | **-0.044 (-0.088, -0.0094)** |
| | vwFC | Schaefer+Aseg | -0.012 (-0.081, 0.065) | 0.027 (-0.015, 0.080) | -0.040 (-0.13, 0.033) | -0.019 (-0.077, 0.034) |
| | | DK+Aseg | -0.039 (-0.12, 0.043) | 0.046 (-0.0018, 0.11) | -0.053 (-0.14, 0.013) | -0.038 (-0.11, 0.024) |
| MNI | ALFF | Schaefer+Aseg | **0.068 (0.051, 0.087)** | **0.062 (0.037, 0.081)** | **0.064 (0.038, 0.090)** | **0.040 (0.024, 0.058)** |
| | | DK+Aseg | **0.061 (0.044, 0.082)** | **0.056 (0.021, 0.087)** | **0.042 (0.014, 0.071)** | 0.017 (-0.0062, 0.033) |
| | fALFF | Schaefer+Aseg | 0.0050 (-0.025, 0.030) | 0.016 (-0.0060, 0.038) | 0.014 (-0.010, 0.040) | **-0.012 (-0.024, -0.0038)** |
| | | DK+Aseg | 0.011 (-0.019, 0.042) | 0.0068 (-0.023, 0.026) | -0.014 (-0.038, 0.0083) | -0.0037 (-0.021, 0.014) |
| | ReHo | Schaefer+Aseg | 0.0051 (-0.0058, 0.019) | 0.0023 (-0.025, 0.023) | -0.0067 (-0.027, 0.013) | -0.0018 (-0.014, 0.0081) |
| | | DK+Aseg | **0.019 (0.0049, 0.034)** | 0.011 (-0.014, 0.028) | 0.0078 (-0.015, 0.032) | 0.00032 (-0.016, 0.017) |
| | FC | Schaefer+Aseg | **0.0094 (0.0025, 0.018)** | **0.0079 (0.0013, 0.020)** | **0.017 (0.0077, 0.033)** | **0.012 (0.0082, 0.017)** |
| | | DK+Aseg | 0.0075 (-0.0036, 0.019) | 0.0016 (-0.018, 0.017) | 0.0029 (-0.0092, 0.016) | **0.012 (0.0038, 0.023)** |
| | vwFC | Schaefer+Aseg | **0.013 (0.0058, 0.022)** | 0.00031 (-0.012, 0.014) | 0.0022 (-0.013, 0.018) | **0.010 (0.0052, 0.017)** |
| | | DK+Aseg | 0.015 (-0.0021, 0.030) | -0.0073 (-0.022, 0.0065) | -0.010 (-0.031, 0.0033) | 0.0073 (-0.0062, 0.020) |

[a] the ICC difference and its 95% CI using different processing variants. Positive ICC difference means the processing variant would improve the reliability of brain features. Numbers in bold means the difference was statistically significant.

Table S28. The validity changes caused by processing variants for resting-state BOLD brain features across all datasets.

| Variant | Feature | Atlas | pNKI | eNKI | SALD |
|---|---|---|---|---|---|
| NOSTC | ALFF | Schaefer+Aseg | **0.0022 (0.00029, 0.0037)**[a] | **0.00091 (0.00016, 0.0017)** | **0.0035 (0.0022, 0.0050)** |
| | | DK+Aseg | **0.0021 (0.00049, 0.0038)** | **0.00089 (0.00021, 0.0016)** | **0.0029 (0.0018, 0.0042)** |
| | fALFF | Schaefer+Aseg | 0.0042 (-0.0040, 0.013) | **0.0070 (0.0043, 0.010)** | **0.0045 (0.0023, 0.0067)** |
| | | DK+Aseg | 0.0042 (-0.0057, 0.014) | **0.0054 (0.0028, 0.0085)** | **0.0042 (0.0016, 0.0068)** |
| | ReHo | Schaefer+Aseg | 0.0019 (-0.0026, 0.0060) | 0.00034 (-0.00097, 0.0018) | 0.00059 (-0.00021, 0.0014) |
| | | DK+Aseg | 0.0015 (-0.0025, 0.0054) | 0.00026 (-0.0011, 0.0018) | 0.00019 (-0.00077, 0.0012) |
| | FC | Schaefer+Aseg | 4.0e-5 (-0.0028, 0.0022) | 0.00044 (-0.00076, 0.0017) | -0.00045 (-0.0016, 0.00032) |
| | | DK+Aseg | -0.00061 (-0.0038, 0.0021) | 0.00078 (-0.00045, 0.0022) | -3.0e-5 (-0.0011, 0.00065) |
| | vwFC | Schaefer+Aseg | 4.0e-5 (-0.0017, 0.0021) | **0.0022 (0.00073, 0.0047)** | 0.00058 (-0.00025, 0.0016) |
| | | DK+Aseg | -0.00045 (-0.0036, 0.0017) | **0.0021 (0.00047, 0.0045)** | 0.00062 (-9.0e-5, 0.0018) |
| NOLP | ReHo | Schaefer+Aseg | **0.026 (0.017, 0.037)** | **0.028 (0.023, 0.034)** | **0.018 (0.013, 0.023)** |
| | | DK+Aseg | **0.022 (0.014, 0.033)** | **0.026 (0.021, 0.032)** | **0.016 (0.011, 0.021)** |
| | FC | Schaefer+Aseg | -0.0012 (-0.015, 0.013) | **0.0065 (0.0011, 0.013)** | 0.0010 (-0.0065, 0.0091) |
| | | DK+Aseg | -0.00077 (-0.016, 0.015) | **0.0081 (0.0026, 0.015)** | 0.0050 (-0.0031, 0.014) |
| | vwFC | Schaefer+Aseg | 0.0091 (-0.0074, 0.030) | **0.028 (0.018, 0.040)** | **0.012 (0.0029, 0.024)** |
| | | DK+Aseg | 0.0061 (-0.012, 0.027) | **0.025 (0.016, 0.037)** | **0.013 (0.0039, 0.026)** |
| GSR | ALFF | Schaefer+Aseg | -0.0019 (-0.0059, 0.0022) | **-0.0036 (-0.0065, -0.00088)** | **0.0032 (0.00063, 0.0055)** |
| | | DK+Aseg | -0.0015 (-0.0047, 0.0025) | **-0.0034 (-0.0058, -0.0012)** | 0.0013 (-0.00091, 0.0034) |
| | fALFF | Schaefer+Aseg | **0.0069 (0.0023, 0.013)** | **0.0058 (0.0018, 0.0099)** | **0.0064 (0.0036, 0.0096)** |
| | | DK+Aseg | **0.0054 (0.0011, 0.011)** | **0.0047 (0.0014, 0.0081)** | **0.0049 (0.0022, 0.0077)** |
| | ReHo | Schaefer+Aseg | 0.0084 (-0.0014, 0.020) | 0.0032 (-0.0022, 0.0086) | 0.00036 (-0.0038, 0.0041) |
| | | DK+Aseg | 0.0045 (-0.0044, 0.015) | 0.0031 (-0.0017, 0.0077) | -0.00069 (-0.0050, 0.0030) |
| | FC | Schaefer+Aseg | 0.0013 (-0.032, 0.013) | 0.0012 (-0.0020, 0.0025) | -0.00022 (-0.0079, 0.0029) |
| | | DK+Aseg | 0.0052 (-0.025, 0.017) | 0.0018 (-0.0022, 0.0034) | 0.0023 (-0.0025, 0.0044) |
| | vwFC | Schaefer+Aseg | **0.019 (0.0044, 0.030)** | -0.0075 (-0.022, 0.0015) | 0.0048 (-0.0015, 0.0080) |
| | | DK+Aseg | 0.016 (-0.00013, 0.028) | -0.0080 (-0.024, 0.00065) | 0.0045 (-0.0045, 0.0086) |
| MNI | ALFF | Schaefer+Aseg | -0.0055 (-0.023, 0.010) | -0.00078 (-0.0070, 0.0056) | -0.00057 (-0.0074, 0.0062) |
| | | DK+Aseg | **-0.019 (-0.034, -0.0064)** | **-0.0084 (-0.014, -0.0035)** | **-0.014 (-0.021, -0.0079)** |
| | fALFF | Schaefer+Aseg | **-0.0098 (-0.022, -0.00091)** | **-0.015 (-0.020, -0.011)** | **-0.0078 (-0.012, -0.0042)** |
| | | DK+Aseg | **-0.012 (-0.028, -0.0018)** | **-0.016 (-0.022, -0.012)** | **-0.0097 (-0.014, -0.0055)** |
| | ReHo | Schaefer+Aseg | **-0.017 (-0.027, -0.011)** | **-0.011 (-0.014, -0.0088)** | **-0.0025 (-0.0046, -0.00067)** |
| | | DK+Aseg | **-0.023 (-0.036, -0.015)** | **-0.0096 (-0.013, -0.0070)** | **-0.0024 (-0.0049, -6.0e-5)** |
| | FC | Schaefer+Aseg | 0.0022 (-0.00075, 0.0065) | 0.00065 (-0.00074, 0.0023) | **0.0034 (0.0014, 0.0061)** |
| | | DK+Aseg | 0.0034 (-0.00029, 0.0083) | -0.00098 (-0.0025, 0.00059) | 0.00061 (-0.0012, 0.0029) |
| | vwFC | Schaefer+Aseg | -0.00044 (-0.0098, 0.012) | **-0.0082 (-0.012, -0.0047)** | 9.0e-5 (-0.0023, 0.0032) |
| | | DK+Aseg | 0.0041 (-0.0096, 0.018) | **-0.0085 (-0.013, -0.0041)** | -0.00054 (-0.0034, 0.0022) |

[a] the Age-R-Squared difference and its 95% CI using different processing variants. Positive Age-R-Squared difference means the processing variant would improve the validity of brain features. Numbers in bold means the difference was statistically significant.

Table S29. The reliability difference between PhiPipe and DPARSF for resting-state BOLD brain features, and between PhiPipe and PANDA for DWI brain features across all datasets.

| Modality | Feature | Atlas | BNU1 | IPCAS1 | IPCAS2 | HNU1 |
|---|---|---|---|---|---|---|
| BOLD | ALFF | Schaefer+Aseg | **0.046 (0.0093, 0.085)**[a] | -0.027 (-0.011, 0.027) | **0.030 (0.001, 0.063)** | -0.023 (-0.050, 0.0042) |
| | fALFF | Schaefer+Aseg | **0.056 (0.013, 0.096)** | 0.021 (-0.053, 0.086) | -0.021 (-0.082, 0.034) | 0.014 (-0.019, 0.041) |
| | ReHo | Schaefer+Aseg | -0.011 (-0.045, 0.020) | -0.0061 (-0.046, 0.033) | 0.0049 (-0.044, 0.043) | 0.023 (-0.0071, 0.055) |
| | FC | Schaefer+Aseg | **0.077 (0.029, 0.13)** | -0.0012 (-0.056, 0.058) | **0.11 (0.069, 0.17)** | **0.048 (0.025, 0.076)** |
| | vwFC | Schaefer+Aseg | **0.11 (0.048, 0.19)** | -0.00039 (-0.061, 0.044) | 0.087 (-0.0051, 0.16) | **0.059 (0.026, 0.11)** |
| DWI | FA | JHU-Tract | **-0.0092 (-0.023, 0.0034)** | -0.039 (-0.13, 0.016) | **-0.053 (-0.11, -0.014)** | / |
| | MD | JHU-Tract | **0.062 (0.038, 0.090)** | -0.016 (-0.078, 0.027) | 0.029 (-0.0026, 0.057) | / |
| | AD | JHU-Tract | 0.011 (-0.018, 0.036) | **-0.076 (-0.13, -0.034)** | 0.014 (-0.024, 0.042) | / |
| | RD | JHU-Tract | **0.028 (0.012, 0.044)** | -0.015 (-0.076, 0.018) | -0.014 (-0.048, 0.012) | / |
| | SC | Schaefer+Aseg | **-0.14 (-0.16, -0.13)** | **-0.059 (-0.080, -0.037)** | **-0.12 (-0.15, -0.10)** | / |

[a] the ICC difference and its 95% CI between PhiPipe and DPARSF/PANDA. Positive ICC difference means the DPARSF/PANDA had better reliability of brain features. Numbers in bold means the difference was statistically significant.

Table S30. The validity difference between PhiPipe and DPARSF for resting-state BOLD brain features, and between PhiPipe and PANDA for DWI brain features across all datasets.

| Modality | Feature | Atlas | pNKI | eNKI | SALD |
|---|---|---|---|---|---|
| BOLD | ALFF | Schaefer+Aseg | **-0.031 (-0.054, -0.013)**[a] | **-0.012 (-0.021, -0.0055)** | **-0.032 (-0.043, -0.021)** |
| | fALFF | Schaefer+Aseg | -0.00063 (-0.022, 0.021) | **-0.011 (-0.020, -0.0018)** | 0.0054 (-0.0042, 0.015) |
| | ReHo | Schaefer+Aseg | -0.0068 (-0.019, 0.0047) | **-0.012 (-0.018, -0.0065)** | 0.0032 (-0.0025, 0.0094) |
| | FC | Schaefer+Aseg | -0.0075 (-0.035, 0.0049) | **0.0019 (1.0e-4, 0.0049)** | 0.00095 (-0.0047, 0.0041) |
| | vwFC | Schaefer+Aseg | 0.0066 (-0.010, 0.037) | 0.00015 (-0.0082, 0.0095) | **0.011 (0.0049, 0.022)** |
| DWI | FA | JHU-Tract | 0.0028 (-0.023, 0.032) | **0.0037 (0.0017, 0.0071)** | / |
| | MD | JHU-Tract | **0.024 (0.0079, 0.045)** | **-0.0041 (-0.0070, -0.00041)** | / |
| | AD | JHU-Tract | -0.0035 (-0.020, 0.019) | -0.0046 (-0.0084, 0.00045) | / |
| | RD | JHU-Tract | 0.010 (-0.0060, 0.027) | 0.00027 (-0.0015, 0.0028) | / |
| | SC | Schaefer+Aseg | **-0.0097 (-0.015, -0.0051)** | **-0.0081 (-0.011, -0.0056)** | / |

[a] the Age-R-Squared difference and its 95% CI between PhiPipe and DPARSF/PANDA. Positive Age-R-Squared difference means the DPARSF/PANDA had better validity of brain features. Numbers in bold means the difference was statistically significant.

Table S31. The multivariate reliability for all brain features across all datasets

| Modality | Feature | Atlas | BNU1 | IPCAS1 | IPCAS2 | HNU1 |
|---|---|---|---|---|---|---|
| T1 | CT | Schaefer | 0.85 (0.81, 0.88)[a] | 0.81 (0.75, 0.87) | 0.86 (0.83, 0.89) | 0.77 (0.74, 0.80) |
| | | DK | 0.79 (0.76, 0.83) | 0.76 (0.68, 0.83) | 0.75 (0.68, 0.81) | 0.67 (0.62, 0.72) |
| | CA | Schaefer | 0.98 (0.98, 0.98) | 0.96 (0.95, 0.97) | 0.97 (0.96, 0.98) | 0.96 (0.95, 0.97) |
| | | DK | 0.98 (0.97, 0.98) | 0.96 (0.95, 0.97) | 0.97 (0.95, 0.98) | 0.97 (0.96, 0.98) |
| | CV | Schaefer | 0.96 (0.95, 0.97) | 0.94 (0.92, 0.96) | 0.95 (0.93, 0.96) | 0.93 (0.92, 0.95) |
| | | DK | 0.96 (0.94, 0.97) | 0.95 (0.93, 0.97) | 0.96 (0.95, 0.97) | 0.95 (0.94, 0.96) |
| | SV | Aseg | 0.95 (0.94, 0.96) | 0.94 (0.86, 0.97) | 0.91 (0.85, 0.95) | 0.88 (0.80, 0.93) |
| BOLD | ALFF | Schaefer+Aseg | 0.53 (0.43, 0.63) | 0.61 (0.52, 0.68) | 0.57 (0.48, 0.67) | 0.64 (0.58, 0.68) |
| | | DK+Aseg | 0.52 (0.41, 0.61) | 0.56 (0.46, 0.66) | 0.55 (0.44, 0.65) | 0.60 (0.55, 0.64) |
| | fALFF | Schaefer+Aseg | 0.43 (0.34, 0.51) | 0.46 (0.35, 0.56) | 0.40 (0.28, 0.51) | 0.48 (0.41, 0.54) |
| | | DK+Aseg | 0.38 (0.27, 0.48) | 0.45 (0.29, 0.58) | 0.34 (0.21, 0.47) | 0.46 (0.38, 0.52) |
| | ReHo | Schaefer+Aseg | 0.48 (0.37, 0.57) | 0.47 (0.39, 0.54) | 0.49 (0.36, 0.60) | 0.56 (0.51, 0.61) |
| | | DK+Aseg | 0.46 (0.33, 0.58) | 0.50 (0.39, 0.60) | 0.46 (0.32, 0.58) | 0.58 (0.52, 0.63) |
| | FC | Schaefer+Aseg | 0.28 (0.22, 0.35) | 0.24 (0.17, 0.31) | 0.29 (0.21, 0.37) | 0.37 (0.32, 0.41) |
| | | DK+Aseg | 0.27 (0.20, 0.34) | 0.21 (0.12, 0.29) | 0.26 (0.18, 0.34) | 0.35 (0.30, 0.40) |
| | vwFC | Schaefer+Aseg | 0.29 (0.20, 0.39) | 0.26 (0.19, 0.32) | 0.35 (0.24, 0.47) | 0.35 (0.28, 0.43) |
| | | DK+Aseg | 0.29 (0.19, 0.40) | 0.22 (0.14, 0.30) | 0.31 (0.21, 0.41) | 0.35 (0.27, 0.43) |
| DWI | FA | JHU-tract | 0.92 (0.90, 0.93) | 0.90 (0.88, 0.93) | 0.88 (0.84, 0.91) | / |
| | | JHU-Label | 0.87 (0.85, 0.89) | 0.84 (0.80, 0.87) | 0.83 (0.78, 0.86) | / |
| | MD | JHU-Tract | 0.84 (0.80, 0.87) | 0.84 (0.76, 0.88) | 0.83 (0.75, 0.88) | / |
| | | JHU-Label | 0.85 (0.82, 0.88) | 0.82 (0.75, 0.87) | 0.83 (0.74, 0.88) | / |
| | AD | JHU-Tract | 0.86 (0.83, 0.88) | 0.83 (0.80, 0.86) | 0.85 (0.78, 0.89) | / |
| | | JHU-Label | 0.82 (0.79, 0.85) | 0.82 (0.76, 0.86) | 0.82 (0.76, 0.87) | / |
| | RD | JHU-Tract | 0.88 (0.85, 0.90) | 0.88 (0.84, 0.91) | 0.85 (0.79, 0.88) | / |
| | | JHU-Label | 0.87 (0.84, 0.90) | 0.84 (0.78, 0.88) | 0.84 (0.75, 0.89) | / |
| | SC | Schaefer+Aseg | 0.74 (0.71, 0.76) | 0.74 (0.73, 0.76) | 0.67 (0.63, 0.72) | / |
| | | DK+Aseg | 0.65 (0.61, 0.69) | 0.63 (0.60, 0.66) | 0.60 (0.55, 0.65) | / |

[a] dbICC and its 95% confidence interval

Table S32. The multivariate predicative validity for all brain features across all datasets

| Modality | Feature | Atlas | pNKI | eNKI | SALD |
|---|---|---|---|---|---|
| T1 | CT | Schaefer | 0.74/0.61[a] | 0.84/0.67 | 0.78/0.66 |
| | | DK | 0.73/0.59 | 0.82/0.66 | 0.74/0.66 |
| | CA | Schaefer | 0.38/0.28 | 0.56/0.31 | 0.62/0.47 |
| | | DK | 0.059/0.26 | 0.58/0.31 | 0.57/0.46 |
| | CV | Schaefer | 0.52/0.50 | 0.76/0.54 | 0.78/0.66 |
| | | DK | 0.43/0.48 | 0.74/0.53 | 0.73/0.66 |
| | SV | Aseg | 0.60/0.52 | 0.72/0.62 | 0.75/0.64 |
| BOLD | ALFF | Schaefer+Aseg | 0.56/0.60 | 0.66/0.56 | 0.75/0.65 |
| | | DK+Aseg | 0.61/0.61 | 0.64/0.55 | 0.73/0.62 |
| | fALFF | Schaefer+Aseg | 0.55/0.60 | 0.62/0.51 | 0.62/0.50 |
| | | DK+Aseg | 0.66/0.62 | 0.62/0.54 | 0.61/0.50 |
| | ReHo | Schaefer+Aseg | 0.75/0.75 | 0.73/0.62 | 0.68/0.51 |
| | | DK+Aseg | 0.78/0.70 | 0.70/0.59 | 0.68/0.56 |
| | FC | Schaefer+Aseg | 0.68/0.57 | 0.70/0.59 | 0.83/0.66 |
| | | DK+Aseg | 0.65/0.49 | 0.65/0.54 | 0.74/0.63 |
| | vwFC | Schaefer+Aseg | 0.61/0.58 | 0.70/0.46 | 0.79/0.57 |
| | | DK+Aseg | 0.56/0.53 | 0.66/0.43 | 0.71/0.52 |
| DWI | FA | JHU-Tract | 0.61/0.58 | 0.68/0.20 | / |
| | | JHU-Label | 0.65/0.64 | 0.80/0.75 | / |
| | MD | JHU-Tract | 0.70/0.62 | 0.68/0.58 | / |
| | | JHU-Label | 0.67/0.63 | 0.74/0.64 | / |
| | AD | JHU-Tract | 0.71/0.68 | 0.65/0.57 | / |
| | | JHU-Label | 0.70/0.71 | 0.74/0.62 | / |
| | RD | JHU-Tract | 0.66/0.64 | 0.69/0.44 | / |
| | | JHU-Label | 0.63/0.65 | 0.77/0.65 | / |
| | SC | Schaefer+Aseg | 0.75/0.73 | 0.79/0.77 | / |
| | | DK+Aseg | 0.76/0.77 | 0.85/0.80 | / |

[a] CV-R using SVR/CPM models
